# Integrated single-cell and spatial multiomic analysis reveals widespread reactivation of developmental programs in diseased human hearts

**DOI:** 10.1101/2025.04.01.646460

**Authors:** William Gao, Peng Hu, Qi Qiu, Xiangjin Kang, Kenneth C. Bedi, Kotaro Sasaki, Kenneth B. Margulies, Hao Wu

**Author notes:** Correspondence should be addressed to W.G. and H.W. These authors contributed equally: W.G. and P.H.

## Abstract

The first organ to develop in utero, the human heart undergoes significant changes during development and must sustain its function over a lifetime. To better characterize molecular changes in human cardiac cell-types across sex, aging, developmental and disease, we analyzed single nucleus RNA sequencing (snRNA-seq) datasets from 299 donors, identifying many more differentially expressed genes (DEGs) across developmental and disease states than by sex and age. In cardiomyocytes and most non-cardiomyocyte cell types, developmental and disease DEGs showed significant overlap. Cardiac development and disease were associated with convergent changes in non-cardiomyocyte intercellular communication, including TGFβ signaling, but differences in cell-type proportions. By integrating snRNA-seq with 106 snATAC-seq datasets, we reveal potential transcriptional factors driving fetal reactivation in disease. Finally, using spatial transcriptomics data, we identify that fetal reactivation is highly localized in niches. This work offers the largest multimodal, cell-type resolved interrogation of the human heart, providing insights into convergence in development and disease.

## Main

Cardiovascular disease is the leading cause of death worldwide. Age is one of the greatest risk factors, with cardiovascular associated mortality growing exponentially after age 65 ^1^. Due to the significant burden of cardiovascular disease, several studies have explored the mammalian heart across development, aging, and disease states, including in human donor hearts. In contrast to initial studies using microarrays and bulk RNA-sequencing ^2–4^, single cell RNA-sequencing (scRNA-seq) enables cell-type resolved analysis ^5^. Since the first human heart single cell atlases were published in 2020 ^6,7^, the number of studies using human donor hearts has grown. A recent study performed a meta-analysis of single-nucleus RNA-seq from 73 non-diseased donors and snATAC-seq from 9 donors to identify sex and age-related transcriptomic and epigenomic changes in the human heart ^8^. However, whether these age-related genes relate to disease burden are unclear. No study has comprehensively compared transcriptional and epigenomic signatures of the human heart across early development, aging, and disease. By combining numerous published and newly generated datasets together to perform the largest such integrated single-cell multiomics analysis to date, we explored how the human heart changes across sex, aging, development, and disease.

Performing an integrated analysis offers distinct challenges, including prominent inter-study batch effects ^9^ and computational bottlenecks, as many analytical tools that do not scale to large datasets ^10^. However, an integrated analysis also offers several advantages. First, most individual studies included cohort sizes fewer than 50 donors, resulting in lower statistical power ^11^. Aggregating multiple datasets can discriminate replicable “ground truth” signals from study-specific artifacts. Second, as the single-cell field has grown, analytical methods have also advanced, allowing for richer utilization of these data. Here, by integrating 299 snRNA-seq, 106 snATAC-seq, and 27 spatial transcriptomics human donor datasets, we identify robust transcriptional changes across cardiac developmental, aging, and disease and place them in an epigenetic and spatial context.

## RESULTS

### Integration of ∼2.3 million left ventricular snRNA-seq nuclei across 299 donors

To study human cardiac maturation and aging in a cell-type specific manner, we initially generated a snRNA-seq dataset of 19 donors (**Fig. S1**), including 2 fetal, 15 non-diseased (ND) postnatal, and 2 diseased donors using the droplet microfluidics based sNucDrop-seq method^12^. To further increase the number of donors to gain statistical power, we then performed a literature search for all published human heart sn/scRNA-seq datasets (**Fig. S2**). Most studies performed snRNA-seq rather than scRNA-seq (**Fig. S3**) since cardiomyocytes do not fit within most droplet-based encapsulation protocols^5,12^ and sampled the left ventricle (LV). Therefore, we focused our analysis on LV snRNA-seq datasets, except for fetal samples, in which the whole heart was typically sampled. To reduce the number of batch effects, we prioritized datasets that satisfied one of these criteria: (1) at least 10 donors, (2) at least 3 donors across two disease + developmental categories. The disease + developmental categories were fetal ND, postnatal ND, and postnatal diseased. To this end, 10 studies with 280 donors satisfied these criteria^7,13–23^. Altogether, we analyzed 299 snRNA-seq LV datasets (**Fig. 1a**). Collectively, these datasets are diverse by sex (111 female, 188 male), age group (13 fetal, 57 young (≤ 40 yr), 128 middle (40-59 yr), 101 old (≥ 60 yr), and disease status (167 ND, 132 diseased) (**Fig. 1b, Table S2**). The diseased donors were diagnosed with many types of heart disease: arrhythmogenic cardiomyopathy (ARVC), dilated cardiomyopathy (DCM), hypertrophic cardiomyopathy (HCM), ischemic cardiomyopathy (ICM), and non-compaction cardiomyopathy (NCCM). However, only DCM was profiled in more than two studies (**Fig. S2a; Table S2**), so we binarized disease status, resulting in 167 ND and 132 diseased donors.

**Fig. 1:**
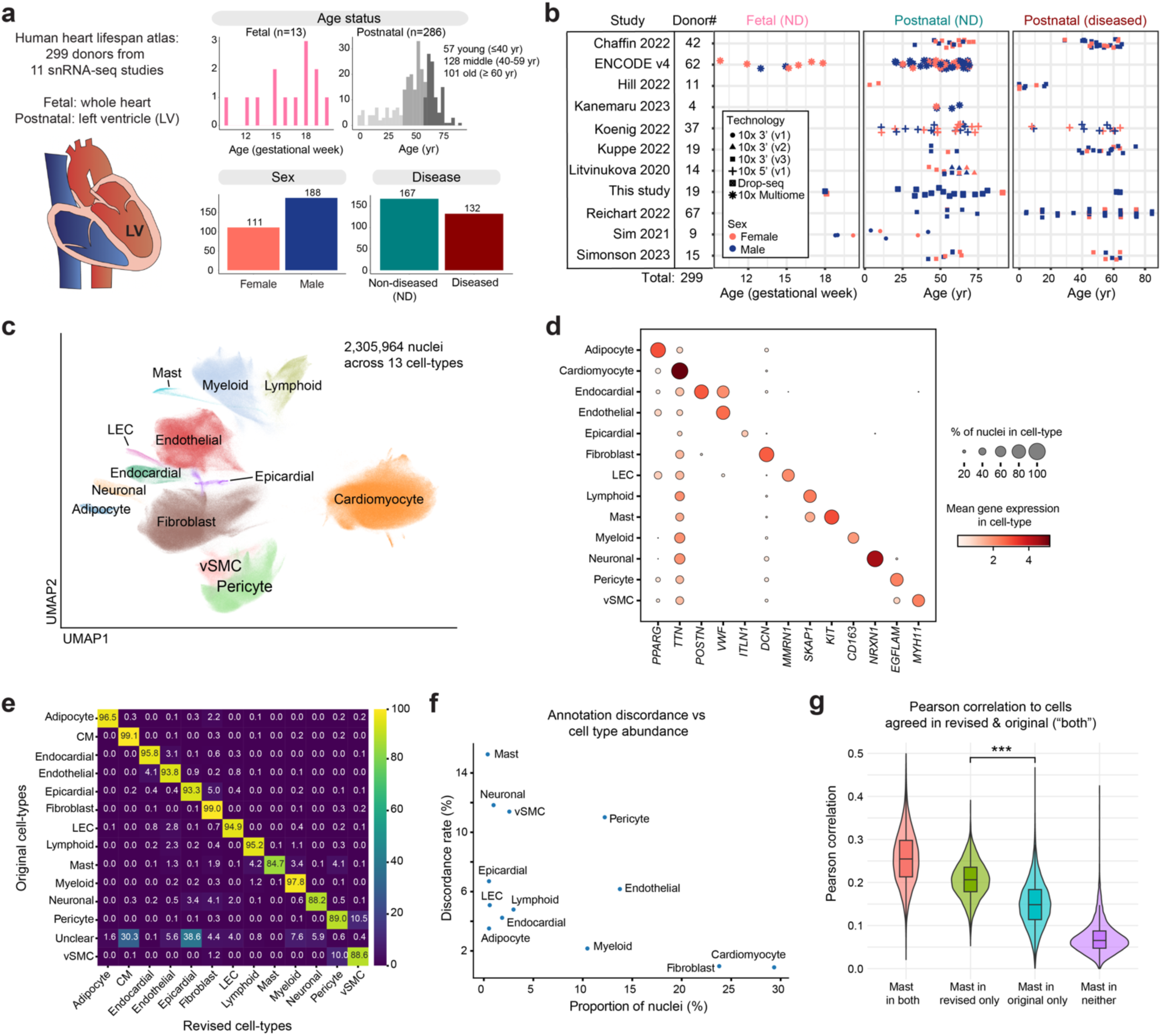
Overview of integrated analysis of single-nucleus RNA sequencing human cardiac datasets. **a)** Analysis involves hearts from 299 human donors, with diversity across age, sex and disease states. **b)** Stratification of donors by study, age + disease status, and snRNA-seq technology platform. **c)** Uniform manifold approximation and projection (UMAP) of 2,305,964 nuclei across 13 cell types. LEC, lymphatic epithelial cells; vSMC, vascular smooth muscle cells. **d)** Marker gene expression across cell type clusters. **e)** Heatmap showing concordance between original and revised cell type annotations. **f)** Annotation discordance rate versus cell type proportion. **g)** Transcriptome-wide Pearson correlation of Mast cells in both Chaffin 2022 and revised annotation (“both”) to other Mast cells in both annotations (“both”), Mast cells only in revised annotation (“revised only”), Mast cells only in Chaffin 2022 annotation (“original only”), and non-Mast cells according to both annotations.

### Integration of datasets refines cell type annotations for rarer cell types

As the first step in our integrated analysis, we combined the processed count matrices provided by each individual dataset and performed standard preprocessing for our newly generated dataset (**Methods**). After quality control filtering, we retained 2,305,964 nuclei (referred from hereon as the “aggregated dataset”). To remove batch effects in both the Uniform Approximation and Manifold Projection (UMAP) embedding and count matrix^1,2^, we used scVI^25^, which can scale to millions of cells^10^ (**Fig. 1c**). With this new embedding, we manually revised the cell type annotations using marker genes (**Fig. 1d**) independently from the original annotation, identifying 13 distinct cell types: adipocyte, cardiomyocyte, endocardial, endothelial, epicardial, fibroblast, lymphatic epithelial (LEC), lymphoid, mast, myeloid, neuronal, pericyte, and vascular smooth muscle (vSMC) cells. We observed strong concordance between our revised annotation and the original annotations provided by the published studies (**Fig. 1e**). Notably, there was a higher discordance between the revised annotation and original annotation for rarer cell types (**Fig. 1f**). We hypothesized that rarer cell types may not form distinct clusters in datasets with fewer sampled cells and could be erroneously annotated as a more common cell type. To test this, we computed the Pearson correlation of cells annotated as a given cell type only in our revised annotation (“revised only”) and cells labeled such only in the original annotation (“original only”) to “ground truth” cells (annotated in both revised + original annotation). We found that “revised only” cells generally had higher correlations to the “ground truth” cells than “original only” cells (**Fig. 1g**). Therefore, integration of multiple datasets enhances cell type annotation. Overall, this aggregated dataset can serve as a reference for annotation of future datasets and allows for more accurate downstream cell-type specific analysis.

### Sex and aging are associated with few differentially expressed genes

With these refined cell type annotations, we identified cell type resolved differentially expressed genes (DEGs) across distinct biological states. To perform this analysis in a cell-type specific manner, we pseudobulked all counts for the same cell type annotation and donor together (**Fig. 2a**). This pseudobulked approach substantially decreased compute time and the number of false positive DEGs compared to approaches in which single cells are treated as individual observation units, since expression profiles within the same donor are highly correlated^9,26^. We then used DESeq2^27,28^ to identify DEGs along 4 contrasts: sex (male vs. female); aging (old vs. young); developmental (fetal vs. young postnatal); disease (Y vs N), while regressing out technology and study, which was identified as the major batch effect (**Fig. S4**). Performing cell type resolved analysis allowed us to (1) compare DEGs across different cell types for the same contrast and (2) compare DEGs in the same cell type across different contrasts to examine their overlap.

**Fig. 2:**
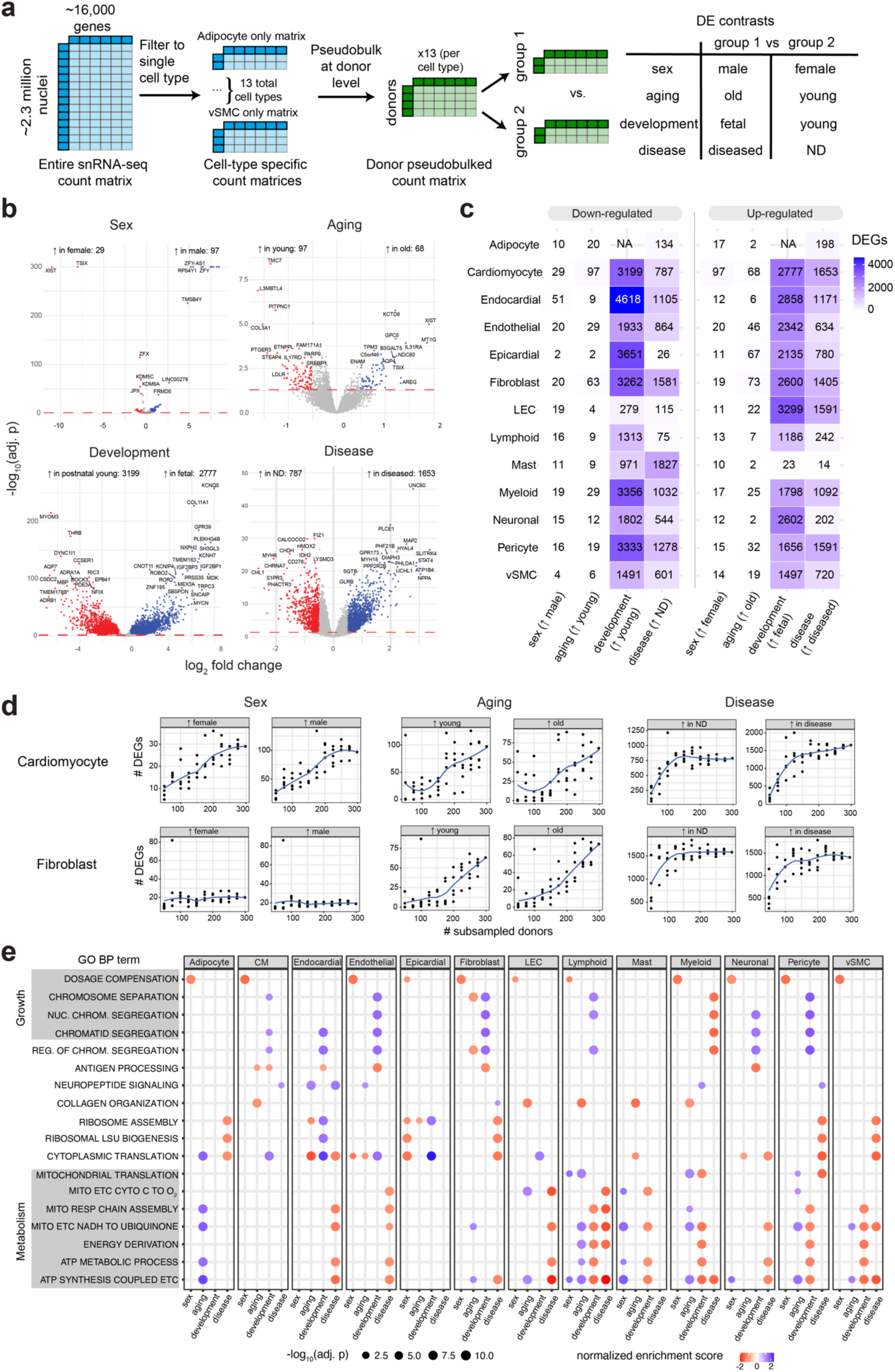
Cell-type resolved differential gene expression and gene set enrichment analysis. **a)** Pseudobulked profiles for each cell-type were obtained by combining counts for all cells belonging to the same cell type for each donor. This pseudobulked count matrix was used for differential gene expression analysis with DESeq2. **b)** The number of differentially expressed genes (DEGs) across developmental, aging, disease, and sex for “cardiomyocyte”. Volcano plots for all cardiac cell types are in Fig. S5-S17. **c)** Number of DEGs per cell type x contrast combination. There were insufficient fetal adipocytes for the adipocyte developmental contrast. **d)** Number of DEGs, across sex, aging, and disease contrasts, in cardiomyocytes and fibroblasts with subsampled number of donors. **e)** Gene set enrichment analysis across cell types and contrasts.

For sex-specific expression, we identified relatively few DEGs (**Fig. 2b-c; Fig. S5-S17; Tables S4-S17**). In most cell types, except for cardiomyocytes, fewer than 15 genes were differentially expressed between males and females. As expected, these sex-associated DEGs included genes located on the X chromosome (e.g. *XIST, TSIX, KDM6A*) and Y chromosome (*DDX3Y, KDM5D, LINC00278*) (**Fig. 2b**). Similarly, aging-related contrasts revealed a limited number of DEGs across all cell types (**Fig. 2c**). In cardiomyocytes, notable DEGs declining with age are *LDLR* (low-density lipoprotein receptor) and its regulator, *SREBF1*^29^ (**Fig. 2b**). Most age-related DEGs were cell-type specific (**Fig. S18**), underscoring the utility of snRNA-seq in resolving cell-type specific transcriptomic changes. However, some DEGs were shared across multiple cell types, including *PTCHD4*, which increased with age in seven human cardiac cell types and may play a role in senescence^4,30^. Notably, we did not observe a significant age-related increase in well-known senescence markers *CDKN1A* (p16) and *CDKN2A* (p21) or in a senescence score derived from the SenMayo gene set^31^ for any cell type (**Fig. S19**). This may reflect how aging and senescence are largely distinct molecular programs. Alternatively, genes such as *PTCHD4* may serve as more relevant markers for cardiac senescence, which differs across tissues^32^. Additionally, transcriptional noise did not exhibit significant age-related changes (**Fig. S20**). Together, these results suggest that healthy human cardiac aging is not associated with extensive transcriptional dysregulation, underscoring the resilience of the heart at the transcriptional level.

### Many genes are differentially expressed during developmental and disease

In contrast to sex and age, we identified thousands of DEGs during the fetal-postnatal transition (**Fig. 2b**). Several cell types display strong fetal-biased expression of insulin-like growth factor binding proteins such as *IGF2BP1* and *IGF2BP3*, which drive cardiac proliferation^33^. In contrast, multiple cell types in the young postnatal heart show increased expression of *THRB*, thyroid receptor B, which inhibits proliferation and drives perinatal cardiac maturation^34^. The mean number of cell types sharing a development DEG was also notably higher than for other contrasts, suggesting that many cell types respond similarly this transitional period (**Fig. S18**).

In the disease state, we also identify relatively large number of DEGs (spanning hundreds to thousands of genes) across most cell types (**Fig. 2c**). In cardiomyocytes, there is strong upregulation of known heart failure markers including atrial natriuretic peptide (*NPPA*)^35^ and decrease of *MYH6*^36^ (**Fig. 2b**). *ACE2*, which reduces blood pressure, inflammation, and adverse cardiac remodeling through the degradation of angiotensin II^1^ and serves as the human receptor for the SAR-CoV2 virus^38^, is specifically downregulated in cardiac fibroblasts (**Fig. S10d**). Additionally, diseased fibroblasts have decreased expression of endothelial receptor B, *EDNRB*, and monoamine oxidase, *MAOA*, which both regulate blood pressure. Surprisingly, in myeloid cells, the strongest downregulated genes are the pro-inflammatory alarmins *S100A8* and *S100A9* (**Fig. S14**). In endothelial cells, the downregulation of several members of the cytochrome P450 system, which perform drug metabolism and lipid synthesis, may underlie endothelial dysfunction in heart failure^39^ (**Fig. S8d**). Our study therefore identifies individual genes with striking changes in the disease state, which are strong candidates for further investigation. Importantly, the large number of donors included in our study allowed us to identify significantly more DEGs across each contrast that would be possible from each of the individual studies (**Fig. 2e**).

### Some age and disease-associated gene pathways are discordant

To extend beyond individual genes that change across each contrast, we looked for pathway level changes using gene set enrichment analysis (GSEA)^40^ against the Gene Ontology Biological Process (GO:BP) gene set^41,42^ (**Fig. 2e; Fig. S21-S33; Tables S18-S30**). As expected, the sex contrast was enriched for the expected GO term of “dosage compensation” across all cell types. The development contrast displayed positive enrichments for pathways related to cell division and cytoplasmic translation, reflecting the higher proliferation of fetal cells. With aging, we observed decrease in collagen fibril organization in multiple cell types, consistent with increased ECM disarray during aging^43^. Interestingly, in 4 cell types, oxidative phosphorylation increases with age contrast to most aging literature, in which this pathway declines^44^. However, we do observe decline of oxidative phosphorylation in 6 cell types with disease. Our results suggest that decline of metabolic pathways may intrinsically be a hallmark of aging, but rather indicative of disease. Studies that do not separate these aging from disease may mis-attribute disease-related pathways as age-related ones.

### Cardiac disease is associated with fetal reactivation across almost all cell types

To better understand how changes across development, aging, and disease relate to each other, we performed intersection analysis to see if there is significant overlap between DEGs that change unidirectionally across pairs of contrasts (aging & development, development & disease, aging & disease) for a given cell type. For each pair of contrasts, we computed the ratio of DEGs that are unidirectionally concordant (either significantly up or down in both contrasts) over the total number of shared DEGs (**Fig. 3a**). To determine if this degree of overlap is statistically significant, we simulated a null distribution of overlap (**Methods**, **Fig. 3b**).

**Fig. 3:**
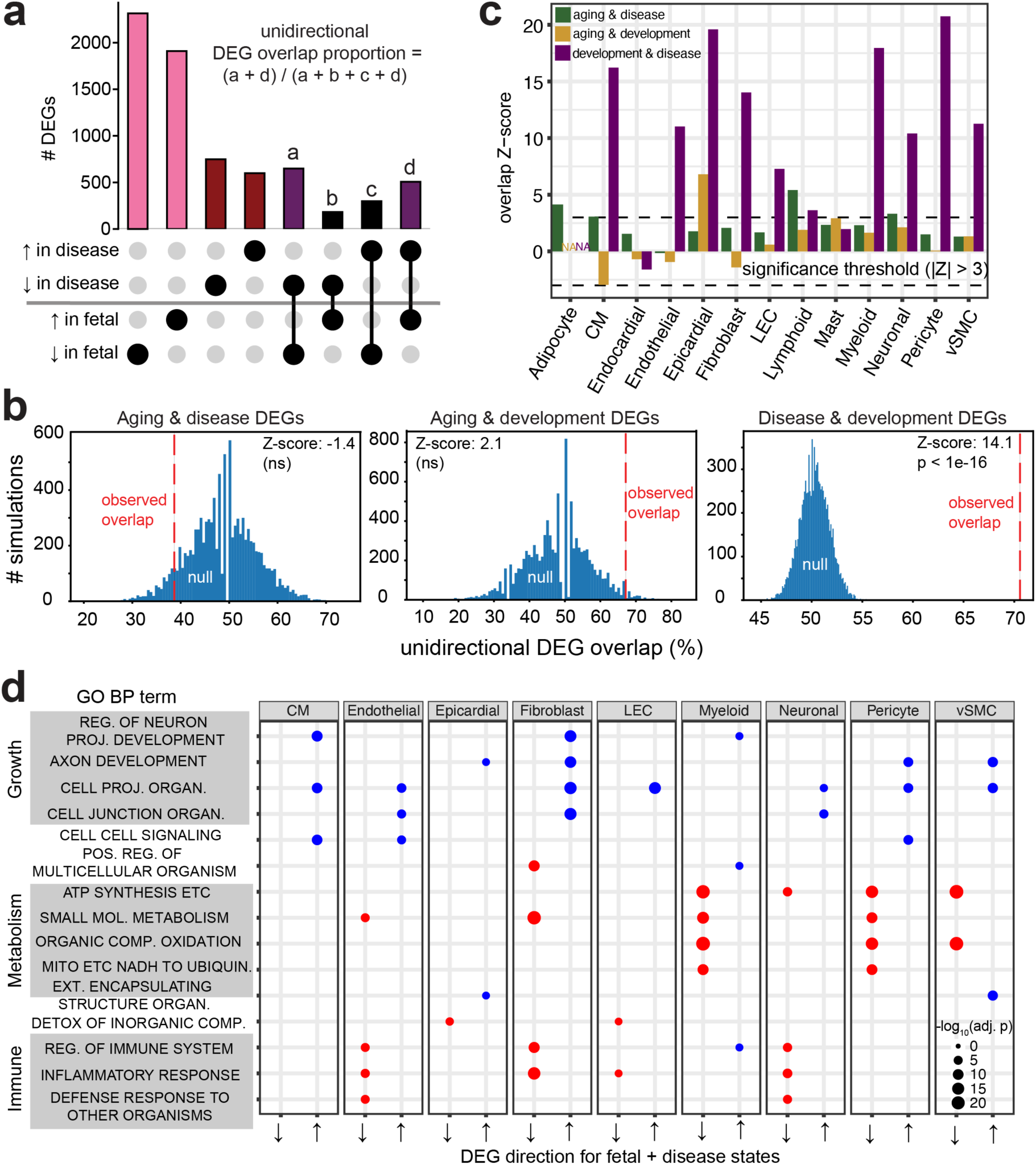
The diseased heart has strong transcriptomic overlap with the fetal heart. **a)** Upset plot, shown for fibroblasts, with the number of disease and developmental DEGs and the amount of overlapping DEGs, which include unidirectional up (a) and down DEGs (d). **b)** For fibroblasts, the observed versus expected degree of DEG overlap for three pairs of contrasts: development + aging (top), development + disease (middle), and disease + aging (bottom). The Z-score is the number of standard deviations that the observed ratio (a + d) / (a + b + c + d) is away from the null distribution, which accounts for DEG set sizes. **c)** The overlap Z-score across all cell types for all pairs of contrasts. **d)** Over-representation analysis for up and downregulated fetal and disease DEGs across all cell types that have a significant fetal Z-score.

Between fetal and aging DEGs, no cell types had a significant overlap (**Fig. 3c**). For aging and disease DEGs, the overlap was marginally significant for only a few cell types, corroborating that our GSEA results that aging and disease-associated processes are not strongly overlapping. In contrast, many cell types showed strong overlap between fetal and disease DEGs (**Fig. 3c**). This is consistent with the “fetal reactivation” hypothesis in heart disease, in which the stressed heart turns on developmental programs^45,46^. While studies on fetal reactivation have largely focused on cardiomyocytes^47^, we demonstrate it is pervasive, affecting nearly all major cardiac cell types in human hearts (**Fig. S34**). Our analysis are consistent with a recent study that observed in pediatric heart failure but focused on cardiomyocytes and fibroblasts^48^ and previous studies focused on bulk tissue^49^.

Whether fetal reversion is adaptive or maladaptive is unclear^45,46^. Therefore, to better characterize fetal reversion, we first identified whether any GO:BP pathways show fetal reactivation. Using overrepresentation analysis (ORA) on shared disease + fetal upregulated and downregulated DEGs (**Fig. 3d**), we found that fetal + disease downregulated genes are enriched for inflammatory and metabolic pathways in many cell types. Conversely, fetal and disease upregulated genes are enriched for cell growth and cell-cell signaling.

### Intercellular TGFβ signaling increases in both fetal and disease states

To further explore what potential cell-cell signaling pathways may be altered in the fetal and diseased state, we performed cell-cell communication (CCC) analysis, which uses statistically significant co-expression of known ligand and receptor pairings to predict cell-cell interactions. We used liana x tensorcell2cell^1,1–3^, a computational method that integrates communication scores across multiple CCC methods^51–55^ to compare cell-cell communication across biological states. Focusing on which CCC programs differ between fetal and diseased hearts relative to ND hearts. As program 5 had higher activity in both fetal and diseased hearts relative to ND hearts, we performed additional analysis of this program (**Fig. 4a**). Program 5 most strongly involves the communication between fibroblasts, pericytes, and vSMCs (**Fig. 4b; Fig. S35**), is most enriched for TGFβ signaling (**Fig. 4c**), and displays increased communication between several collagens and integrin isoforms (**Fig. 4d**). In the disease state, TGFβ contributes to excessive fibrosis, while in developmental contexts it is required for cardiac maturation^56^. Our cell-cell communication analysis thus suggests increased TGFβ signaling in both development and disease.

**Fig. 4:**
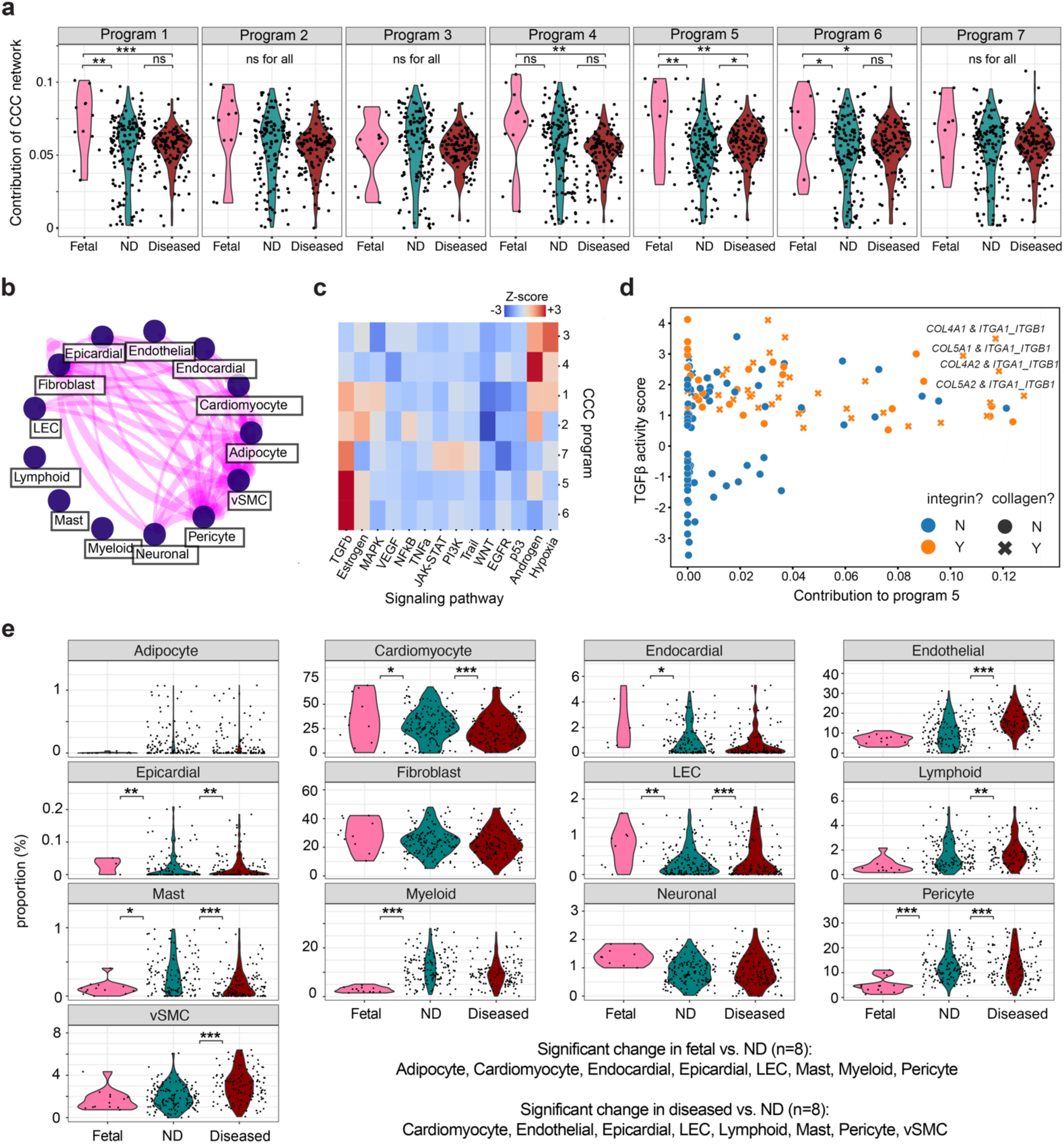
Cell-cell communication and cell-type proportional changes across development and disease. **a)** Cell-cell communications were inferred with liana x tensorcell2cell, which divides the overall cell-cell communication network in several programs and compares their relative contributions across biological states (fetal, non-diseased (ND), diseased). **b)** For program 5, the relative interaction strength of cell types. **c)** For each program, the enriched pathways, with positive enrichment indicating increased pathway activity. **d)** Major ligand-receptor interactions in factor 5 that contribute to TGFβ signaling. **e)** Cell type proportional differences across development and disease relative to ND hearts. Cell type proportional changes, compared by development and disease status. Significance: * (p<0.05), ** (p<0.01), *** p (<0.001), otherwise non-significant (ns).

### Development and disease are associated with significant but divergent differences in cell type proportions

In addition to differential expression of genes and pathways, changes can be caused by differences in cell type proportions. To investigate whether development, sex, age group, and disease are associated with cell type proportional changes, while regressing out technical batch effects, we used scanpro^40^ (based on propeller^57^). The only statistically significant sex difference was an increase of fibroblasts in females, and there were no significant cell type proportion changes with age (**Fig. S36; Tables S31-S34**). Across development, we identified 8 cell types with significant changes between fetal and postnatal young hearts. There is a postnatal increase in adipocytes, mast cells, myeloid cells, and pericytes and a postnatal decrease in cardiomyocytes, epicardial cells, endocardial cells, and LECs (**Fig. 4e**). These proportional changes are consistent with proliferation and diversification of non-myocyte cell types postnatally^57^. Disease is also associated with significant cell type proportion alterations. As expected, diseased hearts have fewer cardiomyocytes due to their replacement with fibrotic tissue^58^, but we did not observe a significant increase in fibroblasts, which could be due to increased fibrogenic activity per cell rather than increased overall abundance. We also observed an increase of endothelial cells, vSMCs, pericytes, and mast cells. Lymphoid cells in diseased hearts while myeloid cells had no significant change (**Fig. 4e**). In summary, there are profound but largely distinct developmental and disease-related cardiac cell type proportion changes, suggesting that while fetal reactivation involves significant reactivation of fetal genes transcriptionally, the overall cell type proportions in the disease heart do not revert towards a more fetal state.

### Integration of snATAC-seq datasets nominates TFs driving fetal reactivation

While snRNA-seq data analysis can reveal DEGs in a cell-type resolved manner, it does not identify possible upstream regulators of these genes. Since gene activation involves cell-type specific transcription factors (TFs) that bind to *cis*-regulatory elements, we sought to identify TFs that could explain fetal reactivation. Therefore, we performed an additional literature search to identify LV snATAC-seq datasets across developmental and disease contexts, resulting in snATAC-seq datasets from 95 different donors^14,18,59^. At the same time, we generated new snATAC-seq datasets of 5 fetal donors and 6 postnatal ND donors (**Fig. S37**). Altogether, the aggregation of 106 total donors (**Fig. 5a-b; Table S2**), while smaller than the snRNA-seq dataset, is the largest analysis of human cardiac snATAC-seq datasets to date. Our dataset is diverse in terms of sex (38 female, 68 male), age groups (17 fetal, 14 young, 44 middle, 31 old) and disease state (98 N, 13 Y).

**Fig. 5.**
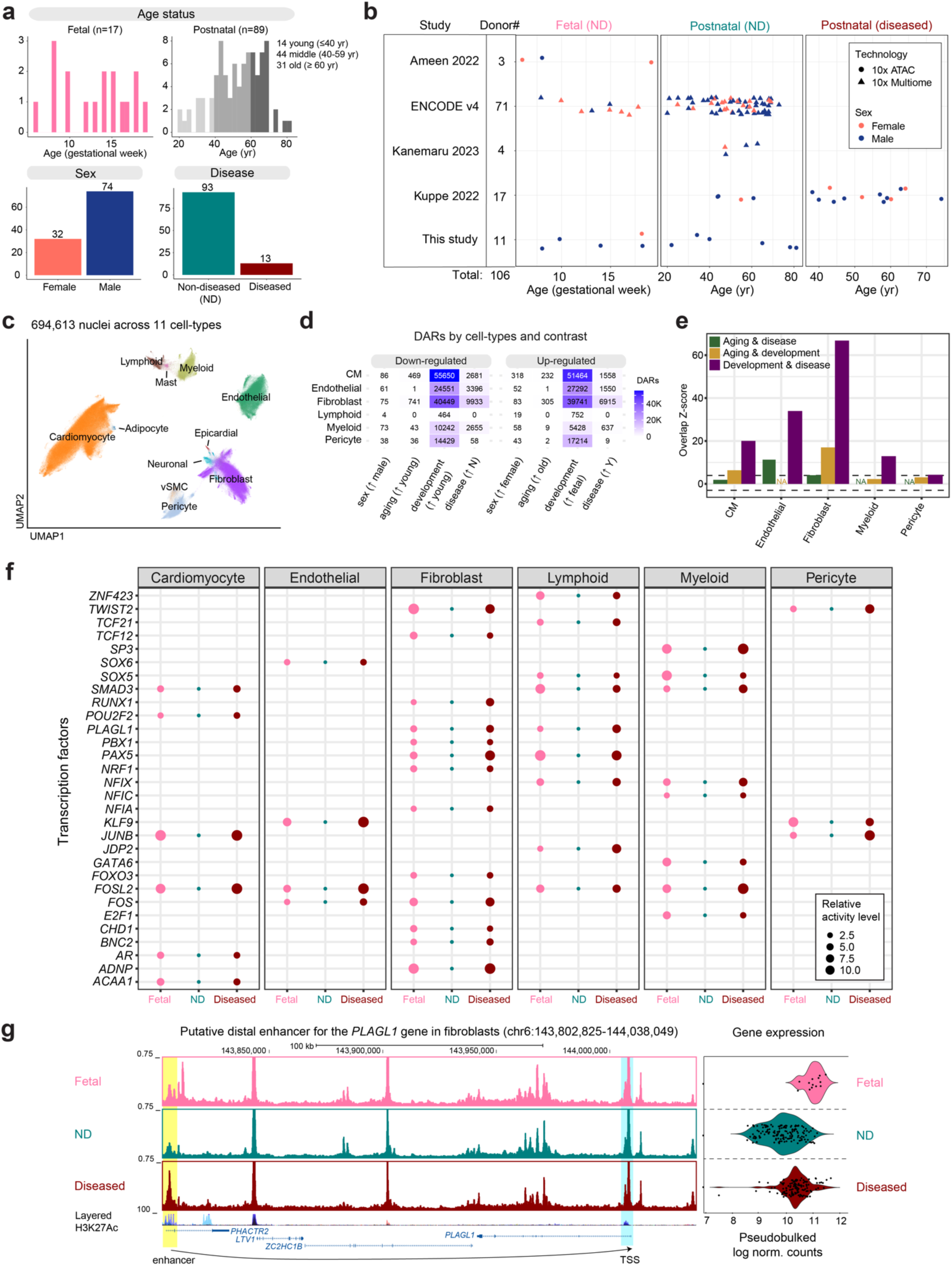
Single-nucleus ATAC-seq identifies putative TFs associated with fetal reactivation. **a)** snATAC-seq analysis involves 106 human donors, with diversity across age, sex and disease states **b)** Stratification of donors by study, age + disease status, and snRNA-seq technology platform **c)** UMAP of 694,613 nuclei across 10 resolved cell types **d)** Number of differentially accessible regions (DARs) per cell type x contrast combination, for cell types with at least >500 nuclei across fetal, ND, and diseased states. **e)** The Z-score for overlap of DARs across all cell types for all pairs of contrasts. **f)** Activator TFs with >2 fold increased SCENIC+ predicted activity in both fetal and disease states relative to ND are displayed. **g)** Genome browser view of putative *PLAGL1* enhancer in fibroblasts, which contains a *BNC2* motif, and may drive increased *PLAGL1* expression in fetal and diseased states. Layered H3K27ac track based on 7 ENCODE cell lines.

After quality control and batch integration, we obtained 694,613 high quality nuclei across 11 cell types (**Fig. 5c** and **S38a-d**; **Methods**). Distal peaks located >1kb from transcriptional start starts displayed 97.6% overlap with enhancers identified by a study that performed H3K27ac ChIP-seq on bulk heart tissue^2^ (**Fig. S38e-f**). Next, we called cell-type specific peaks using MACS3 ^60^. The promoter and gene bodies of marker genes displayed significantly higher accessibility in the corresponding cell type (**Fig. S39**). Altogether, this suggests that snATAC-seq recovers important *cis*-regulatory elements.

To confirm that fetal reactivation is observed at the epigenetic level, we performed differential accessibility analysis (DAR), using the pseudobulked DEG approach (**Fig. 2a**) for 6 cell types in which at least 500 nuclei were present for fetal, ND, and diseased states (cardiomyocyte, endothelial, fibroblast, lymphoid, myeloid, and pericyte). Concordant with the transcriptional level, we observed significantly more developmental and disease DARs than sex and aging DARs (**Fig. 5d; Tables S35-47**). Performing the same intersection analysis across contrasts, we confirm that the developmental and disease DARs strongly overlap across all annotated cell types (**Fig. 5e**), suggesting that fetal reactivation is at least partially driven by epigenetic regulation at the transcriptional level.

To identify putative TFs that could drive fetal reactivation, we integrated our snRNA-seq and snATAC-seq datasets using SCENIC+^61^, which links TFs to enhancer and target genes to construct gene regulatory networks (**Fig. 5f**). Using the activity score calculated by SCENIC+ for each cell type and state (fetal, ND, diseased), we identified 30 cell-type specific activator TFs for which there is >2 fold increased or decreased activity in both the disease and fetal states. Notably, we observe several AP-1 TFs (*FOS*, *FOSL2*, and *JUNB*) which play roles in both cardiac maturation and stress response^62,63^. We identify androgen receptor (*AR*) and *NFIX*, as reported in a bulk CAGE-seq study^64^, and *TCF12* and *TCF21*, as reported in a bulk ChIP-seq study^2^. In fibroblasts specifically, we observe fetal reactivation of *RUNX1*, which plays a role in many fibrotic diseases^65^ and has recently been mechanistically linked to fibroblast activation in heart failure^66^. Consistent with our cell-cell signaling analysis, we observe in cardiomyocytes and immune cells an increased activity of *SMAD3*, which is downstream of TGFβ and transcriptionally activates pro-fibrotic genes^67^. In some cases, fetal reactivation TFs are predicted to activate other such TFs. For example, a *BNC2* binding motif is located within an enhancer that promotes increased fetal and diseased *PLAGL2* expression, which has been shown to affect collagen expression^68^ (**Fig. 5g**). In summary, integrating snATAC-seq and snRNA-seq datasets allowed us to identify TFs and their gene regulatory networks (GRNs) that may contribute to fetal reactivation, and which may serve as perturbation targets for better characterizing the adaptive and maladaptive effects of fetal reactivation.

### Fetal reactivation is focalized in spatial niches

Finally, we explored whether fetal reactivating genes are localized in specific regions of the diseased heart or spatially homogeneously. To this end, we analyzed spatial transcriptomics from recently published 10X Visium datasets of non-diseased^14^ and MI^18^ donors (**Fig. 6a**). Using the set of shared fetal + disease DEGs for each cell type, we computed a “fetal reactivation score” for each spatial spot. Several of the diseased donors displayed high fetal reactivation scores, with the localization of fetal reactivation distinct from the localization of gene activity score obtained using a random set of genes DEGs (**Fig. S46-55**). Fibrotic and ischemic regions had significant higher proportion of spots with positive fetal reactivation scores than ND donors and the remote, non-infarct regions of MI donors dataset^14^ (**Fig. 6c**). Notably, several donors displayed significant spatial focalization as measured by the spatial autocorrelation metric Moran’s I, with specific niches of strong fetal reactivation (**Fig. 6b and 6d**). Additionally, the regions of fetal reactivation for each cell type are strongly overlapping, suggesting that different cell types may communicate to establish this fetal reactivated state (**Fig. 6e**). Altogether, our analysis establishes for the first time that fetal reactivation across diseased cells is spatially restricted to specific regions within diseased hearts.

**Fig. 6.**
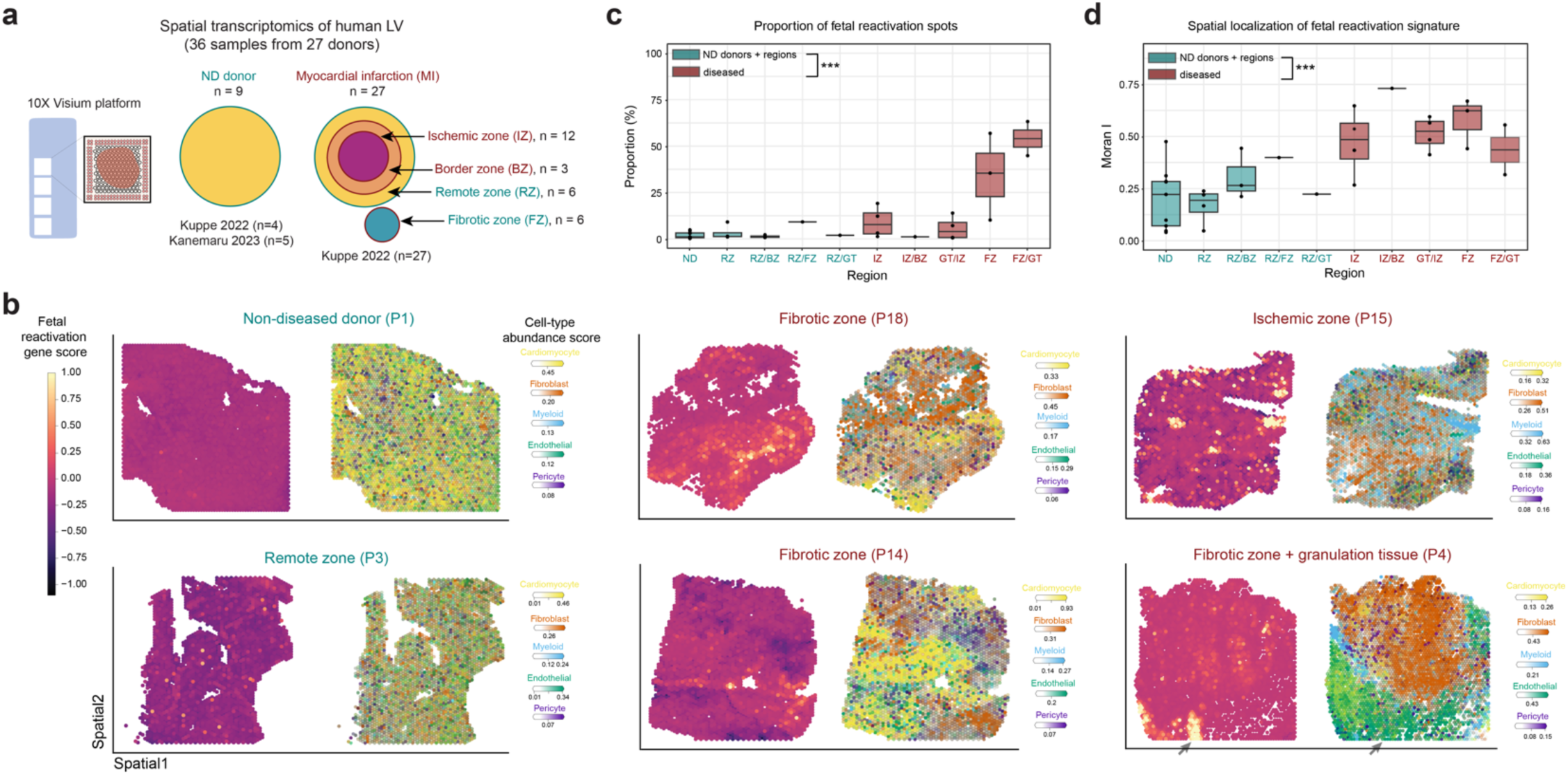
Spatial patterns of fetal reactivation in diseased human hearts. **a)** Overview of spatial transcriptomics datasets analyzed. **b)** Cell-type proportion weighted fetal reactivation scores (left) and abundance scores of five major cell-types (right: Cardiomyocyte – yellow, Fibroblast – red, Myeloid – blue, Endothelial – green, and Pericyte – Purple) in representative ND and diseased regions. A representative region of fetal reactivation in diseased heart is highlighted by arrow heads. **c)** Proportion of fetal reactivation spots in ND and diseased regions. **d)** Spatial localization, as calculated by Moran’s I, in ND and diseased regions. Significance: * (p<0.05), ** (p<0.01), *** p (<0.001), otherwise non-significant (ns).

## DISCUSSION

In this study, we performed the largest integrated analysis of snRNA-seq human heart datasets thus far. We also generated 19 new snRNA-seq donor and 11 snATAC-seq donor datasets, which will serve as a rich resource for future functional studies. We addressed several challenges with size and batch effects to leverage increased statistical power. Still, for human donor datasets, it may be difficult to fully resolve some batch effects (e.g., postmortem interval, exclusion criteria, biopsy strategy). Another major possible confounder is genetic variations among human donors. Though we lack genotyping information for most donors here, future genotyping efforts will enable studying the effect of genetic variants on expression at cell-type resolution^69^ and will complement efforts that have performed GWAS^70,71^ and bulk expression quantitative trait loci (eQTL) analyses.

By comparing transcriptomic profiles across 4 contrasts, we found few DEGs across sex and age but hundreds to thousands across development and disease states. By comparing the degree of shared DEGs across contrasts, we found no strong overlap between age and disease DEGs. Some pathways such as oxidative phosphorylation even change in opposing directions between aging and disease despite age being a strong risk factor for cardiac disease. This discrepancy could be due to diseased samples being mostly end-stage explanted hearts. Alternatively, aging may not be intrinsically a disease-increasing process and “healthy” aging may diverge from disease. Whether there are age-related pathways that protect the heart from disease is an interesting possibility, in line with efforts to study cardiac resilience^72^.

In contrast to aging DEGs, we found developmental DEGs to strongly overlap with disease DEGs. We demonstrate that shared fetal and disease pathways span many different biological processes, including TGFβ signaling. To better identify upstream regulators that could be driving fetal reactivation, we integrated a large snATAC-seq dataset to nominate a small subset of fetal-reactivation TFs, which include known TFs associated with cardiac phenotypes and novel ones. Our focused re-analysis of spatial transcriptomics data identified the spatial focalization of fetal reactivation. The significance of these fetal reactivation niches, and whether they have adaptive, pro-regenerative capabilities, or are primarily maladaptive requires further study.

Altogether, our study identifies exciting aspects of human cardiac biology. While the direct human relevance of these data is a major strength, we only analyzed nuclear expression, which does not account for post-transcriptional, translational, and metabolic changes. Moreover, despite generating the largest multiomic atlas to date, we do not see saturation of aging DEGs (**Fig. 2d**), suggesting that a larger sample size may be required for detecting more age-related changes. Finally, our findings are associative and would require experimentation to establish causality. Though model organisms may be suitable for testing some of these associations, some of these findings may be only applicable to humans. As iPSC-derived cardiac organoid protocols improve^73^, they may help bridge the gap. The combination of this work, which has identified numerous DEGs and putative fetal reactivation TFs, with more targeted mechanistic studies will help further understand the human heart across its entire phenotypic spectrum.

## ACKNOWLEDGEMENTS

We are grateful to members of the Wu lab for helpful discussion. This work was in part supported by a National Heart Lung and Blood Institute (NHLBI) grant DP2-HL142044 (to H.W.). W.G. and H.W. are supported by a NHGRI grant U01-HG012047 and a Leducq Foundation grant.

## AUTHOR CONTRIBUTIONS

W.G and H.W. conceived of the project, in collaboration with K.M, and K.S. K.B. and K.M. procured all postnatal human heart tissue samples. K.S. procured all fetal human heart tissue samples. W.G. combined Penn datasets with external datasets and performed all integrative sn/scRNA-seq, snATAC-seq, and spatial transcriptomics analyses. P.H. generated all Penn snRNA-seq and snATAC-seq datasets and performed Penn dataset analyses, with help from Q.Q and K.X. W.G. and H.W. wrote the manuscript, with contributions from all authors.

## Methods

### Human cardiac tissue procurement

Newly generated snRNA-seq and snATAC-seq were prepared from the adult hearts deposited in the Penn Heart Tissue Biobank. Hearts were harvested at time of cardiac surgery from subjects with heart failure (2 DCM patients) or from non-diseased donor hearts that were unable to be transplanted within the surgical window. Hearts were transported to the research lab in cardioplegic solution to prevent ischemic damage, and single nucleus suspensions of tissue were biopsied from the left ventricular wall. Written consent for research use of these tissue was obtained from the subject providing tissue or next of kin and approved by the University of Pennsylvania institutional review board.

Prenatal human hearts (8-18 wpf) were obtained from donors undergoing elective abortion at the University of Pennsylvania, with all donors providing written consent for the use of the conceptus in research. Fetal age was determined via ultrasonographic measurement of crown-rump length or head circumference. The heart tissues were submerged in RPMI-1640 medium and meticulously dissected under a stereomicroscope to remove surrounding connective and adipose tissues.

### Human cardiac nuclei isolation and FANS

Human cardiac tissues were flash-frozen in liquid nitrogen and stored at -80°C before nuclear isolation. Nuclei were isolated and purified as previously described with some modifications^12^. Briefly, 14 ml sucrose cushion (1.8M sucrose (Sigma-Aldrich, RNase and DNase free, ultra-pure grade), 10 mM Tris-HCl pH 8.0, 3 mM MgAc2 (Sigma-Aldrich), and protease inhibitor cocktail (Roche) was added to the bottom of centrifuge tubes (Beckman Coulter). Using a glass homogenizer (Wheaton), a frozen mouse cortical sample was dounce homogenized (19 times with loose pestle followed by 4 times with tight pestle) in 12mL homogenization buffer (0.32 M sucrose, 5 mM CaCl2 (Sigma-Aldrich), 3 mM MgAc2, 10 mM Tris-HCl pH 8.0, 0.1% Triton X-100 (Sigma-Aldrich), 0.1 mM EDTA (Invitrogen), protease inhibitor cocktail (Roche). Homogenates (∼12 ml) were carefully layered onto the sucrose cushion in the centrifuge tubes, and 10mL homogenization buffer was added atop the homogenates. The tubes were then balanced and centrifuged in a Beckman Coulter L7-65 Ultracentrifuge at 82,705 g at 4°C for 120 min using a Beckman Coulter SW28 swing bucket rotor (Beckman Coulter). The supernatant was carefully removed via aspiration. 1ml chilled DPBS was added to resuspend the nuclear pellet, and nuclei were subsequently transferred to a 1.5-ml tube. Nuclei were pelleted at 500 g for 10 min at 4°C and then resuspended in 0.01% BSA (Sigma-Aldrich) in DPBS. After resuspension, nuclei were filtered through a 40-μm cell strainer (Fisher Scientific), visually inspected for morphology and quality assurance, and counted using a Fuchs-Rosenthal counting chamber before FANS.

Isolated nuclei from were resuspended in 1.5 ml blocking buffer (1× PBS, 0.5% BSA (Sigma A4503), Roche Complete Protease Inhibitor without EDTA) and blocked for 20 min at 4°C with rotation. After blocking, nuclei were incubated with DAPI (1:1000) for 5 min. Nuclei were then washed, pelleted, and resuspended in 1mL FACS buffer (1× PBS, 1% BSA (Sigma A4503), Roche Complete Protease Inhibitor without EDTA), and passed through a 35 µm strainer (Corning #352235) in preparation for flow cytometry. FANS was performed using a BD Biosciences Influx cell sorter at the University of Pennsylvania Flow Cytometry and Cell Sorting Facility. Singlet nuclei were identified using DAPI fluorescence.

### Single-nucleus RNA sequencing (sNucDrop-seq)

Single nucleus RNA sequencing analysis of FANS-sorted human cardiac nuclei was performed using sNucDrop-seq protocol^12,74^. Briefly, FANS-sorted nuclei were individually diluted to a concentration of 100 nuclei/mL in DPBS containing 0.01% BSA. Approximately 1.25 ml of this single-nucleus suspension was loaded for each sNucDrop-seq run. The single-nucleus suspension was then co-encapsulated with barcoded beads (ChemGenes) using an Aquapel-coated PDMS microfluidic device (μFluidix) connected to syringe pumps (KD Scientific) via polyethylene tubing with an inner diameter of 0.38mm (Scientific Commodities). Barcoded beads were resuspended in lysis buffer (200 mM Tris-HCl pH8.0, 20 mM EDTA, 6% Ficoll PM-400 (GE Healthcare/Fisher Scientific), 0.2% Sarkosyl (Sigma-Aldrich), and 50 mM DTT (Fermentas; freshly made on the day of run) at a concentration of 120 beads/ml. The flow rates for nuclei and beads were set to 4,000 ml/hour, while QX200 droplet generation oil (Bio-rad) was run at 15,000 mL/hr. A typical run lasts 20 min. Droplet breakage with Perfluoro-1-octanol (Sigma-Aldrich), reverse transcription and exonuclease I treatment were performed, as previously described, with minor modifications. For up to 120,000 beads, 200 μl of reverse transcription (RT) mix (1x Maxima RT buffer (ThermoFisher), 4% Ficoll PM-400, 1 mM dNTPs (Clontech), 1 U/mL RNase inhibitor, 2.5 mM Template Switch Oligo (TSO: AAGCAGTGGTATCAACGCAGAGTGAATrGrGrG) ^1^, and 10 U/ mL Maxima H Minus Reverse Transcriptase (ThermoFisher) were added. The RT reaction was incubated at room temperature for 30min, followed by incubation at 42°C for 120 min. After determining an optimal number of PCR cycles for amplification of cDNA using real-time PCR analysis (Applied Biosystems QuantStudio 7 Flex) ^2^, aliquots of 6,000 beads was amplified by 50-μl PCR reactions (25 μl of 2x KAPA HiFi hotstart readymix (KAPA biosystems), 0.4 μl of 100 mM TSO-PCR primer (AAGCAGTGGTATCAACGCAGAGT) ^1^, 24.6 μl of nuclease-free water) with the following thermal cycling parameter (95°C for 3 min; 4 cycles of 98°C for 20 sec, 65°C for 45 sec, 72°C for 3 min; 10-12 cycles of 98°C for 20 sec, 67°C for 45 sec, 72°C for 3 min; 72°C for 5 min, hold at 4°C). After two rounds of purification with 0.6x SPRISelect beads (Beckman Coulter), amplified cDNA was eluted with 10 μl of water. We then tagmented cDNA using the Nextera XT DNA sample preparation kit (Illumina, FC-131-1096), starting with 550 pg of cDNA pooled in equal amounts, from all PCR reactions for a given run. Following cDNA tagmentation, we further amplified the tagmented cDNA libraries with 12 enrichment PCR cycles using the Illumina Nextera XT i7 primers along with the P5-TSO hybrid primer ^1^. After quality control analysis by Qubit 3.0 (Invitrogen) and a Bioanalyzer (Agilent), libraries were sequenced on an Illumina NextSeq 500 instrument using the 75-cycle High Output v2 Kit (Illumina). We loaded the library at 2.0 pM and provided Custom Read1 Primer (GCCTGTCCGCGGAAGCAGTGGTATCAACGCAGAGTAC) at 0.3 mM in position 7 of the reagent cartridge. The sequencing configuration was 20 bp (Read1), 8 bp (Index1), and 60 bp (Read2).

### Single-nucleus ATAC-seq (10x Genomics)

Using the nuclei isolation protocol detailed for sNucDrop-seq protocol, nuclei were aliquoted for single nucleus ATAC sequencing analysis of human cardiac nuclei. snATAC-seq was performed using the 10X Genomics ATAC v1 kit, with a target of 10K nuclei were incubated with Tn5 transposase per reaction. After tagmentation, nuclei were encapsulated using Chromium Chip E in the manufacturer’s Chromium Controller. The library was sequenced on an Illumina NextSeq 500 using these read parameters: 50 bp for Read1, 8 bp i7 index, 16 bp i5 index, and 50 bp for Read2.

### Penn snRNA-seq dataset preprocessing

Raw FASTQ files were aligned against the hg38 reference genome (refdata-cellranger-arc-GRCh38-2020-A-2.0.0) using STARSolo (STAR v2.7.11a), using the parameters --soloUMIlen 8 --soloCBlen 2 -- soloFeatures Gene GeneFull --soloCellFilter EmptyDrops_CR. The raw and filtered (based on the UMI threshold set by EmptyDrop_CR) directories produced were passed as inputs to SoupX for ambient RNA removal. The filtered, ambient RNA removed count matrices were concatenated together and analyzed together using scanpy. We performed scVI-based integration using donor_id as the batch key. Using the KNN graph in the scVI embedding, we called Leiden clusters with a resolution of 0.5 and annotated clusters based on marker genes. The primary marker genes that assisted in this manual annotation included known markers reported in the literature^6,7,14^ including *TTN/RYR2* (Cardiomyocyte); *DCN/COL3A1* (Fibroblast); *PPARG/ADIPOQ* (Adipocyte); *VWF* (Endothelial); *POSTN + VWF* (Endocardial); *ITLN1*/*MSLN/WT1* (Epicardial); *MMRN1* (LEC); *MS4A1/SKAP1* (Lymphoid); *KIT* (Mast); *CD163* (Myeloid); *NRXN1/XKR4* (Neuronal); *EGLAM/DLC1/PDGFRB* (Pericyte); *MYH11/CARMN* (vSMC). More information about the Penn snRNA-seq dataset is provided in Fig. S1.

### Penn snATAC-seq dataset preprocessing

Raw FASTQ files were aligned against the hg38 reference genome (refdata-cellranger-arc-GRCh38-2020-A-2.0.0) using cellranger-atac 2.1.0 under default parameters to produce fragment files. These fragment files were inputted to SnapATAC2 for preprocessing, which included calculating the number of fragments and TSS enrichment score per barcode. We filtered the cells to those with at least 1000 unique fragments and a TSS enrichment score > 3 and combined all samples together. We then performed harmony batch integration (using donor_id as the batch key), followed by SnapATAC2’s spectral embedding. Leiden clusters were identified using a resolution of 0.5. Cluster annotation was performed using MAGIC imputed gene expressions of marker genes noted in the snRNA-seq section above. Peak calling was performed using MACS3. For information about the Penn snATAC-seq dataset is provided in Fig. S37.

### External snRNA-seq dataset processing

External datasets^7,13–20,23^ were downloaded according to the data availability sections of each paper. The link for downloading the processed data is provided in Table S1 under the “RNA” section. The accessions of the 62 ENCODE v4 snRNA-seq datasets we downloaded are provided under the “ENCODE RNA” section^75^.

### snRNA-seq integration and cell type annotation

The raw count matrices (filtered to remove droplets containing non-cells) across all the studies were concatenated together using Scanpy^76^. For ambient RNA removal for our new dataset and published datasets in which unfiltered count matrices were readily available, we performed ambient RNA removal using SoupX^77^. For datasets with only processed filtered count matrices, these count matrices were used, as ambient RNA removal was already performed by each study using either SoupX or cellbender^78^. We used scVI^25^ to remove batch effects at the UMAP embedding level, with tech_plus_study serving as the batch and donor_id serving as a categorical covariate in the model. scVI batch corrected counts for all genes were stored, using the ENCODE v4 study as the conditional batch as it had the largest number of non-diseased donors. These batch corrected counts were used for batch-corrected analysis with liana x tensorcell2cell^50^, as recommended. Raw counts were used for differential gene expression, as batch corrected counts have been shown to distort differential gene expression analysis^9^. Leiden clustering at a resolution of 0.5 was performed using Scanpy. To remove possibly batch-induced clusters, Leiden clusters in which >75% of cells came from a single study were removed. After this filtering, the top 50 marker genes per Leiden cluster, including marker genes noted above in the Penn dataset analysis, were used to independently annotate each Leiden cluster, agnostic to the cell annotations from the original studies. After this first step of annotation, we subsetted the count matrix to each individual cell type. For each of these cell types, we re-examined the UMAP embedding and confirmed marker gene expression. In this cell-type filtered count matrix, cells that segregated away from the main cluster and/or that expressed marker genes for other cell types may represent doublets, so we manually removed these cells. Finally, we combined all filtered cell type count matrices to produce a final dataset 2,305,964 nuclei across 13 different cell types.

### Differential gene expression analysis

We performed donor-based pseudobulked differential gene expression, by aggregating the raw counts of all nuclei of the same cell type annotation for each donor. This produces a donor x gene matrix for all 13 cell types. We performed differential gene expression with pydeseq2 ^28^, the python-based implementation of DESeq2 ^27^, using this design matrix: expression ∼ sex + age_group + disease_binary + tech_plus_study. We opted to include age as a categorical variable rather than a continuous variable as all other variables were categorical and to account for possible non-linear changes with age. The age groups therefore included fetal, young age (<40), middle age (40-59), and old (>60), with age cutoffs chosen to construct sufficient sizes per age group. As several of the different types of cardiac disease are represented only by 1-2 studies, the disease status was binarized to either diseased or non-diseased for this analysis. Many different droplet-based technologies were used for snRNA-seq data generation, including Dropseq, 10X Genomics 3’ v1-v3, and 10X Multiome v1. As some studies used multiple different technologies, we created a variable called tech_plus_study that combines both the study and technology information together. Without regressing out this technical batch effect, the pseudobulked PC1 and PC2 show significant correlation with tech_plus_study. Using limma^79^ to remove batch effects at the pseudobulked level reduced the clear segregation of samples by technology and study (**Fig. S3**).

DESeq2 returns a log2 fold change (FC) and false discovery rate (FDR)^80^ adjusted p-value. We considered significant upregulated genes to be those with log2FC > 0.5 (corresponding to 1.41-fold change) and adjusted p < 0.05. We considered significant downregulated genes to be those for log2FC < - 0.5 (0.707-fold change) and adjusted p < 0.05. The interpretation of the “up” and “down” depends on the contrast. The contrasts we focused on for this study include sex: male vs. female, development: fetal vs. young (age group), aging: young vs. old (age group), and disease: Y vs. N (disease_binary), as indicated in Fig. 2a. As an example, an “upregulated” gene in the developmental contrast (fetal vs. young) would represent a gene significant higher in fetal donors (group 1 of contrast) compared to the postnatal young donors (group 2 of contrast). A “downregulated” gene would represent a gene significantly higher in the postnatal young donors.

### Gene set enrichment analysis (GSEA) and over-representation analysis (ORA)

GSEA was performed using gseapy^81^, a fast python implementation of GSEA^82^. To reduce redundancy in gene set names, we constructed a custom gene set based on the Gene Ontology Biological Processes (GO:BP)^41,42^ gene set. Using a similar approach to Shen et al. 2024^83^, we calculated the Jaccard similarity between all pairs of gene sets, which is equal to the number of genes shared between the two gene sets over the union of all genes present in either gene set. We then computed a distance metric as 1 – Jaccard similarity (which ranges from 0 to 1) and performed hierarchical clustering. Clusters were defined as gene set groups for which the distance is less than 0.7. We randomly selected one gene set per cluster and used this gene set of 3913 less redundant gene sets, in place of the original GO:BP list, which contains 7647 gene sets.

After performing the DE workflow above for each contrast and cell type, we obtained a results table for each gene, including log2 fold change. Genes were sorted in terms of decreasing log2 fold change and then passed into gseapy prerank() function. Significant GSEA results were defined as gene sets with an FDR-adjusted p value < 0.05. Up to 5 significant gene sets with positive and negative enrichment scores per cell type are displayed in **Fig. S18-30**. To enhance readability, only gene sets that were significant in at least 5 cell-type x contrast pairs were retained in **Fig. 2d**.

For over-representation analysis, the list of fetal reactivation genes was inputted against the Jaccard simplified gene sets. Fetal reactivation genes were defined as the DEGs that are shared between the developmental and disease contrasts that are unidirectional. For example, fetal reactivation “up” genes are those that are higher in the fetal hearts relative to postnatal young heart, and higher in diseased hearts relative to postnatal non-diseased hearts.

### DEG/DAR unidirectional overlap significance of across contrasts

To determine whether there is a significant unidirectional overlap between DEGs in two contrasts within a given cell type, we simulated expected overlap under a null distribution that accounts for the size of the gene sets. For example, for two contrasts 1 (e.g., old vs young) and 2 (e.g., fetal vs young), there are 4 total gene sets of varying size: up in 1 (W), down in 1 (X), up in 2 (Y), down in 2 (Z). W and X are disjoint (no overlapping elements) as no gene can both be up or down regulated in the same contrast. Y and Z are also disjoint. The amount of unidirectional overlap is the sum of total number of genes that intersect between W and Y (= a in **Fig. 3a**) plus the total number of genes that intersect between X and Z (= d in **Fig. 3a**). To normalize this quantity, we divide by the total number of shared DEGs. To simulate a null distribution, we randomly sample W genes (“up in 1”) from the gene universe (all genes represent in the count matrix for that cell type) and then remove those W genes to sample X (“down in 1”) genes. We then sample Y genes (“up in 2”) from the gene universe and then remove those Y genes to sample Z genes (“down in 2”) genes. We calculate the degree of overlap across 1000 simulations to obtain an approximately normal null distribution with estimated μ and σ. The Z-score of the observed degree of overlap (X) is calculated as (X - μ) / σ. A positive Z-score would indicate unidirectional DEG overlap between two contrasts greater than expected by chance, while a negative Z-score would indicate overlap lower than expected by chance (mutual exclusivity). A Z-score greater than 3 was considered significant.

### Cell-cell communication analysis

Liana x tensorcell2cell was performed on scVI-batched integrated counts, as recommended by the software developments. The counts were all transformed as if generated from the ENCODE v4 Multiome batch, using the scVI transform_batch() function. Without performing this batch correction, we obtained spurious significant differences in cell-cell communication that seemed to be driven by sequencing depth differences and other study-specific technical artifacts. We followed the tutorial titled “Intercellular Context Factorization with Tensor-Cell2cell” to build a 4D tensor, representing contexts, interactions, sender cell types, and receiver cell types. To reduce jargon, “program” was used in this manuscript instead of “factor”. The optimal number of programs identified using optimal rank estimation was 7. Next, we identified programs that showed significant differences across 3 age + disease status conditions: fetal non-diseased, postnatal non-diseased, and postnatal diseased using the c2c.plotting.context_boxplot() function. Identification of enriched pathways was performed using decoupleR ^84^ with PROGENy ^85^ gene sets.

### Cell type proportional analysis

Cell type proportional analysis was performed by first normalizing # nuclei for each cell type by the total number of nuclei per donor. To reduce noise from sampling, only donors with at least 1000 high-quality nuclei were considered for this analysis, which removed 7 donors, leaving 292 donors. To account for batch effects such as technology and study, we used scanpro^40^, a python implementation of the propeller^57^ package, which performs mean-variance stabilized estimation of cell type proportions. This method accounts for the interdependent nature of cell type proportions (as all proportions within a donor must sum to one) while also accounting for batch effects. To identify sex, aging, and disease related changes in cell type proportion, we filtered the donors studied to postnatal donors only (279 donors). In scanpro, we then specified the contrast of interest (either ‘sex’, ‘disease_binary’, ‘age_group’) as the condition column ‘conds_col’ and the donor id as the ‘samples_col’. The other covariates in the model included all but the covariate in ‘conds_col’ along with ‘tech_plus_study’ as a batch covariate. To identify fetal-specific differences in cell type proportion, we filtered the count matrix to only to fetal and young postnatal non-diseased donors (44 donors) and then performed differential testing with ‘age_group’ as the contrast (‘conds_col’) with ‘sex’, ‘disease_binary’, and ‘tech_plus_study’ as covariates. For binary contrasts (sex: male vs. female; age_group: fetal vs. young, disease_binary: Y vs N), the t-test default was used to determine statistical significance. The resultant p-value was FDR-corrected to account for testing all cell types. For aging related changes, there are three categories (young, middle, old), so the ANOVA test was used for statistical significance.

### External snATAC-seq dataset processing

External datasets^14,18,23,59^ were downloaded according to the data availability sections of each paper. The link for downloading the processed data is provided in Table S1 under the “ATAC” section. The accessions of the 71 ENCODE v4 snATAC-seq datasets we downloaded are provided under the “ENCODE RNA” section^75^.

### snATAC-seq integrative preprocessing

We combined Penn fragment files with previously published fragment files from the other datasets as input to SnapATAC2^86^. Using SnapATAC2, we performed quality control filtering by TSS enrichment score and number of fragments per cell were used to define per-sample cutoffs. After combining high quality cells across samples, we performed spectral embedding on the tile count matrix (5 kb tiles across the genome) and Harmony integration. Leiden clustering with resolution of 1 was performed using the spectral embedding nearest neighbor graph. Cell type annotation of these clusters was performed using two approaches. First, gene expression using imputed using MAGIC^87^ and marker genes for snRNA-seq were used to assist with cell type annotation. While this approach worked for more common cell types, we found the second approach, which involved label transferring of multiome cells, to be more effective at annotating smaller clusters with rarer cell types (e.g., adipocyte, epicardial). In this second approach, we filter the snATAC-seq nuclei to those produced with 10X multiome and examined the proportion of snRNA-seq annotations for each Leiden cluster. Leiden clusters with >90% sharing a single snRNA-seq annotation were annotated as that cell type. For clusters with >20% of a rare cell type, these leiden clusters were kept and then marker gene imputation with MAGIC was performed to identify likely true cells. All other leiden clusters were removed, as they may represent doublets. The inability to identify distinct clusters for two cell types present in snRNA-seq (LEC, Endocardial) may be less pronounced epigenetic vs. transcriptomic differences for LEC and Endocardial cells, which were located within the Endothelial cluster. After cell type annotation, reads were aggregated at the cell type level for peak calling with MACS3^60^, producing a cell by peak count matrix. Harmony batch integration and UMAP dimensionality reduction was reperformed on this matrix with peaks as features to generate the final UMAP plot in Fig. 5c.

### Differential accessible region (DAR) analysis

Similar to DEG analysis, we performed donor-based pseudobulked accessibility analysis, by aggregating the raw counts of all nuclei of the same cell type annotation for each donor. We performed differential accessibility analysis with pydeseq2^28^ using this design matrix: expression ∼ sex + age_group + disease_binary + technology. Study was not included in the design matrix to reduce collinearity, as all the diseased cell types come from the same study (Kuppe 2022).

### Intersection of snATAC-seq peaks with other feature sets

The cell-type specific peaks called by MACS3 were merged into 500 nucleotide regions using SnapATAC2. The peak coordinates were converted to BED format^88^. The Spurrell et al. 2022 list with 33317 enhancers^2^, called from the healthy donors was also converted to BED format. For intersection of Spurrell enhancers and snATAC-seq peaks, snATAC-seq peaks were filtered only those that are located >1 kb from a transcriptional start site were used for intersection with Spurrell enhancers, following their definition of enhancers. The distance from the nearest TSS was computed using ChIPSeeker^89^. Due to the significant length discrepancy between snATAC-seq peaks (fixed length of 500 nt) and then Spurrell enhancers (median length 4244 nt) we calculated the number of Spurrell enhancers with at least one snATAC-seq overlapping it, using: bedtools intersect -a <SPURRELL_BED> -b <SNATAC_ENHANCER_PEAKS_BED> -c. Using bedtools shuffle, we generated shuffled snATAC-seq peak sets and then used bedtools intersect to obtain a null distribution.

### SCENIC+ gene regulatory analysis

SCENIC+^61^ integrates snATAC-seq and snRNA-seq to infer gene regulatory networks. As SCENIC+ does not scale to millions of cells, we created subsampled 750 nuclei for cell types in which we had at least 750 nuclei for both ATAC and RNA modalities across fetal, ND, and diseased donors. Therefore, we only performed analysis on 6 cell types: cardiomyocyte, endothelial, fibroblast, lymphoid, myeloid, and pericyte. The rest of the analysis was performed using the SCENIC+ tutorial (https://scenicplus.readthedocs.io/en/latest/tutorials.html). Using the area under the curve (AUC) metric that measures the activity of a TF-driven GRN in a particular cell type and postnatal + disease context, we identified GRNs with at least a 2-fold increased or decreased AUC in the fetal and diseased cells when compared to non-diseased postnatal cells. We filtered these GRNs to activator regulons where TF expression and chromatin accessibility of TF binding sites is positively correlated with gene expression (+/+ in SCENIC+ notation). Repressor regulons were removed from analysis as recommended by the SCENIC+ authors.

### Spatial transcriptomics analysis

Processed spatial transcriptomics datasets from Kuppe et al. 2022^18^ and Kanemaru et al. 2023^14^ were downloaded. Combined spot level expression of fetal reactivation genes were calculated using scanpy’s score_gene() function^76^ using the gene set of shared upregulated DEGs in the developmental and disease contrasts. A weighted fetal reactivation score was computed by multiplying the per-cell-type fetal reactivation score by the cell type proportion, computed using cell2location^90^. The proportion of fetal reactivation spots for each sample was determined as the (# spots with fetal reactivation score > 0) / (# total spots). For spatial autocorrelation, Moran’s I was calculated for the fetal reactivation scores for each cell type.

### Genome browser visualization

Visualization of genome browser tracks in **Fig. 5f** was performed using UCSC Genome browser. Visualization of promoter accessibility for marker genes was performed using ArchR plotBrowserTrack()^91^.

## Data availability

The aggregated snRNA-seq dataset with 2,305,964 nuclei, the aggregated snATAC-seq dataset with 694,613 nuclei, and the raw expression matrices and raw sequence files generated for this study (labeled as “This study” in Fig. 1a and Fig. 5a) will be available on the Gene Expression Omnibus (GEO) and dbGaP upon peer-reviewed publication.

## Code availability

All code used for analysis in this study are provided in Github: https://github.com/wgao688/Human-cardiac-multiome-analysis.

---

**Supplemental Figure 1.**
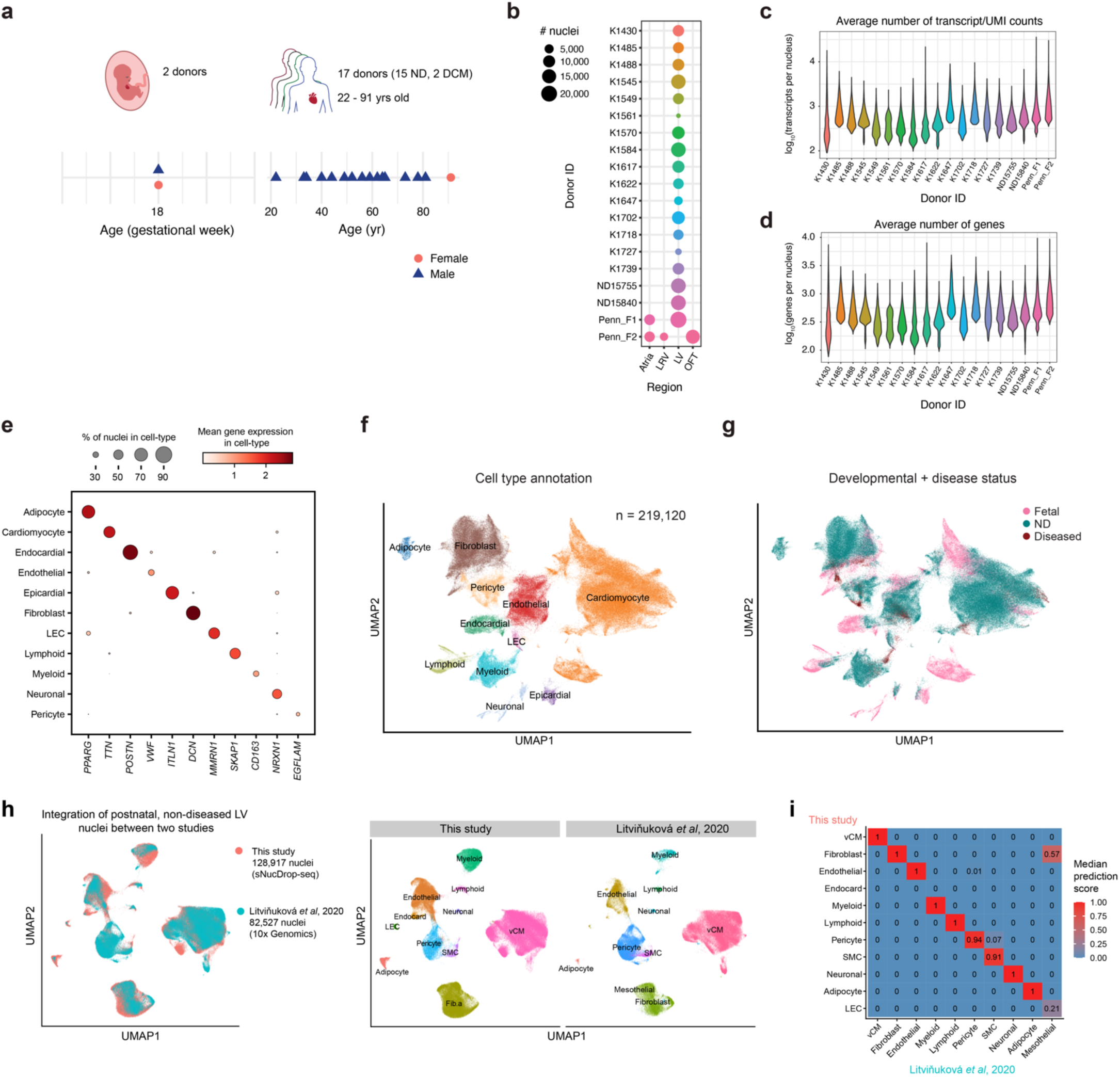
Penn single-nucleus RNA-seq dataset: **(a)** Summary of new donors included in this study, including 2 fetal donors and 17 postnatal donors (ages 22-91). 2 donors had a diagnosis of dilated cardiomyopathy (DCM). **(b)** Cardiac regions sampled per donor and number of high-quality nuclei per region x donor. **(c-d)** Summary statistics of snRNA-seq libraries with different nuclei isolation methods, including number of genes and transcripts/UMIs per nucleus, in log scale, respectively. **(e)** Dotplot showing marker gene expression across annotated cell-types **(f-g)** UMAP visualization of cardiac cells colored by annotated cell-types and developmental + disease status, respectively. **(h)** UMAP visualization of integrated snRNA-seq datasets of adult human left ventricle (LV, non-diseased) from this study and a published study (Litvinukova et al. 2020). **(i)** Heatmaps showing the median prediction scores of paired clusters between this and published studies using label transfer.

**Supplemental Figure 2.**
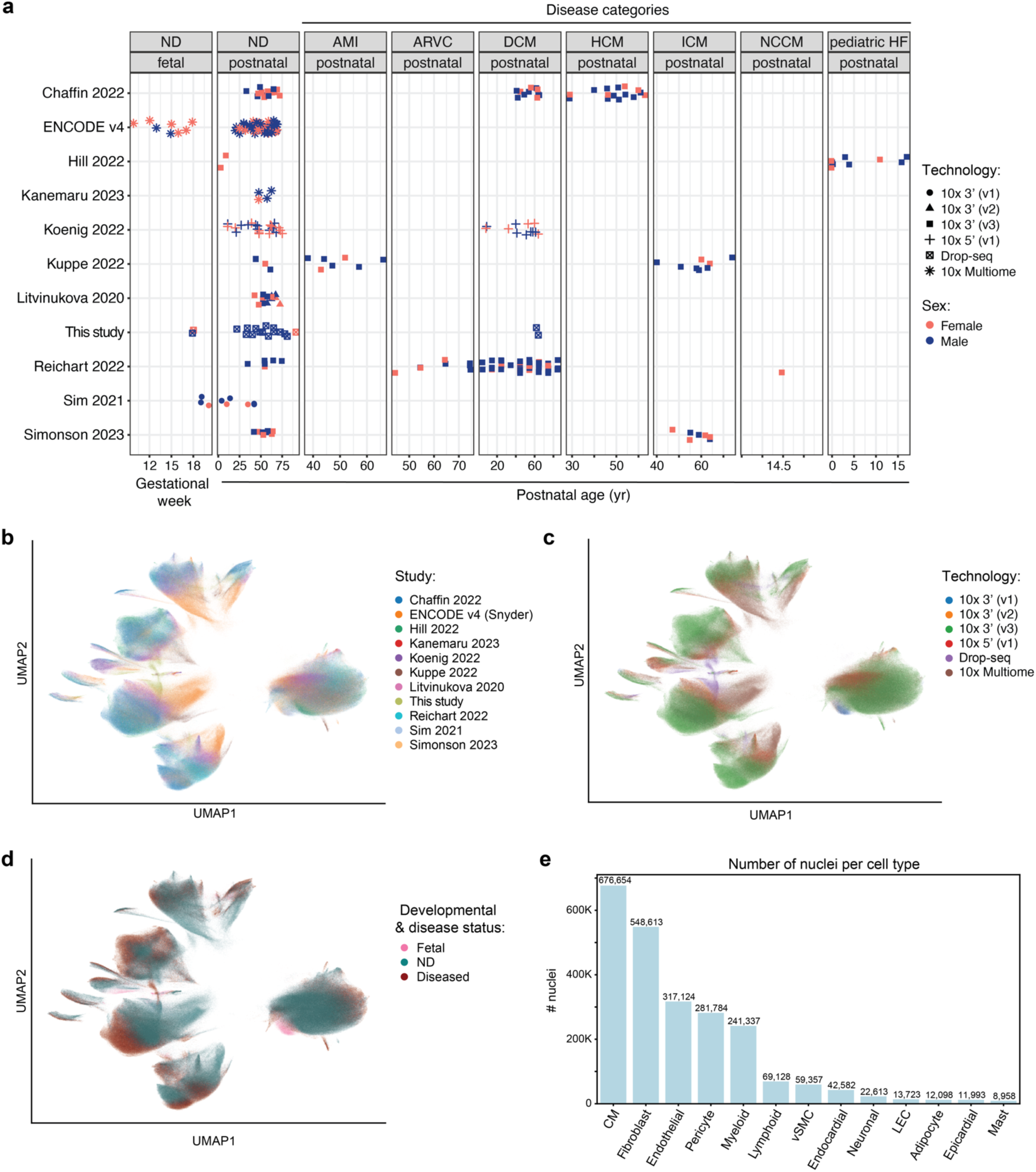
snRNA-seq additional figures: **(a)** Donors divided by age status and disease class (not binarized but split according to the actual disease) **(b-d)** scVI-based UMAP embedding based on study, technology, and developmental + disease status, respectively. **(e)** Number of nuclei per cell type in snRNA-seq aggregated dataset.

**Supplemental Figure 3.**
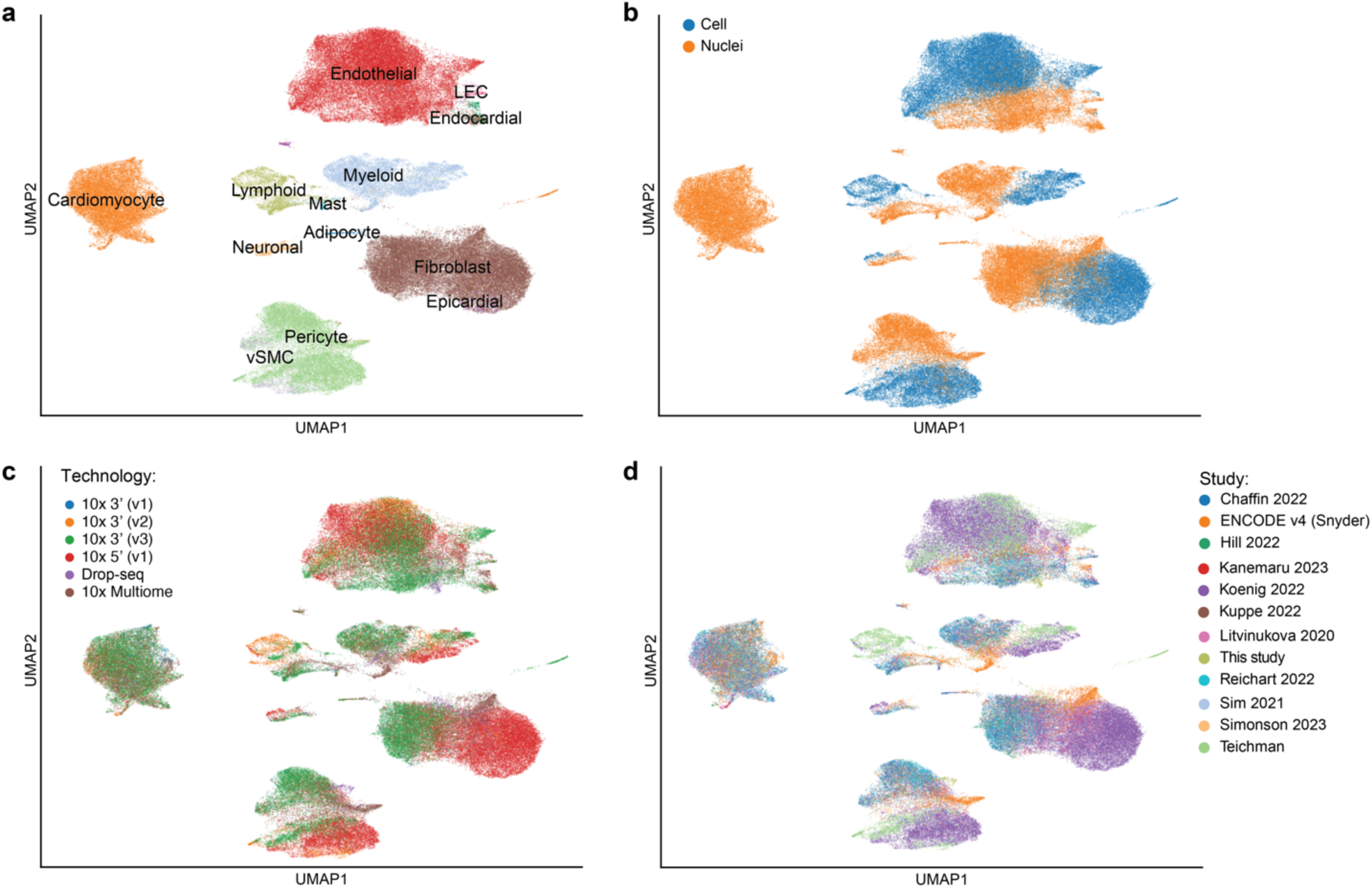
single cell RNA-seq vs. single nucleus RNA-seq: 50,000 scRNA-seq cells from the only two datasets (Koenig et al. 2022 and Kanemaru et al. 2023) that performed single-cell were combined with 50,000 snRNA-seq randomly subsampled from the aggregated snRNA-seq dataset (comprised of ∼2.3 million nuclei) were combined. Integration with scVI was attempted, with cell_vs_nuclei as a major batch effect in the model. Integration is only partially effective, with cells and nuclei not fully mixing. UMAP plots by **(a)** original cell type annotation, **(b)** cell or nuclei, **(c)** technology, and **(d)** study are displayed.

**Supplemental Figure 4.**
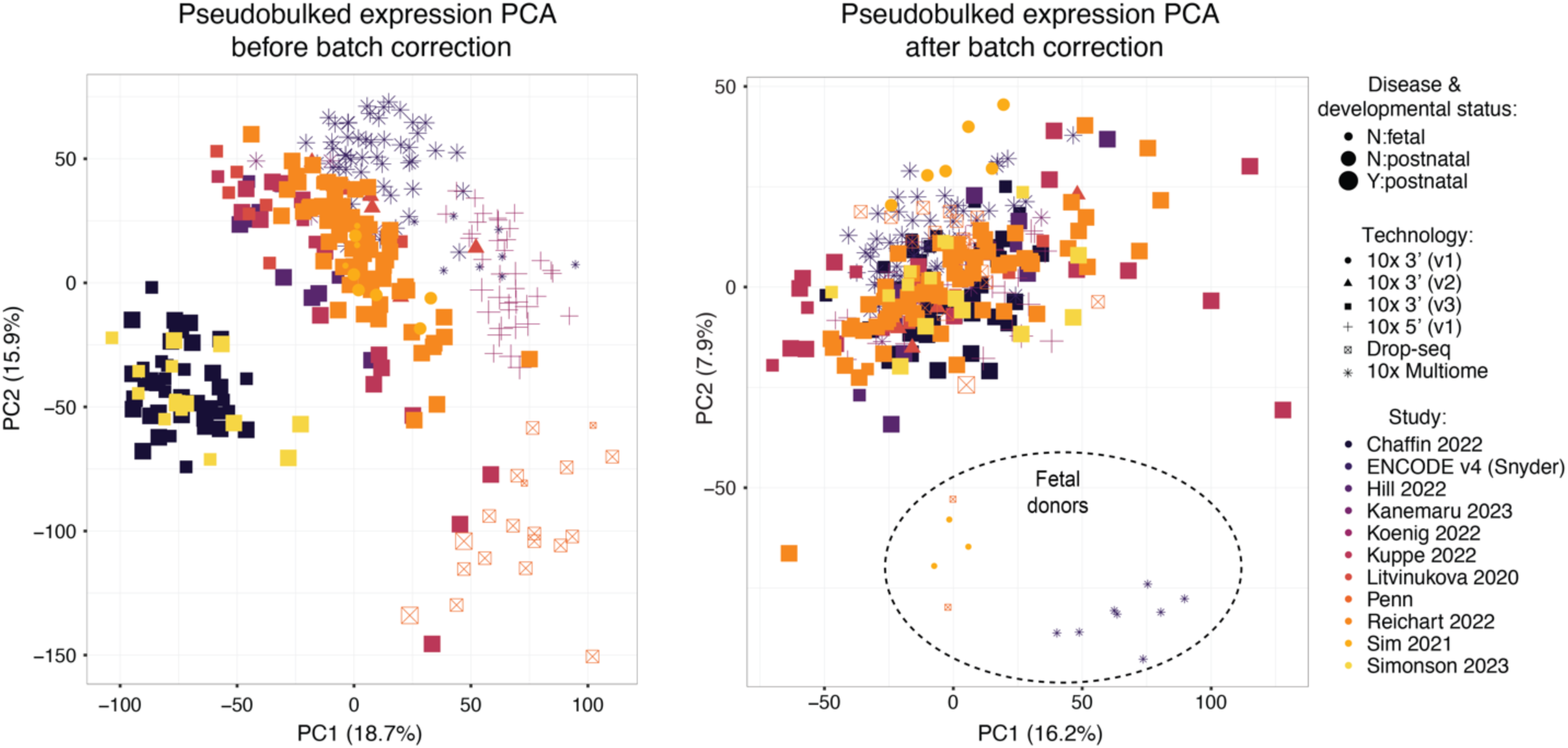
Batch effects in snRNA-seq count matrix: PCA of fully pseudobulked variance stabilized counts before and after batch correction (tech_plus_study). Left: Without removing tech_plus_study, the samples clearly cluster by study of origin. Right: By regressing out the effect of tech_plus_study (using limma), we observe better biological separation of samples, with PC2 separating fetal and postnatal donors.

**Supplemental Figure 5.**
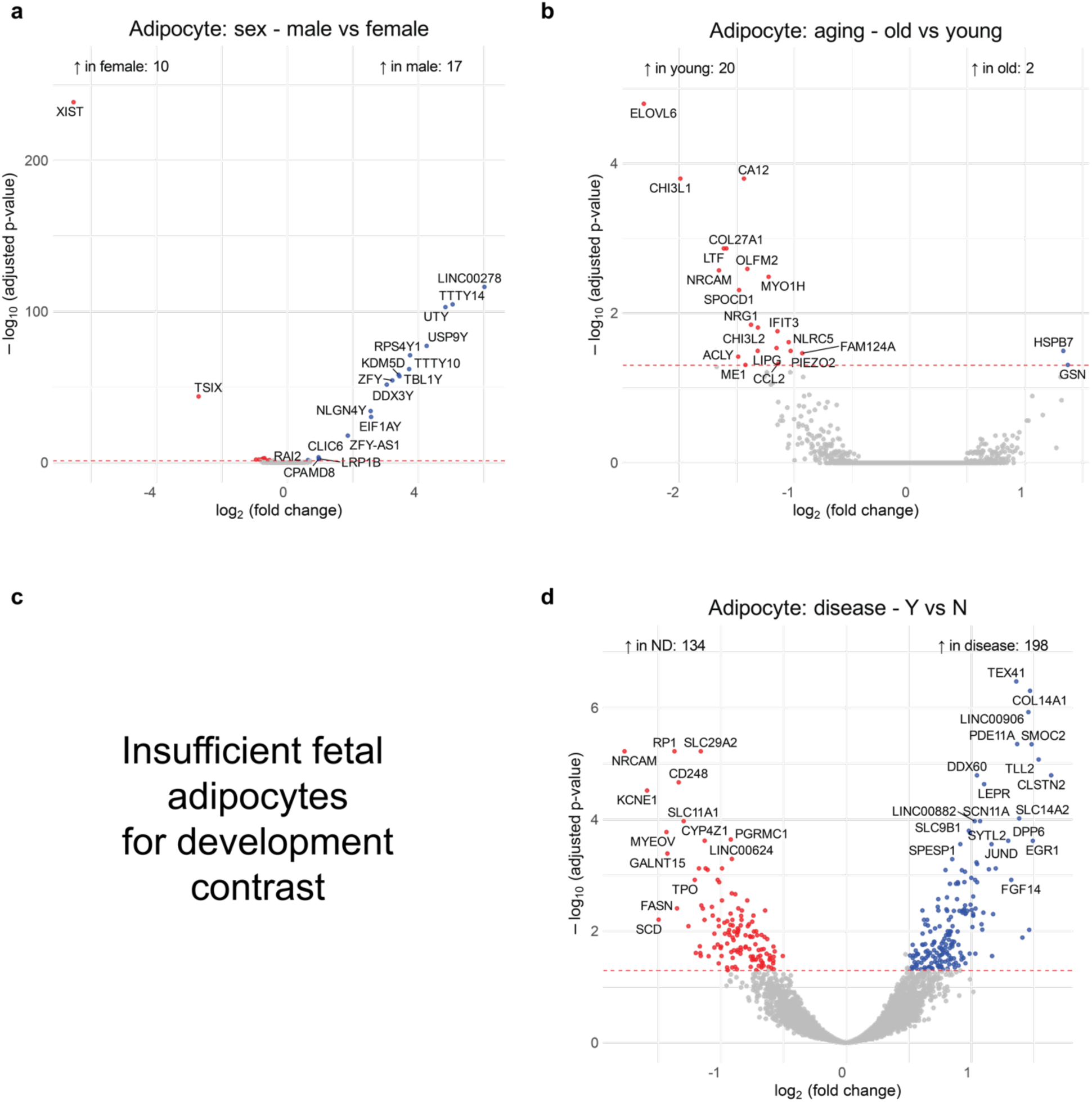
snRNA-seq Adipocyte volcano plots: Using DESeq2, donor-pseudobulked snRNA-seq Adipocyte volcano plots for the **(a)** sex, **(b)** aging, **(c)** development and **(d)** disease contrasts. As there are almost no fetal adipocytes, we did not perform development contrast for this cell type. Significance thresholds: |log2 fold change| > 0.5 and Benjamini-Hochberg adjusted p-value < 0.05.

**Supplemental Figure 6.**
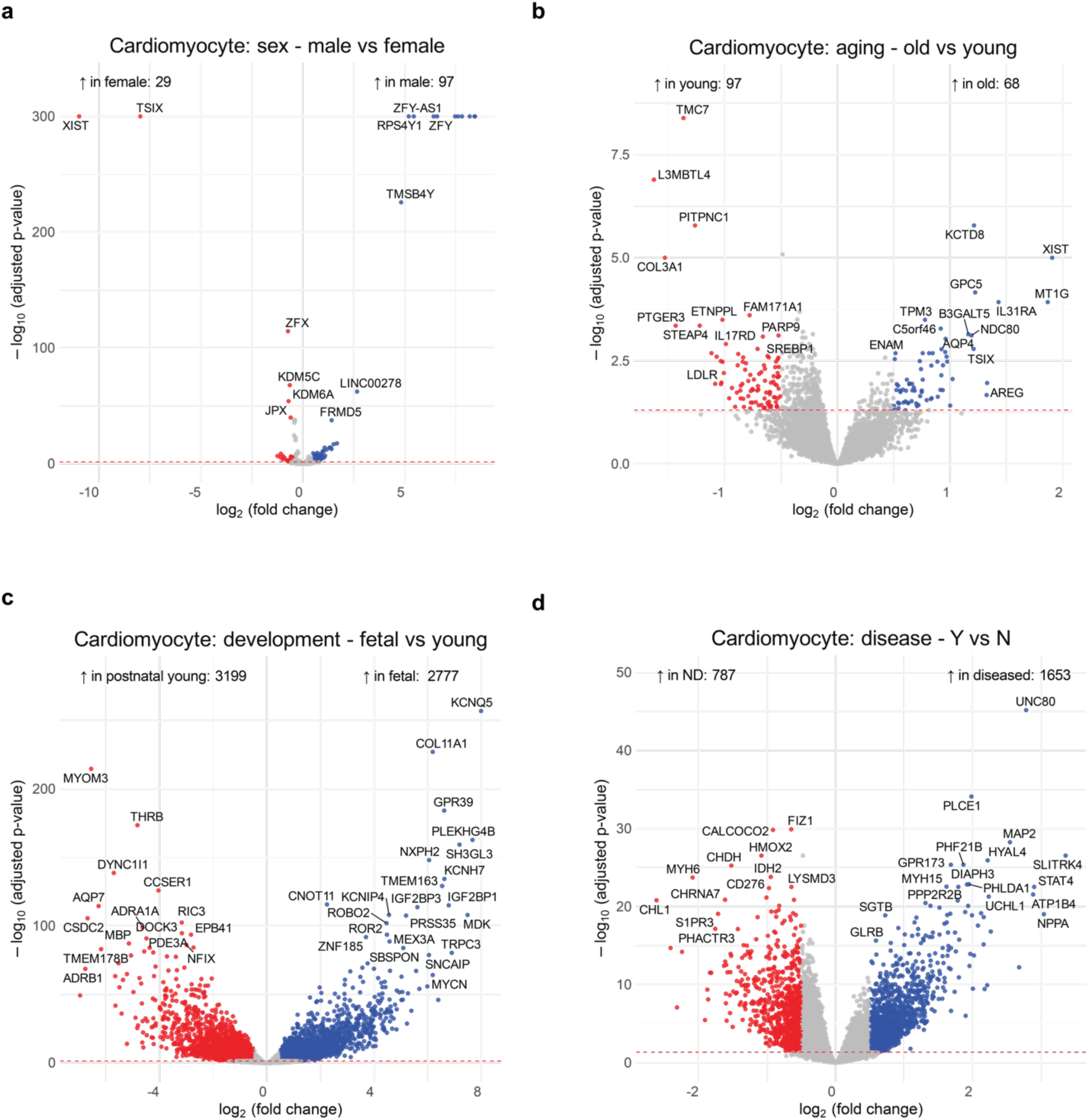
snRNA-seq Cardiomyocyte volcano plots: Using DESeq2, donor-pseudobulked snRNA-seq Cardiomyocyte volcano plots for the **(a)** sex, **(b)** aging, **(c)** development and **(d)** disease contrasts. Significance thresholds: |log2 fold change| > 0.5 and Benjamini-Hochberg adjusted p-value < 0.05.

**Supplemental Figure 7.**
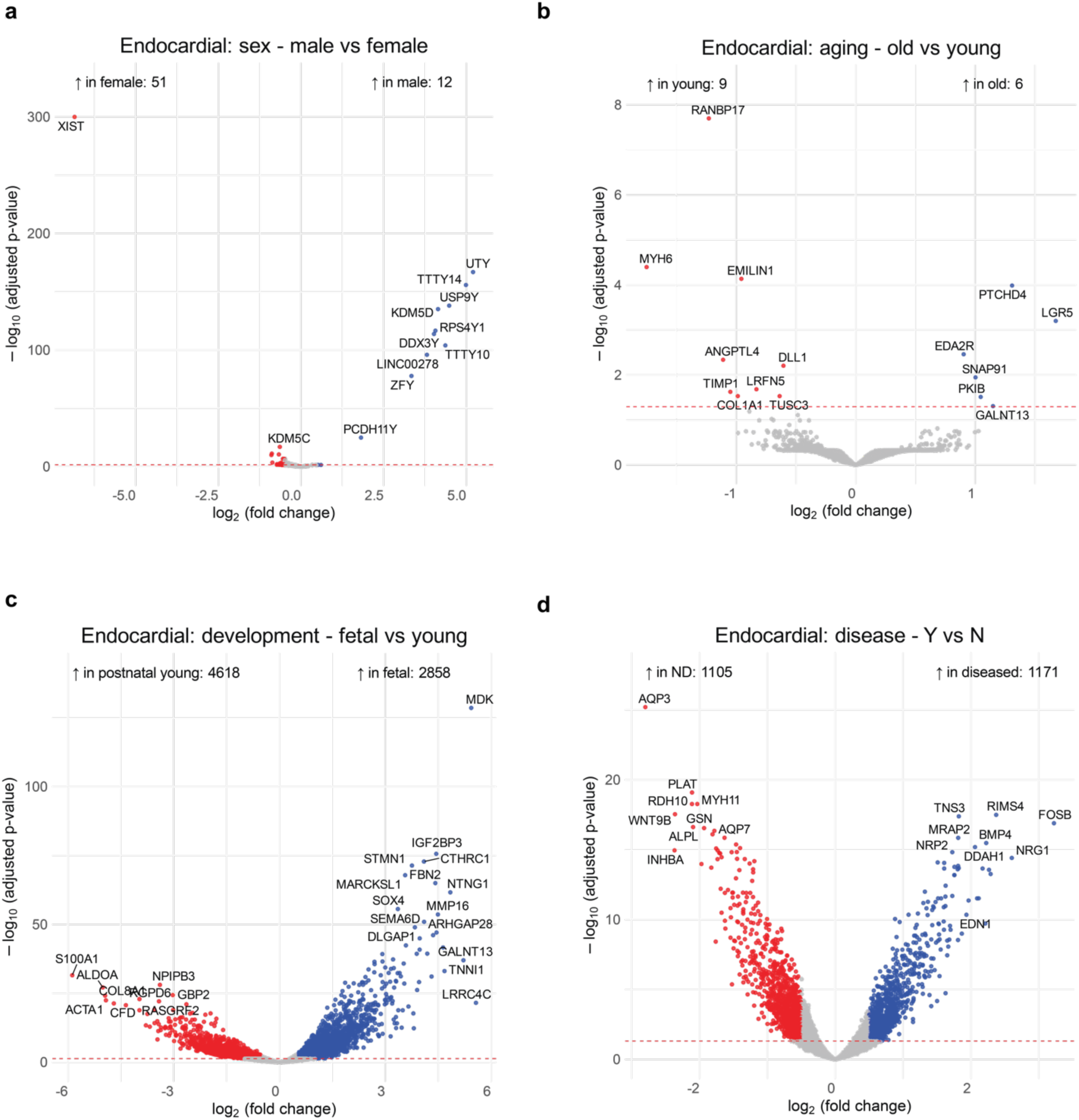
snRNA-seq Endocardial cell volcano plots: Using DESeq2, donor-pseudobulked snRNA-seq Endocardial volcano plots for the **(a)** sex, **(b)** aging, **(c)** development and **(d)** disease contrasts. Significance thresholds: |log2 fold change| > 0.5 and Benjamini-Hochberg adjusted p-value < 0.05.

**Supplemental Figure 8.**
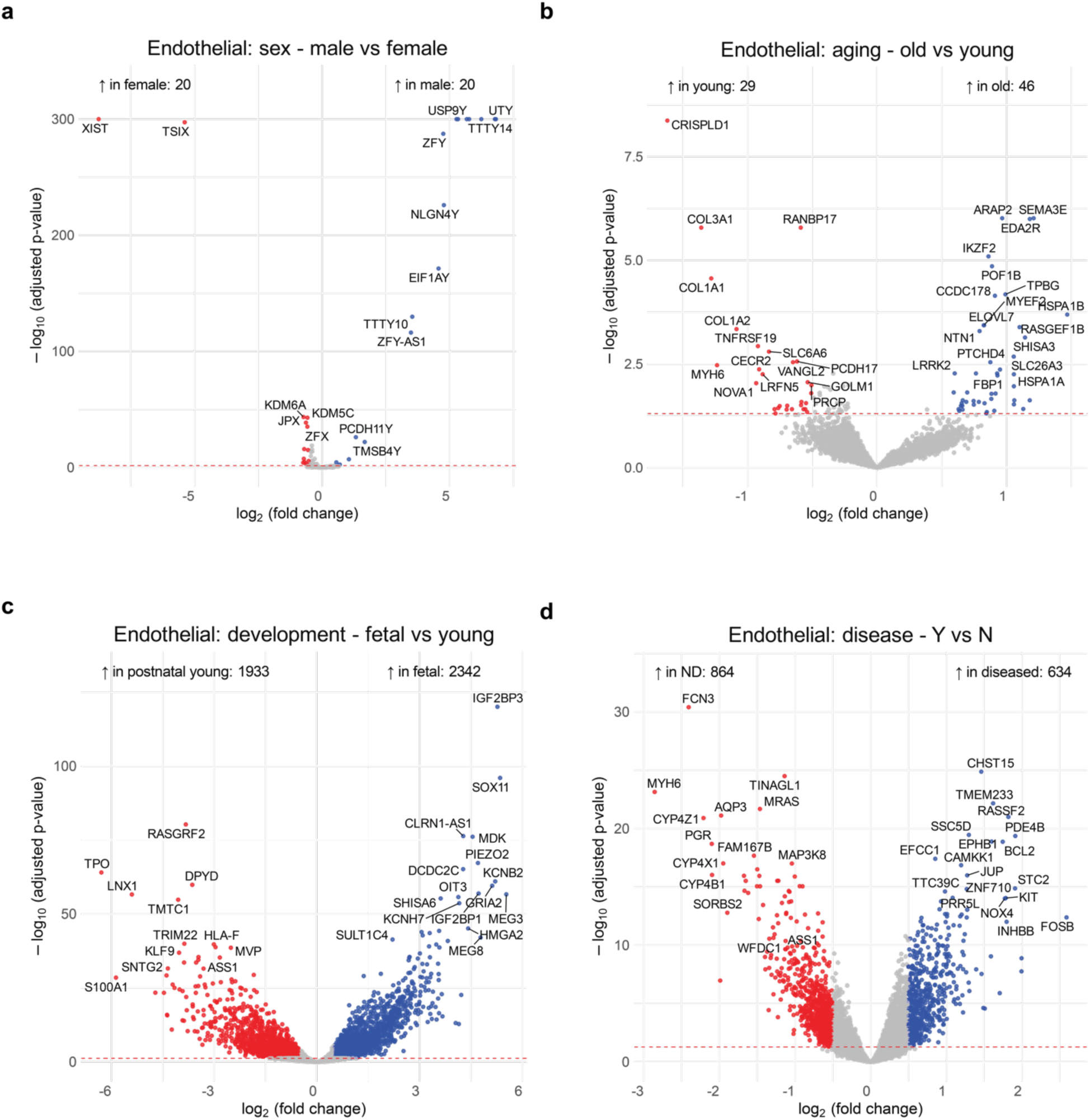
snRNA-seq Endothelial cell volcano plots: Using DESeq2, donor-pseudobulked snRNA-seq Endothelial volcano plots for the **(a)** sex, **(b)** aging, **(c)** development and **(d)** disease contrasts. Significance thresholds: |log2 fold change| > 0.5 and Benjamini-Hochberg adjusted p-value < 0.05.

**Supplemental Figure 9.**
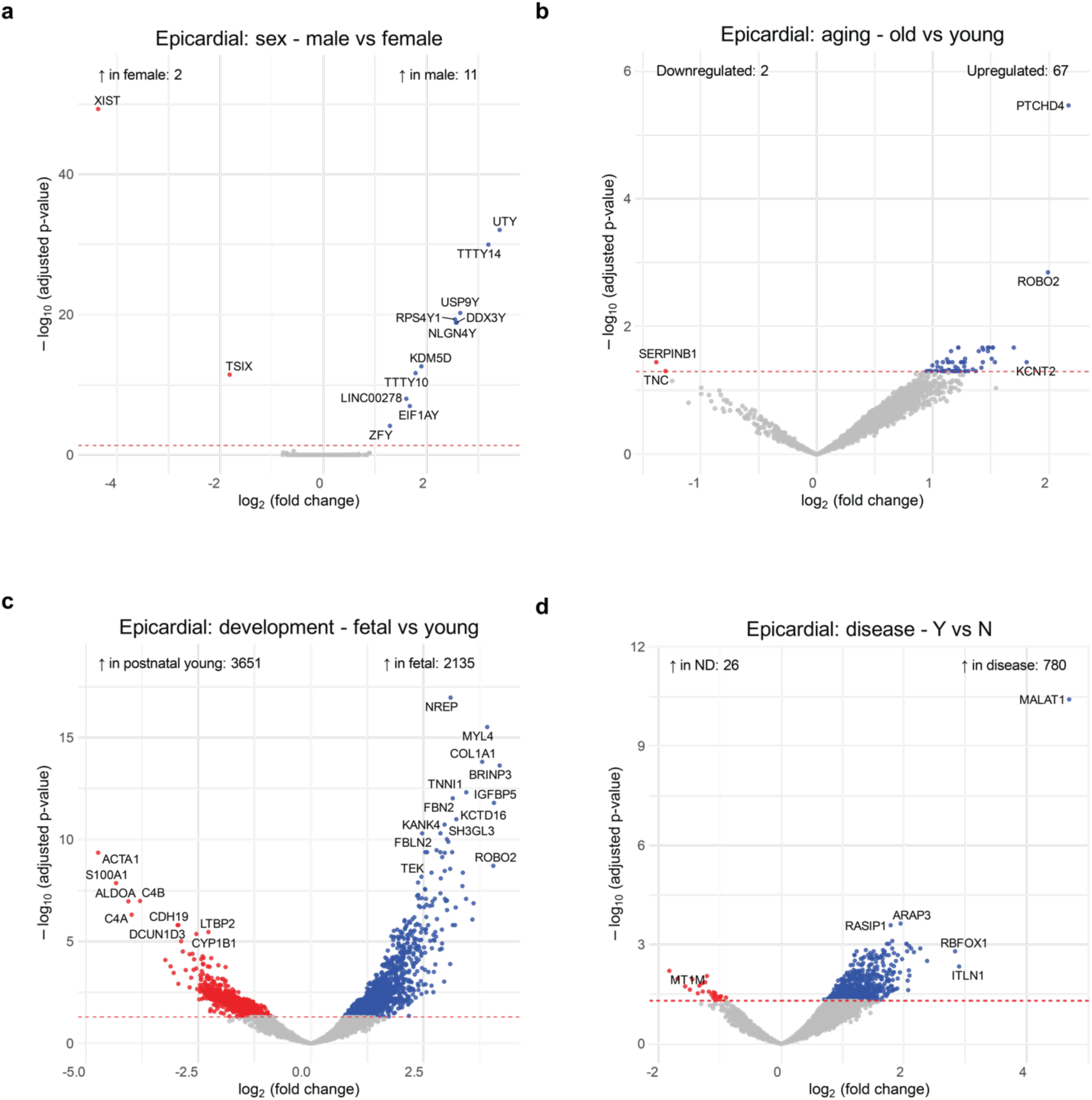
snRNA-seq Epicardial cell volcano plots: Using DESeq2, donor-pseudobulked snRNA-seq Epicardial volcano plots for the **(a)** sex, **(b)** aging, **(c)** development and **(d)** disease contrasts. Significance thresholds: |log2 fold change| > 0.5 and Benjamini-Hochberg adjusted p-value < 0.05.

**Supplemental Figure 10.**
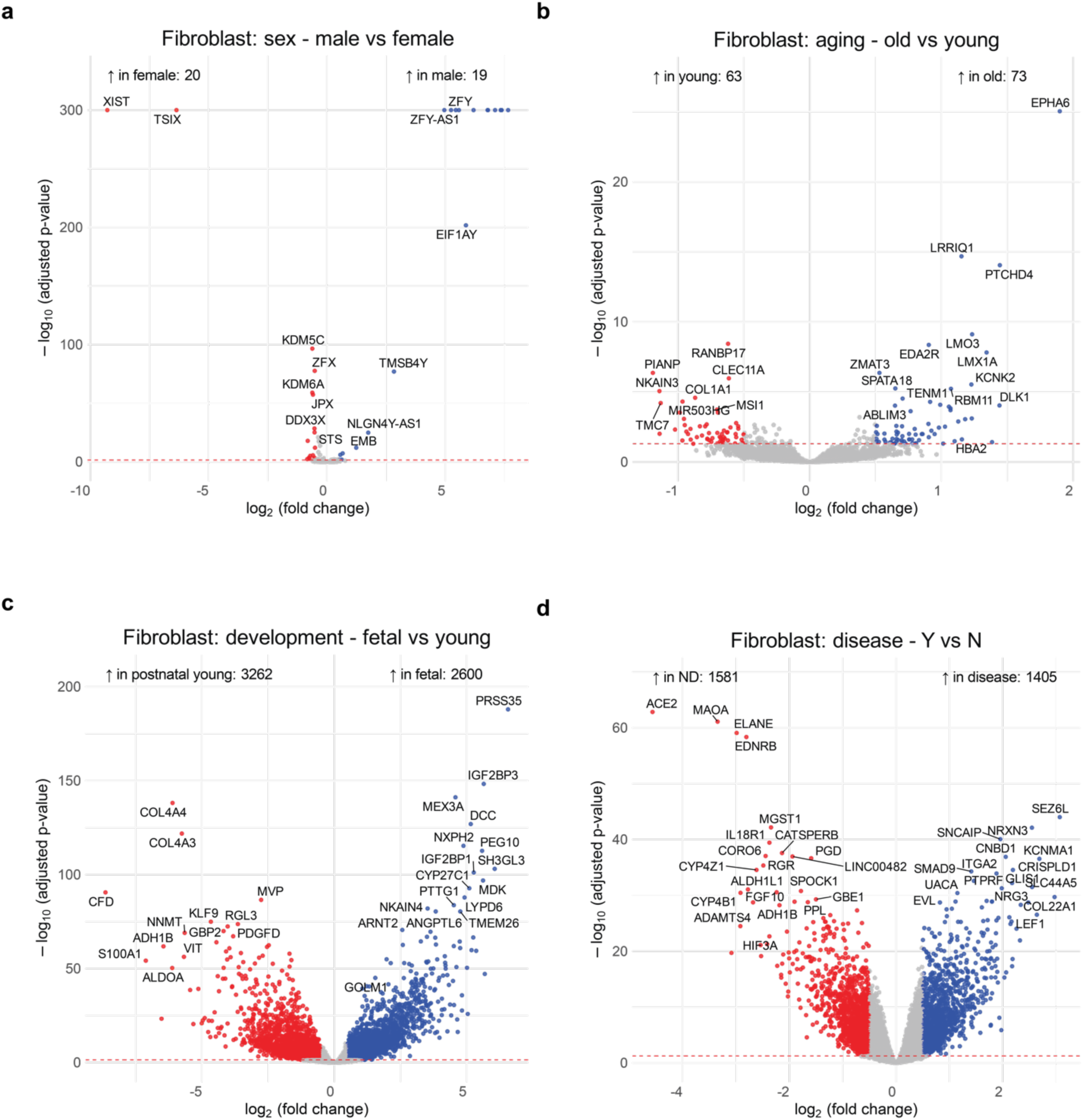
snRNA-seq Fibroblast volcano plots: Using DESeq2, donor-pseudobulked snRNA-seq Fibroblast volcano plots for the **(a)** sex, **(b)** aging, **(c)** development and **(d)** disease contrasts. Significance thresholds: |log2 fold change| > 0.5 and Benjamini-Hochberg adjusted p-value < 0.05.

**Supplemental Figure 11.**
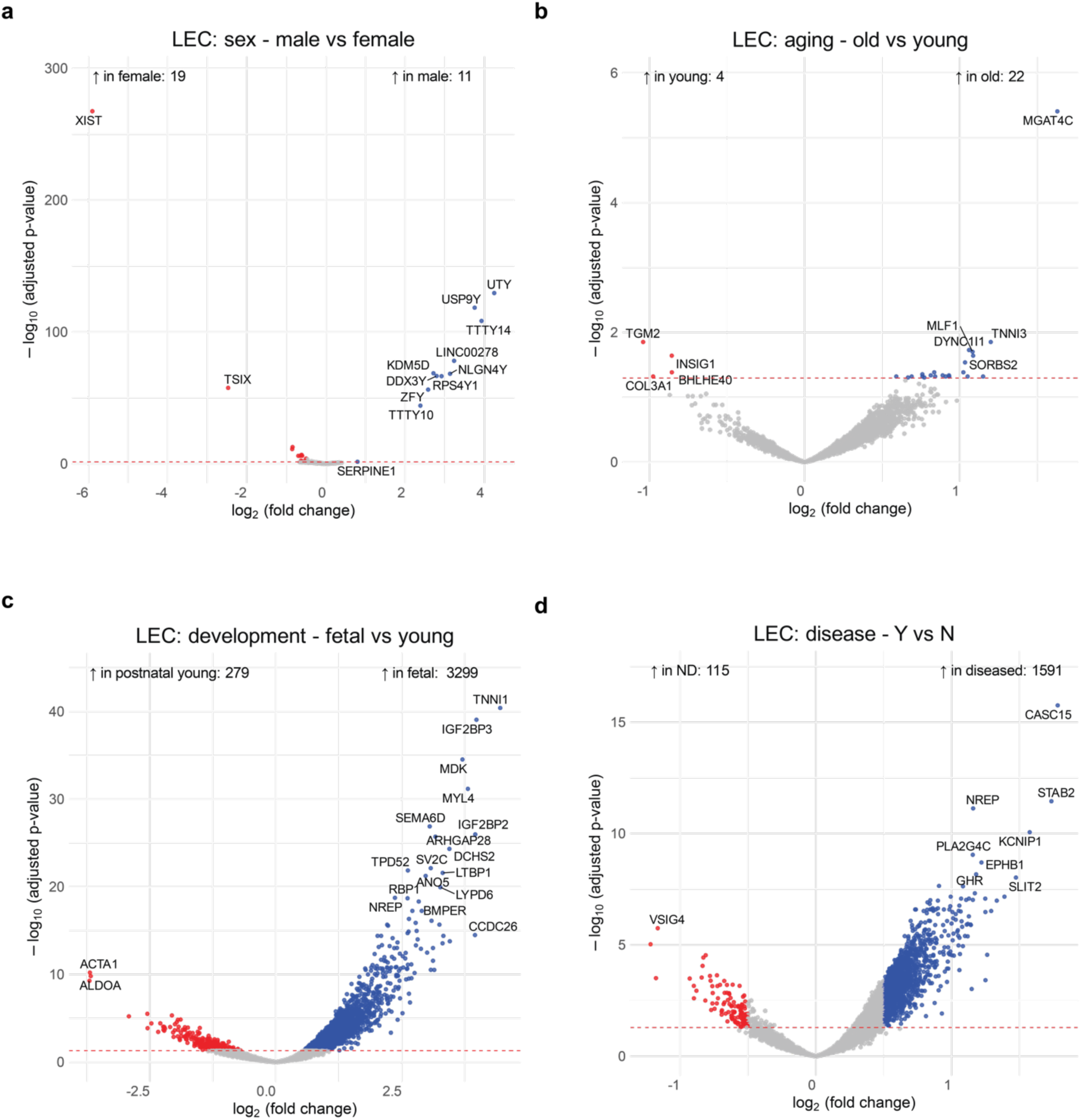
snRNA-seq LEC volcano plots: Using DESeq2, donor-pseudobulked snRNA-seq Lymphatic epithelial cell (LEC) volcano plots for the **(a)** sex, **(b)** aging, **(c)** development and **(d)** disease contrasts. Significance thresholds: |log2 fold change| > 0.5 and Benjamini-Hochberg adjusted p-value < 0.05.

**Supplemental Figure 12.**
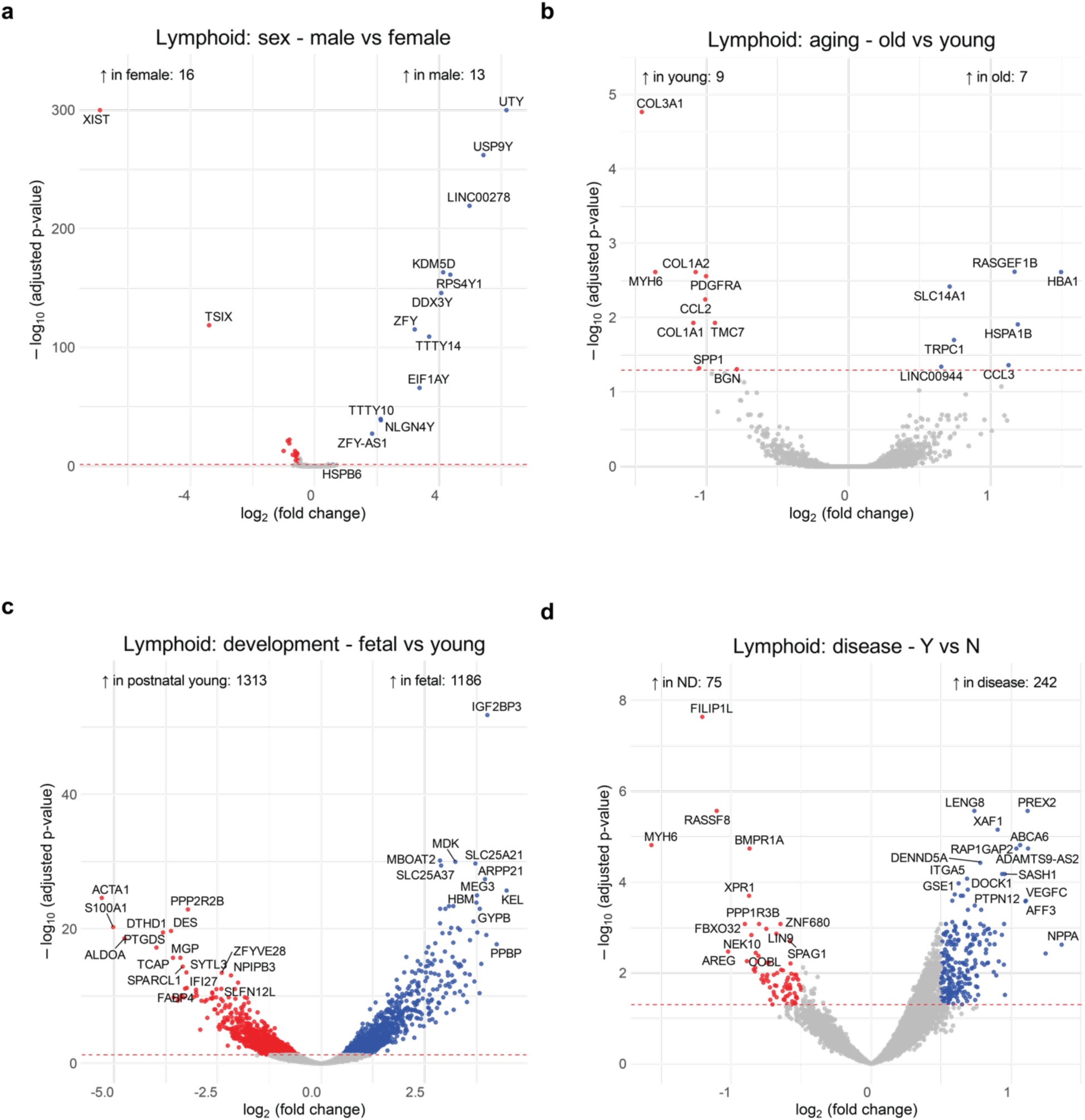
snRNA-seq Lymphoid cell volcano plots: Using DESeq2, donor-pseudobulked snRNA-seq Lymphoid volcano plots for the **(a)** sex, **(b)** aging, **(c)** development and **(d)** disease contrasts. Significance thresholds: |log2 fold change| > 0.5 and Benjamini-Hochberg adjusted p-value < 0.05.

**Supplemental Figure 13.**
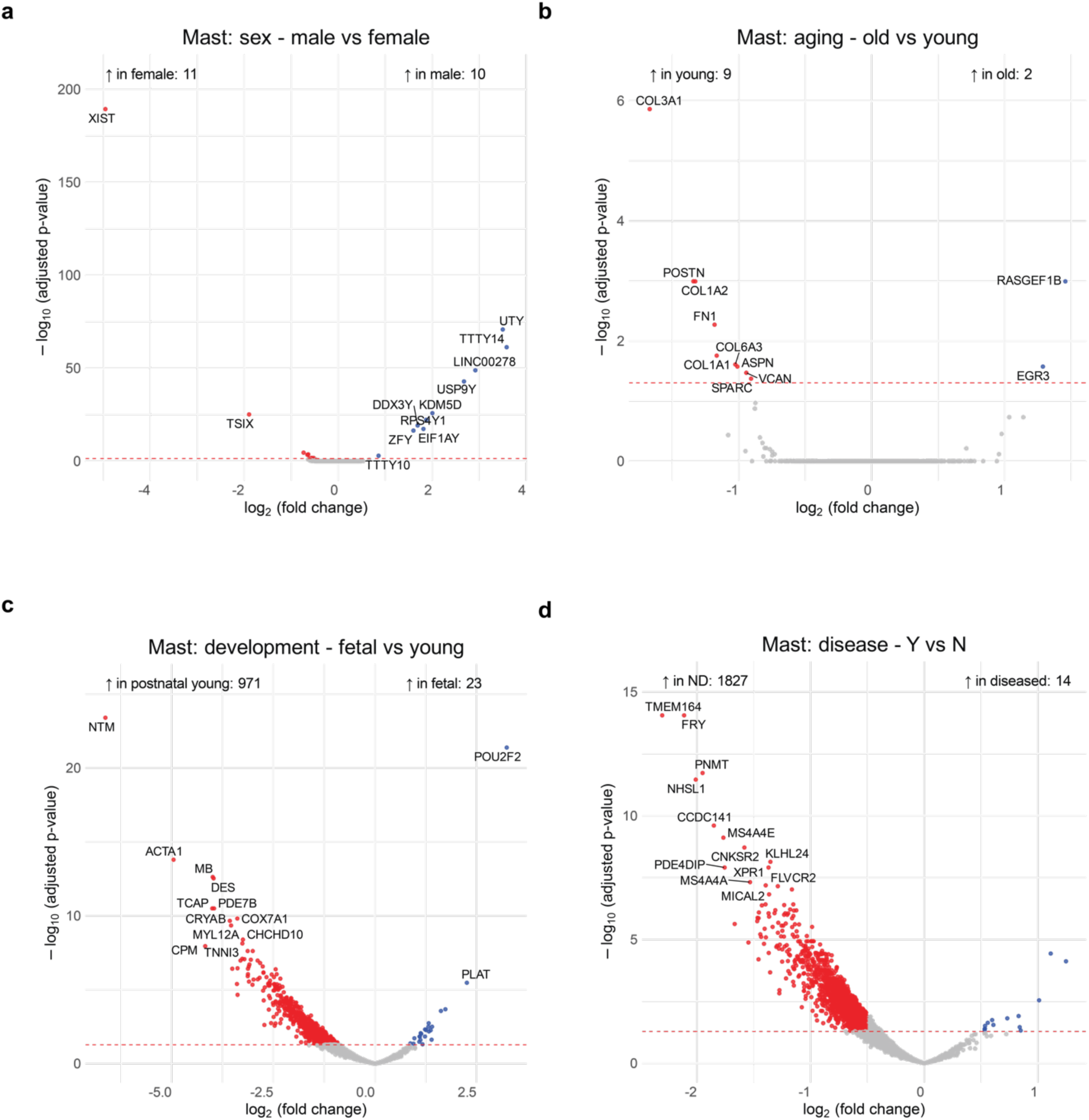
snRNA-seq Mast cell volcano plots: Using DESeq2, donor-pseudobulked snRNA-seq Mast volcano plots for the **(a)** sex, **(b)** aging, **(c)** development and **(d)** disease contrasts. Significance thresholds: |log2 fold change| > 0.5 and Benjamini-Hochberg adjusted p-value < 0.05.

**Supplemental Figure 14.**
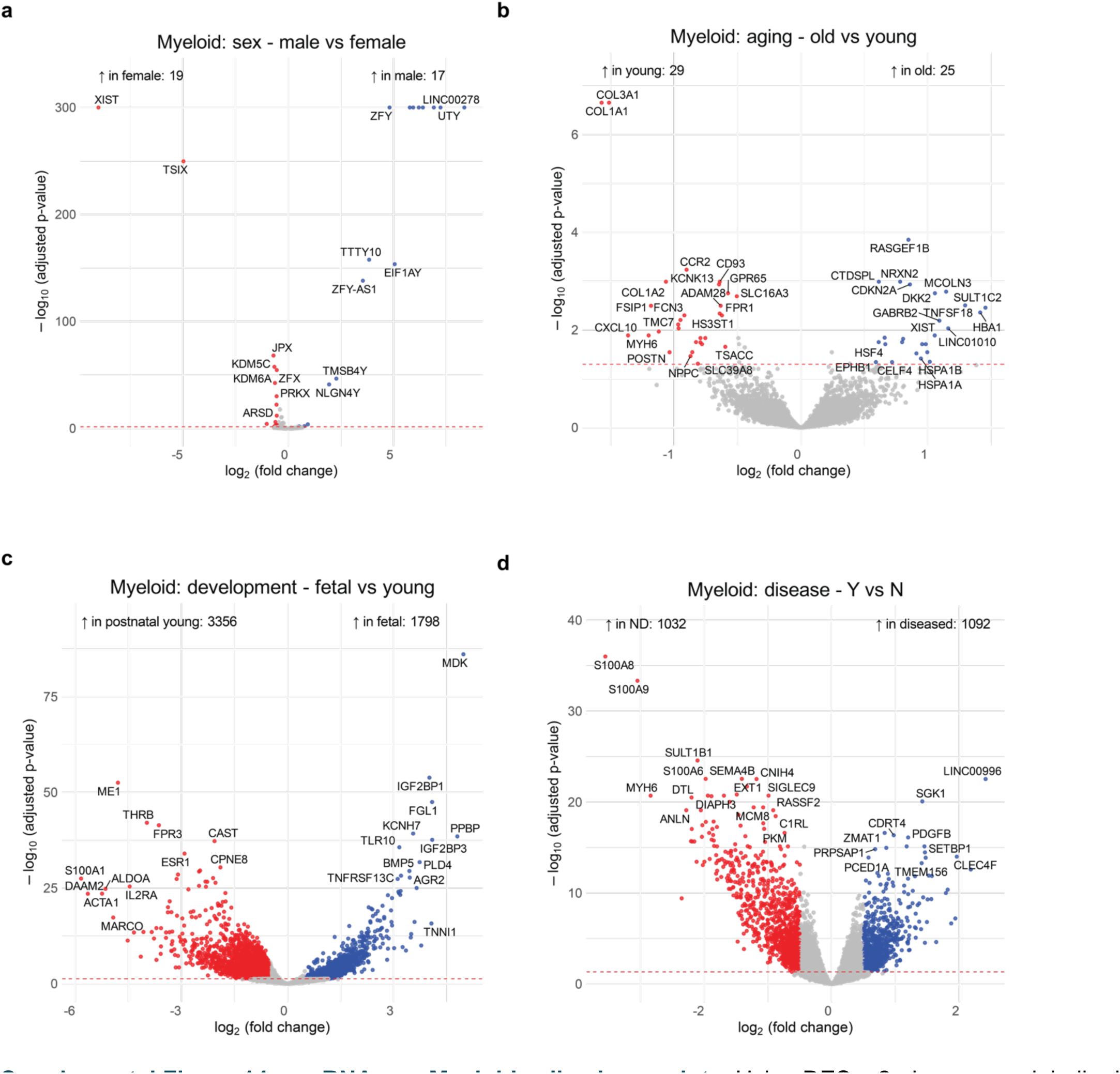
snRNA-seq Myeloid cell volcano plots: Using DESeq2, donor-pseudobulked snRNA-seq Myeloid volcano plots for the **(a)** sex, **(b)** aging, **(c)** development and **(d)** disease contrasts. Significance thresholds: |log2 fold change| > 0.5 and Benjamini-Hochberg adjusted p-value < 0.05.

**Supplemental Figure 15.**
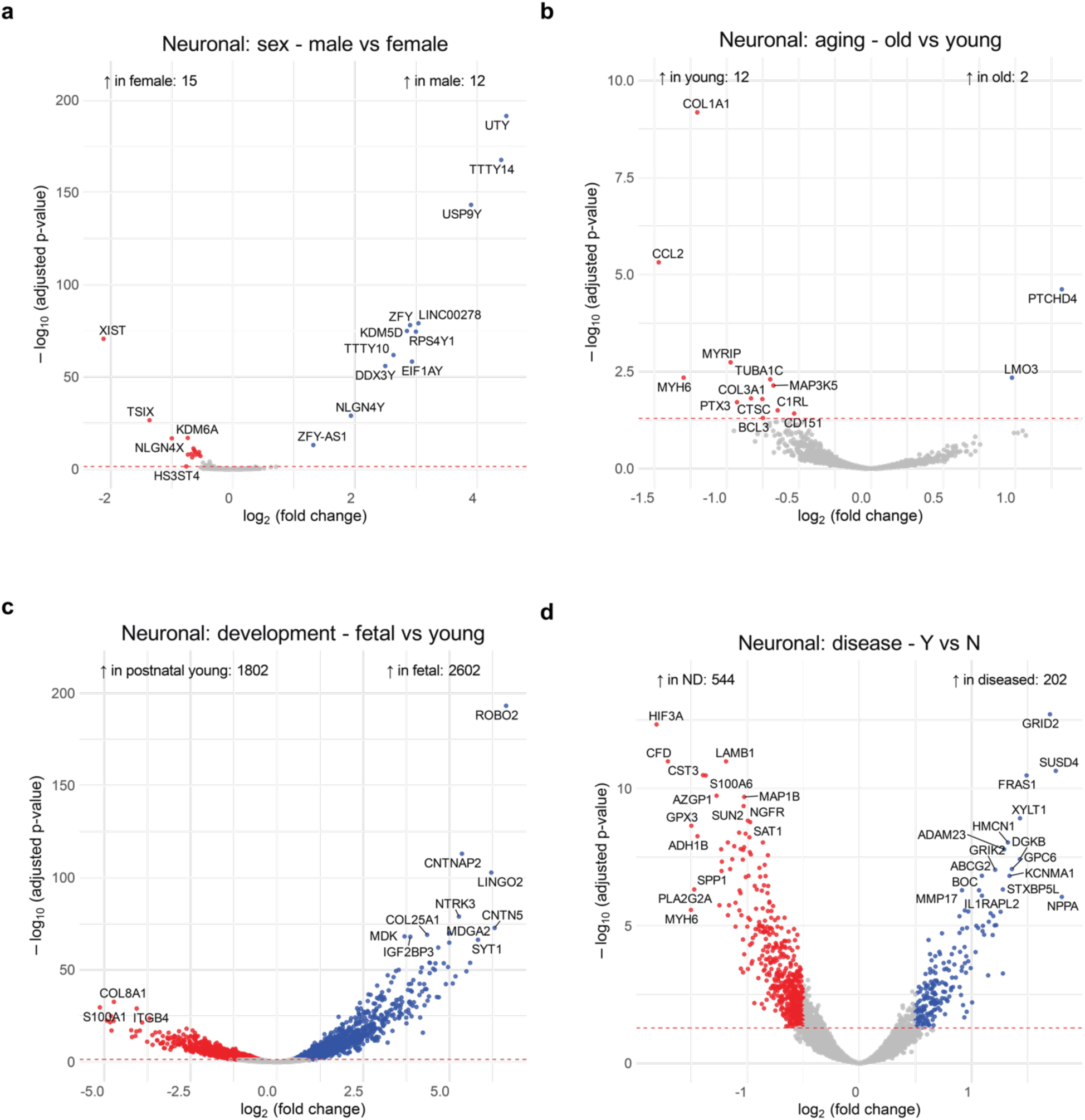
snRNA-seq Neuronal cell volcano plots: Using DESeq2, donor-pseudobulked snRNA-seq Neuronal volcano plots for the **(a)** sex, **(b)** aging, **(c)** development and **(d)** disease contrasts. Significance thresholds: |log2 fold change| > 0.5 and Benjamini-Hochberg adjusted p-value < 0.05.

**Supplemental Figure 16.**
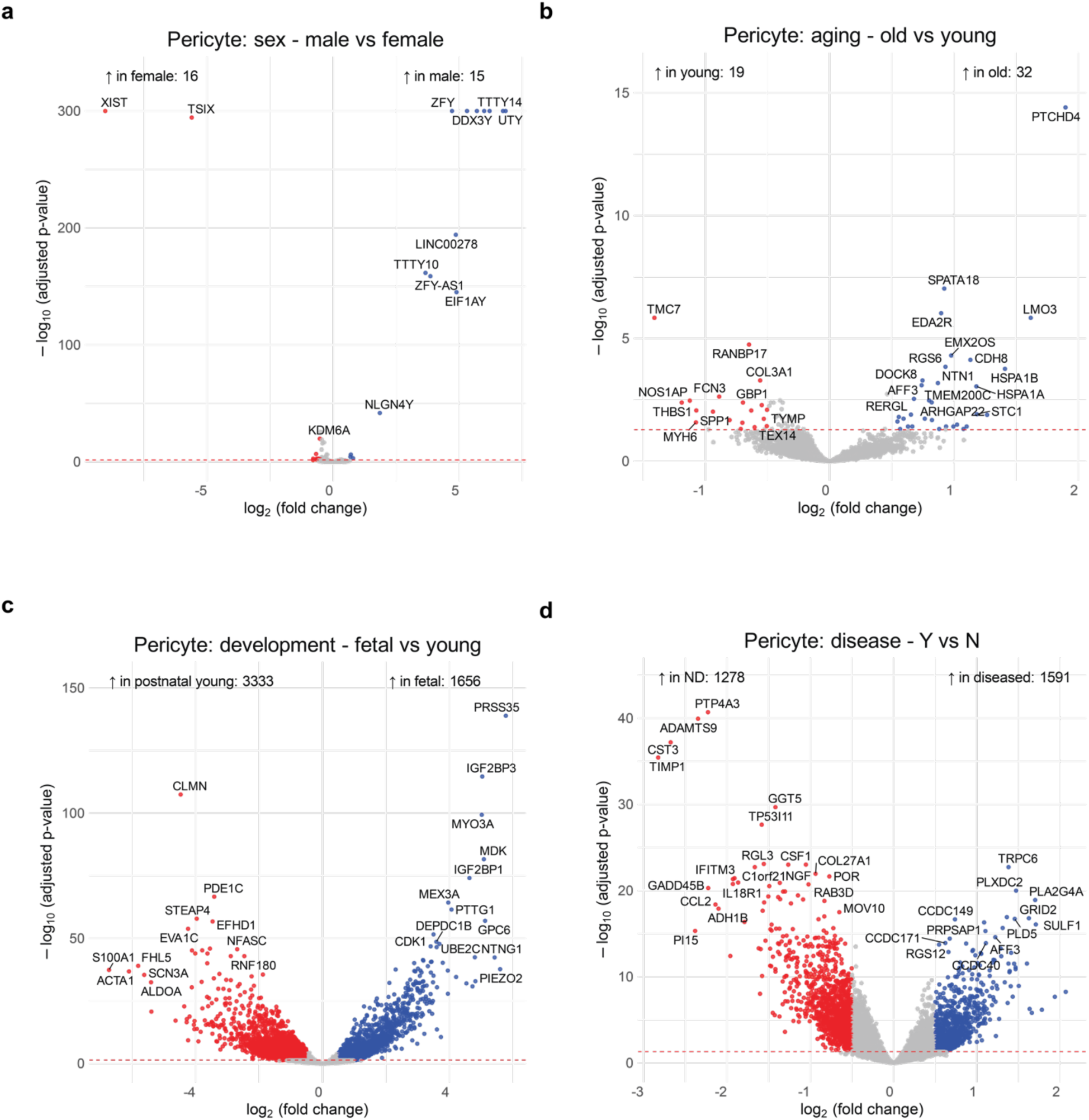
snRNA-seq Pericyte volcano plots: Using DESeq2, donor-pseudobulked snRNA-seq Pericyte volcano plots for the **(a)** sex, **(b)** aging, **(c)** development and **(d)** disease contrasts. Significance thresholds: |log2 fold change| > 0.5 and Benjamini-Hochberg adjusted p-value < 0.05.

**Supplemental Figure 17.**
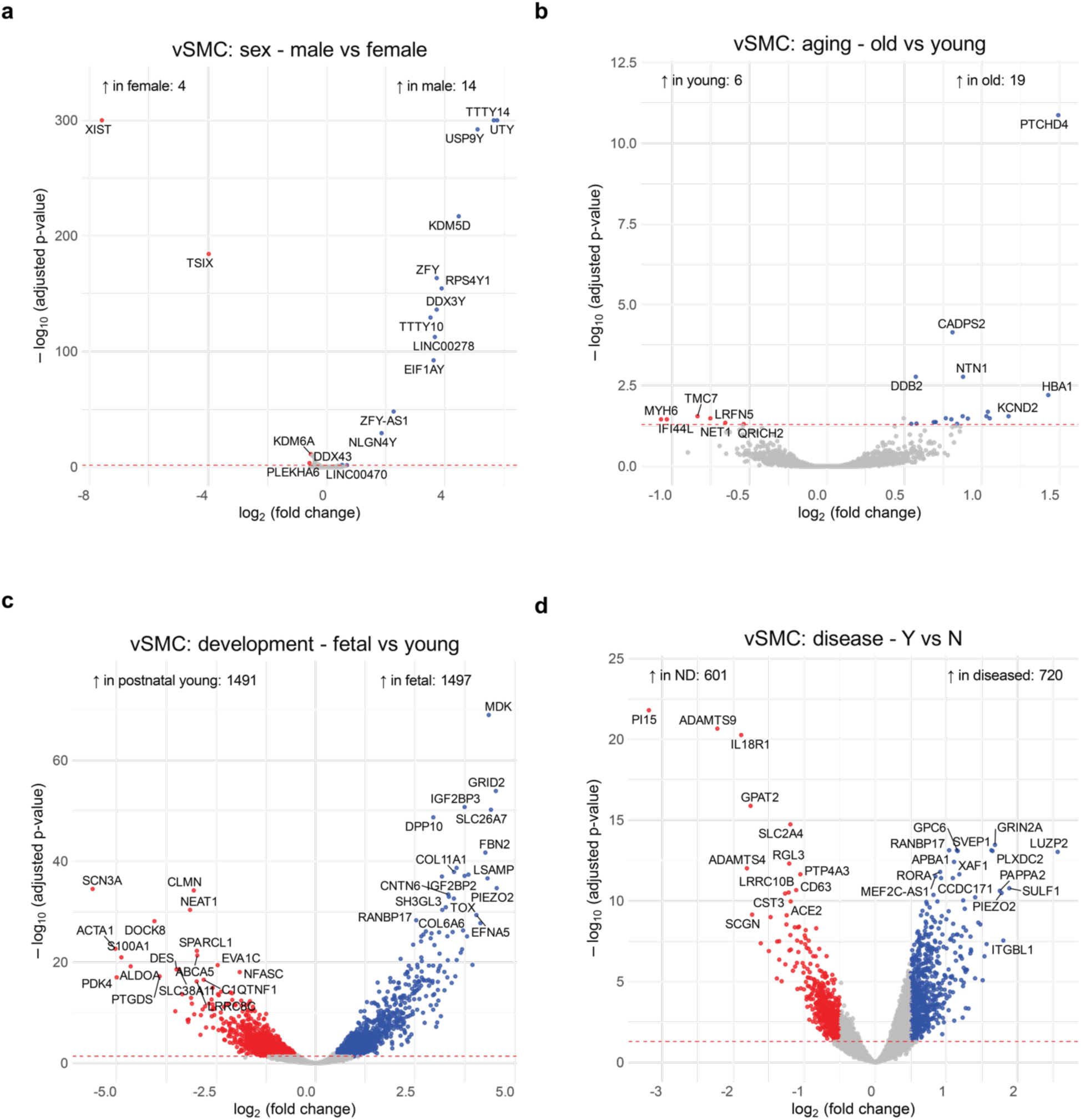
snRNA-seq vSMC volcano plots: Using DESeq2, donor-pseudobulked snRNA-seq vSMC volcano plots for the **(a)** sex, **(b)** aging, **(c)** development and **(d)** disease contrasts. Significance thresholds: |log2 fold change| > 0.5 and Benjamini-Hochberg adjusted p-value < 0.05.

**Supplemental Figure 18.**
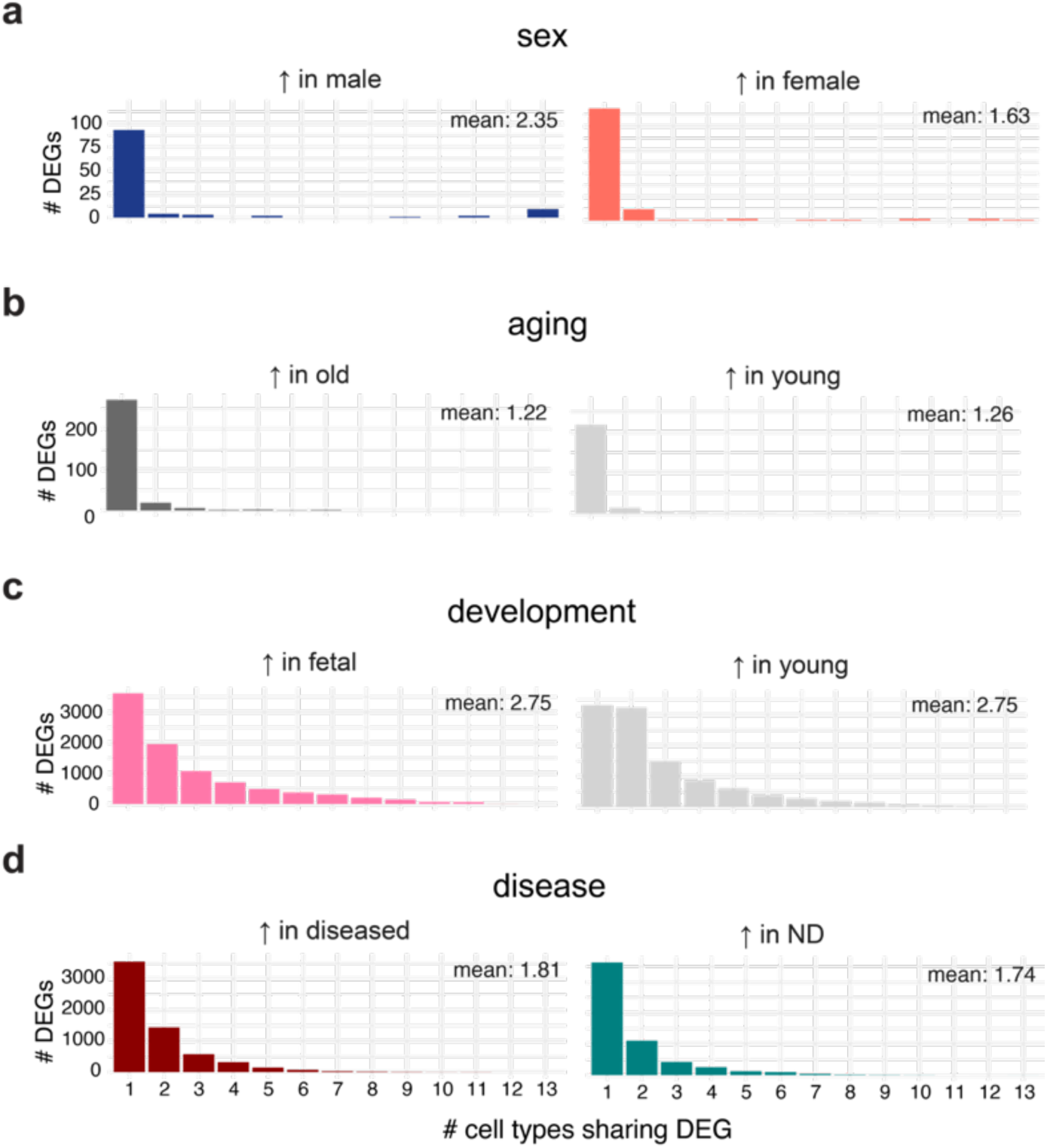
Number of shared DEGs across cell types: To determine how similar DEGs are across cell types, we computed for each DEG the number of cell types for which it is found as a DEG. Number of DEGs shared across cell types by **(a)** sex, **(b)** aging, **(c)** development, and **(d)** disease.

**Supplemental Figure 19.**
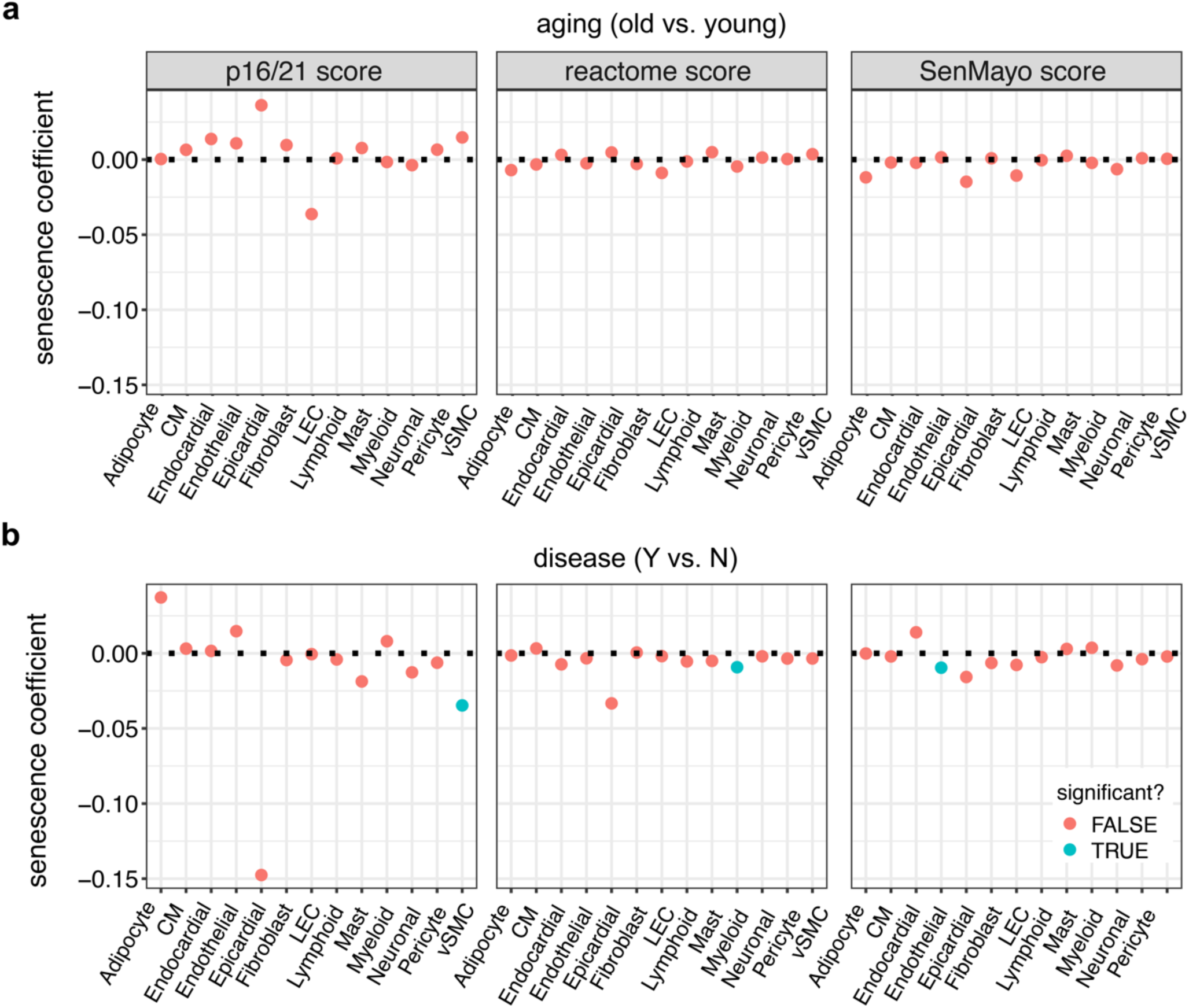
Senescence analysis: Using the scanpy score_genes() function, we computed the per-nucleus activity score using three different gene sets for senescence. The “p16/p21 score” involves only two genes classically associated with senescence, p16 (*CDKN1A*) and p21 (*CDKN2A*). The “reactome score” includes the genes in R-HSA-2559583. The SenMayo score includes the genes in the SenMayo gene set. The mean senescence score per donor was computed. For each senescence score, the value was modeled using a fixed effect model: score ∼ sex + age_group + disease_binary + tech_plus_study to obtain a coefficient and p-value. A positive coefficient corresponds to an increase in senescence with age or disease, while a negative coefficient corresponds to a decrease in senescence. Significant effects of either age group or disease are colored in green (Benjamini-Hochberg adjusted p-value < 0.05).

**Supplemental Figure 20.**
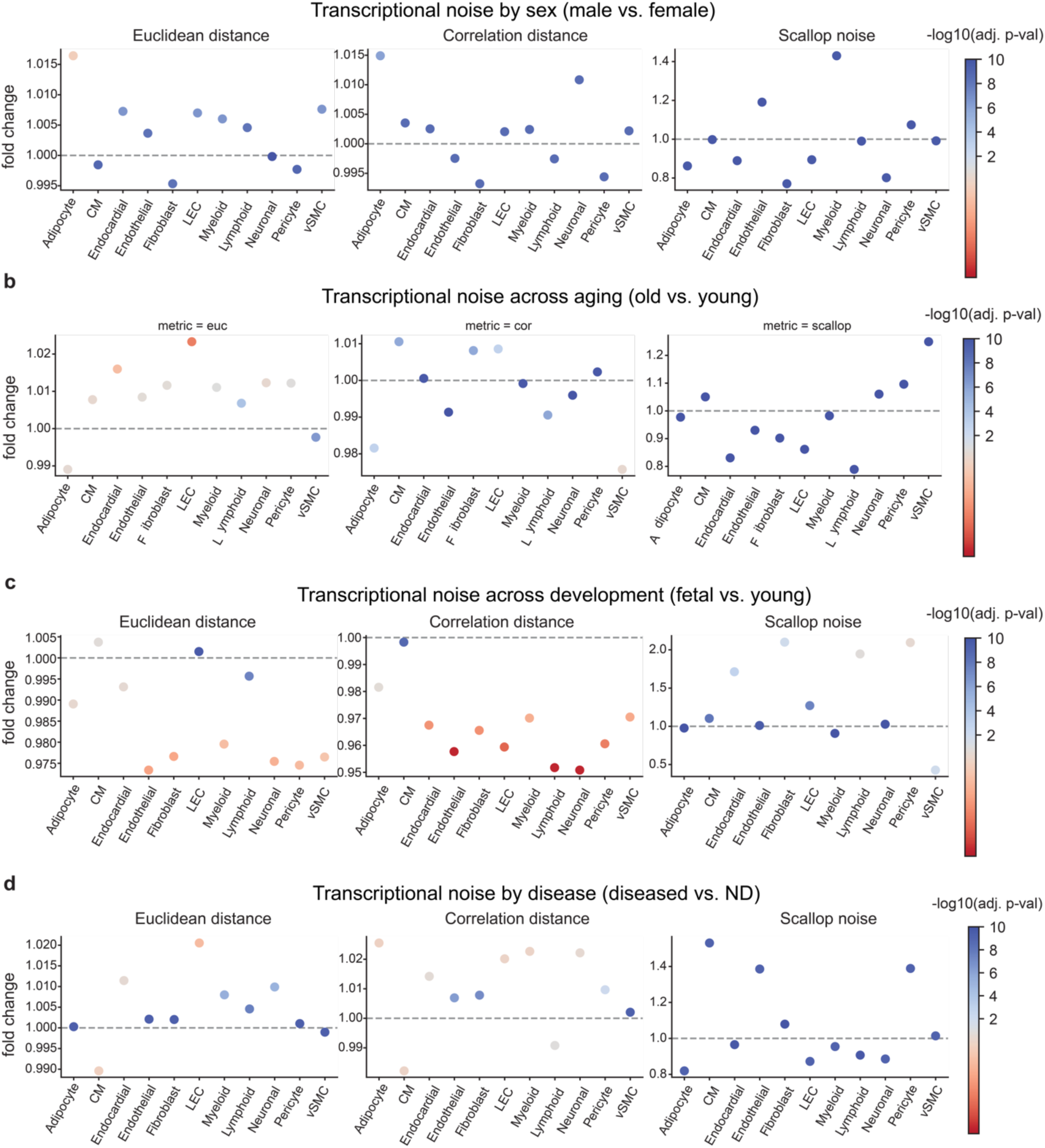
Transcriptional noise analysis: Transcriptional noise analysis using scallop. Three noise metrics (Euclidean distance, correlation distance, and scallop noise) were calculated for each cell type. The fold change of the mean between two groups in a contrast (e.g., male vs. female for the sex contrast) were calculated, with statistical significance determined using the Wilcoxon ranked sum test. (**a**) Transcriptional noise differences by sex: male vs. female (b) Transcriptional noise differences by aging: old vs. young **(c)** Transcriptional noise differences by development: fetal vs. postnatal young **(d)** Transcriptional noise differences by disease: diseased vs. ND.

**Supplemental Figure 21.**
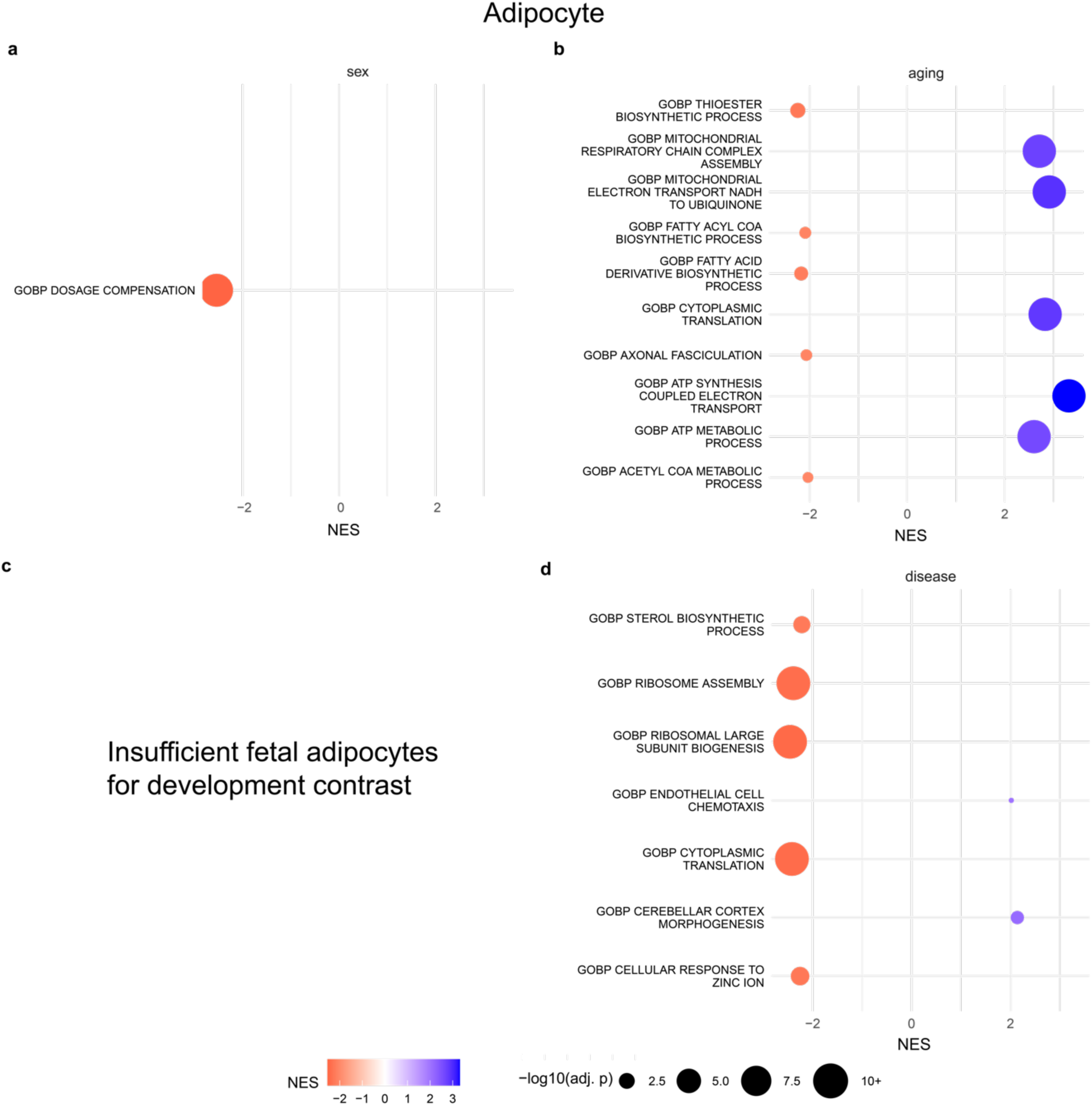
snRNA-seq Adipocyte gene set enrichment analysis: Adipocyte gene set enrichment analysis using the ordered list of genes for **(a)** sex, **(b)** aging, **(c)** development and **(d)** disease contrasts. NES = normalized enrichment score, so a negative NES for the sex contrast (male vs. female) would be interpreted as gene sets enriched in females and a positive NES would be interpreted as gene sets enriched in males. Only the top 5 positive and negative NES gene sets, ranked by Benjamini-Hochberg adjusted p-value are displayed.

**Supplemental Figure 22.**
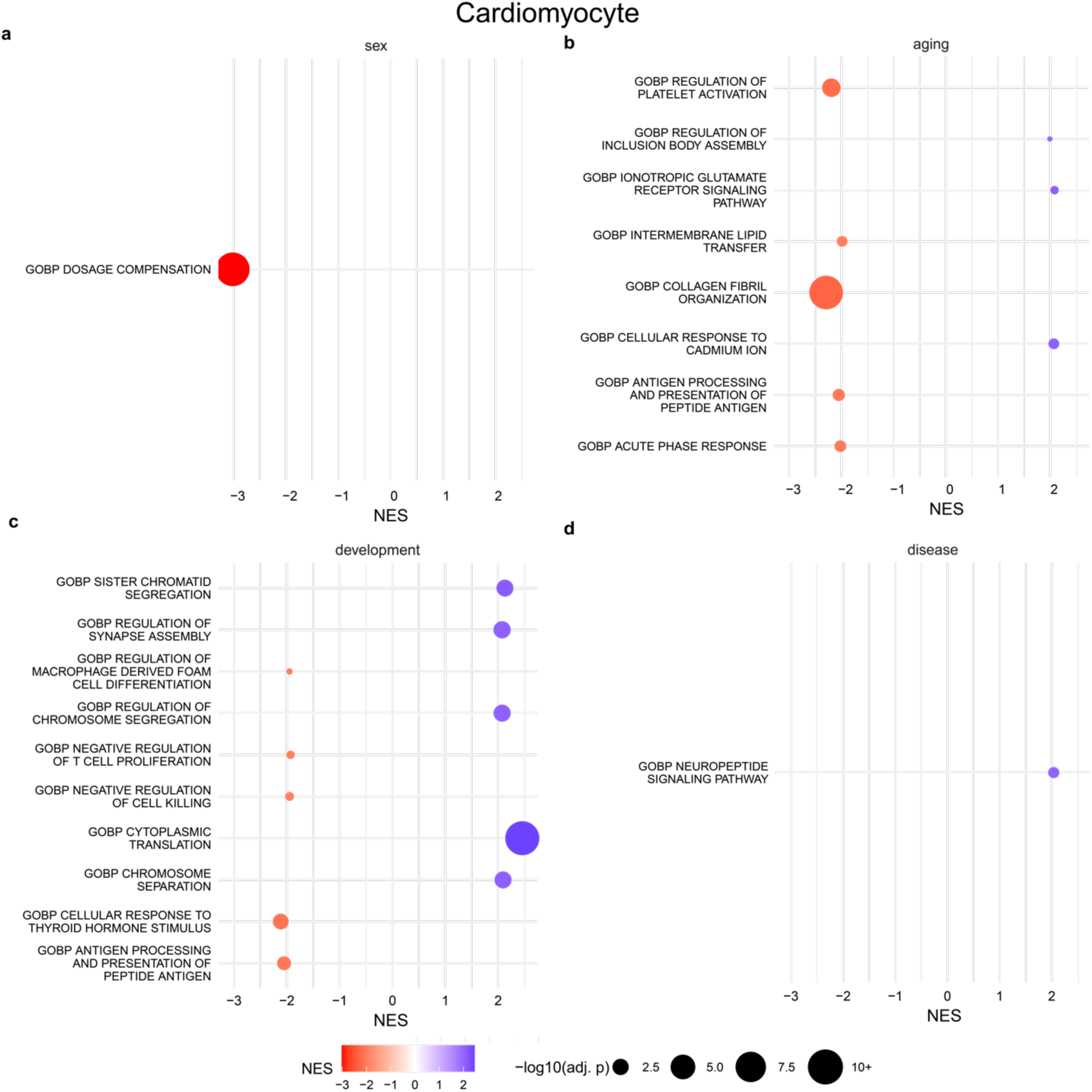
snRNA-seq Cardiomyocyte gene set enrichment analysis: Adipocyte gene set enrichment analysis using the ordered list of genes for **(a)** sex, **(b)** aging, **(c)** development and **(d)** disease contrasts. NES = normalized enrichment score, so a negative NES for the sex contrast (male vs. female) would be interpreted as gene sets enriched in females and a positive NES would be interpreted as gene sets enriched in males. Only the top 5 positive and negative NES gene sets, ranked by Benjamini-Hochberg adjusted p-value are displayed.

**Supplemental Figure 23.**
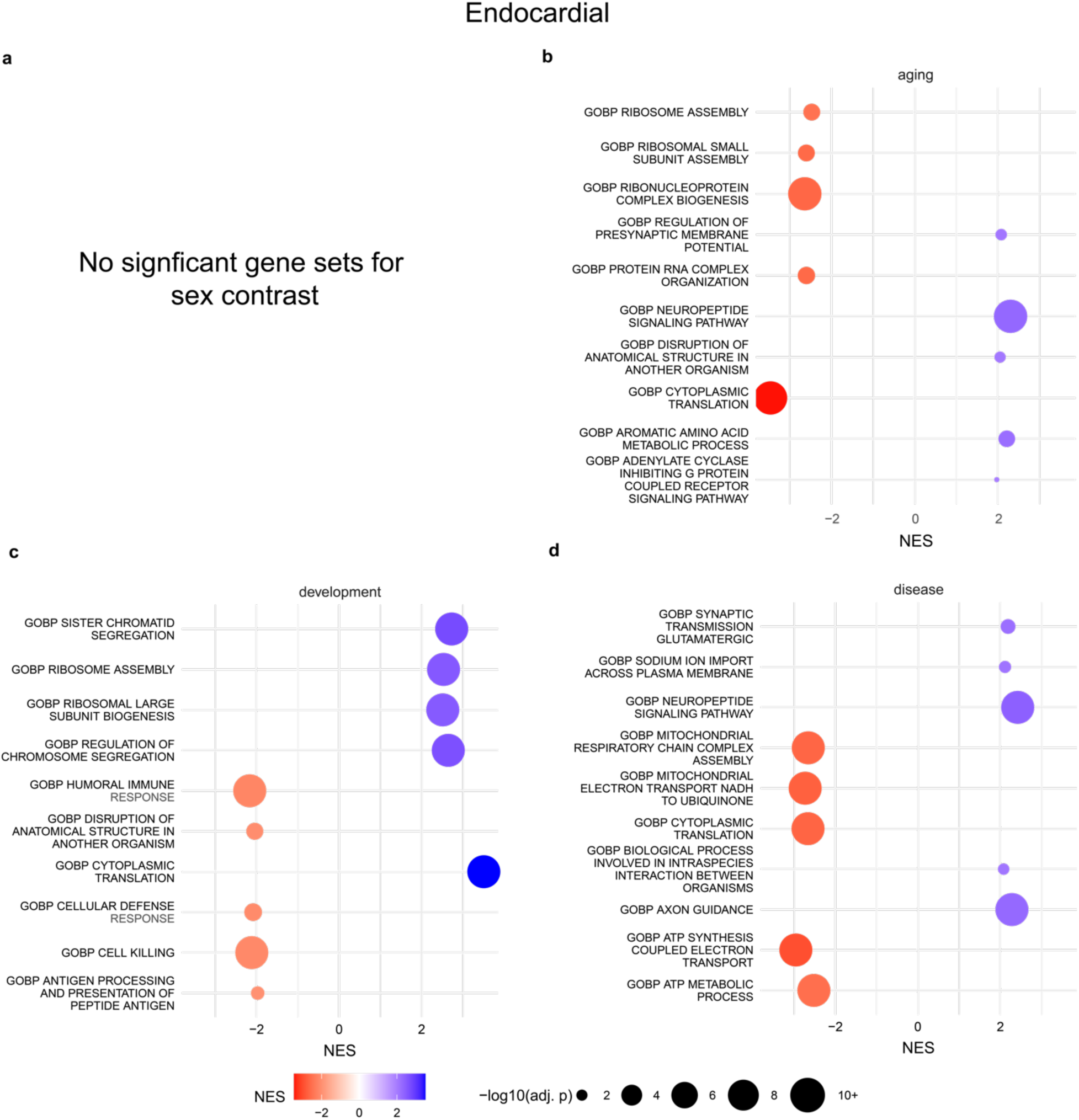
snRNA-seq Endocardial gene set enrichment analysis: Adipocyte gene set enrichment analysis using the ordered list of genes for **(a)** sex, **(b)** aging, **(c)** development and **(d)** disease contrasts. NES = normalized enrichment score, so a negative NES for the sex contrast (male vs. female) would be interpreted as gene sets enriched in females and a positive NES would be interpreted as gene sets enriched in males. Only the top 5 positive and negative NES gene sets, ranked by Benjamini-Hochberg adjusted p-value are displayed.

**Supplemental Figure 24.**
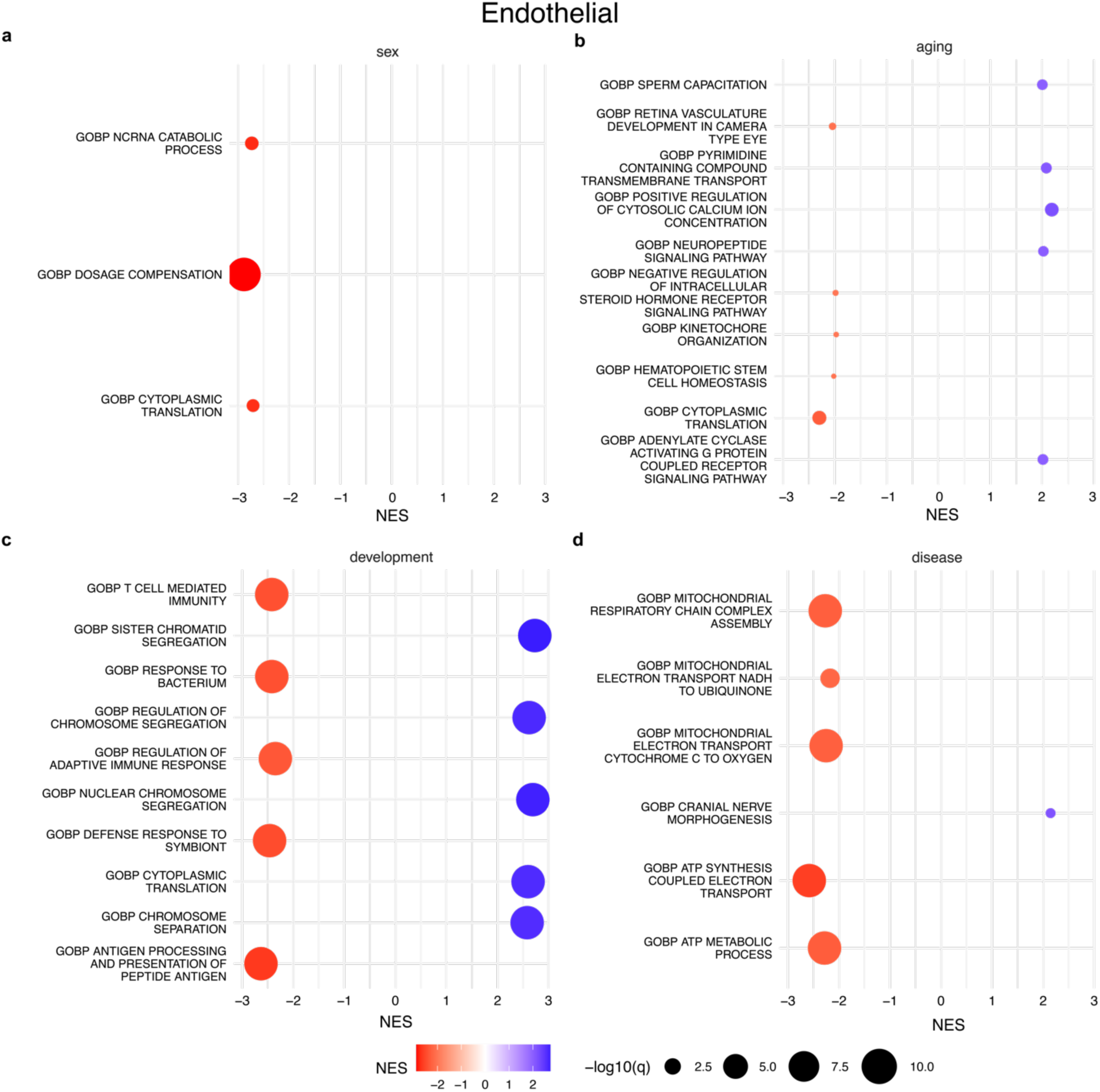
snRNA-seq Endothelial gene set enrichment analysis: Adipocyte gene set enrichment analysis using the ordered list of genes for **(a)** sex, **(b)** aging, **(c)** development and **(d)** disease contrasts. NES = normalized enrichment score, so a negative NES for the sex contrast (male vs. female) would be interpreted as gene sets enriched in females and a positive NES would be interpreted as gene sets enriched in males. Only the top 5 positive and negative NES gene sets, ranked by Benjamini-Hochberg adjusted p-value are displayed.

**Supplemental Figure 25.**
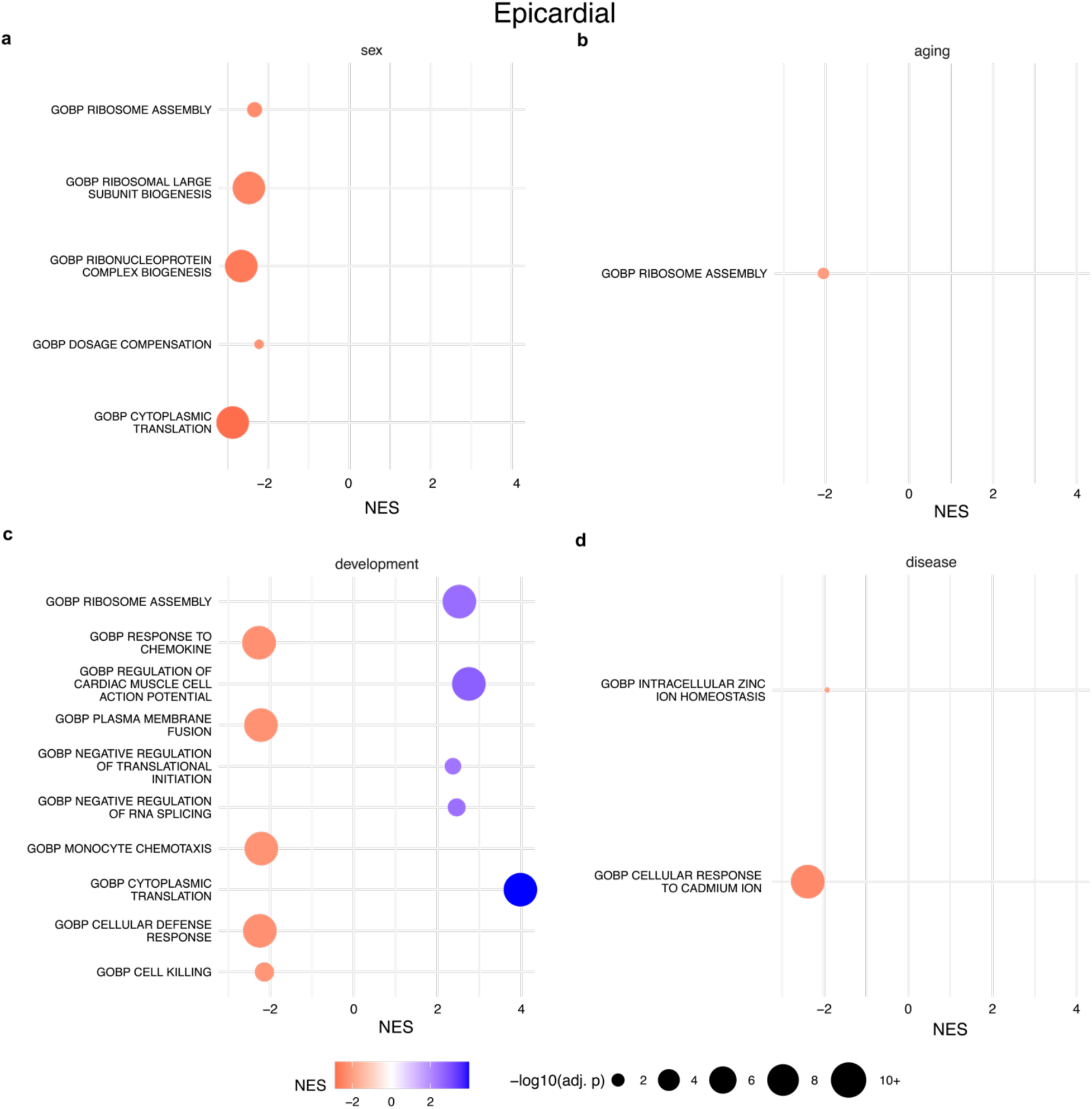
snRNA-seq Epicardial gene set enrichment analysis: Adipocyte gene set enrichment analysis using the ordered list of genes for **(a)** sex, **(b)** aging, **(c)** development and **(d)** disease contrasts. NES = normalized enrichment score, so a negative NES for the sex contrast (male vs. female) would be interpreted as gene sets enriched in females and a positive NES would be interpreted as gene sets enriched in males. Only the top 5 positive and negative NES gene sets, ranked by Benjamini-Hochberg adjusted p-value are displayed.

**Supplemental Figure 26.**
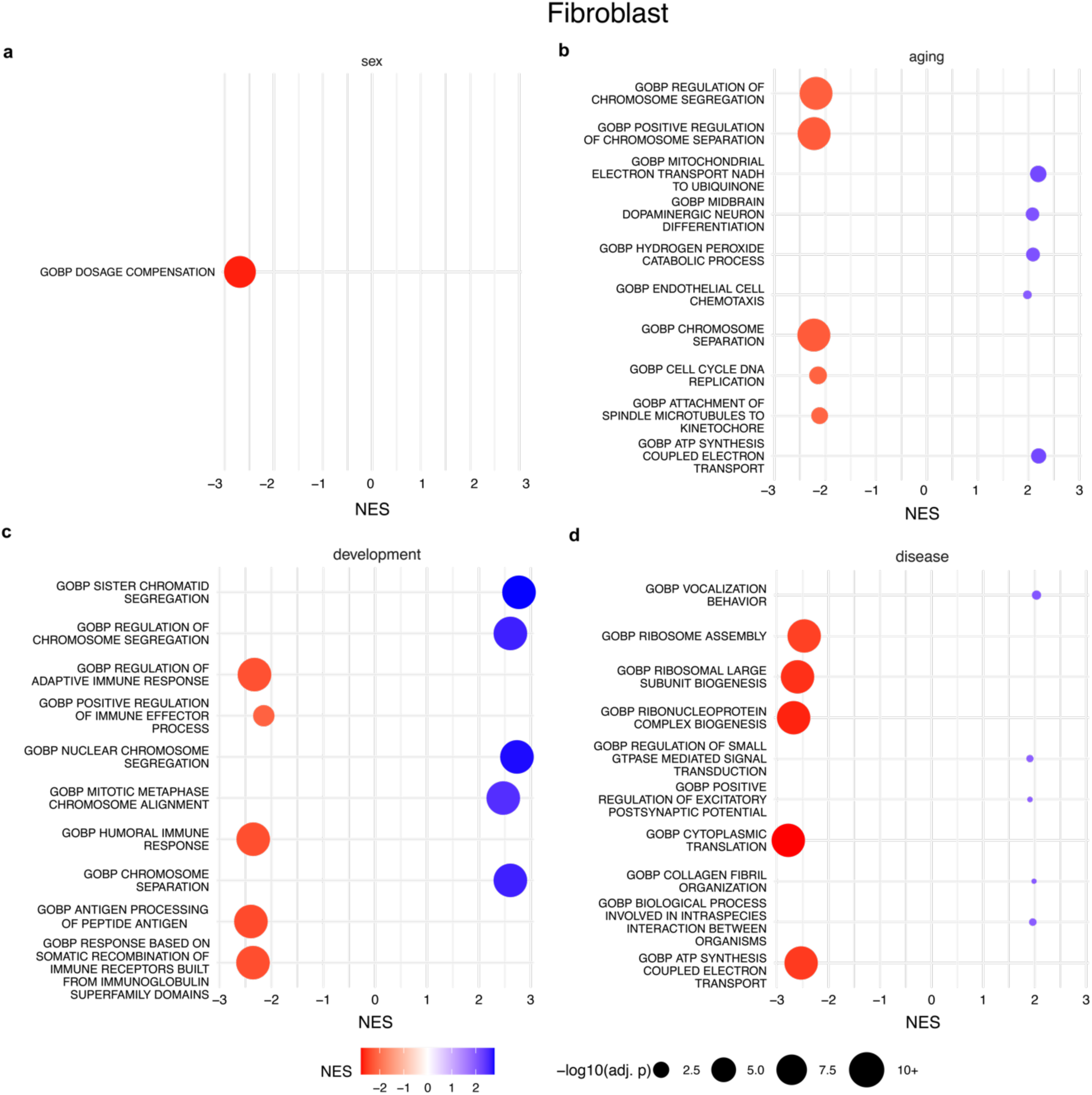
snRNA-seq Fibroblast gene set enrichment analysis: Adipocyte gene set enrichment analysis using the ordered list of genes for **(a)** sex, **(b)** aging, **(c)** development and **(d)** disease contrasts. NES = normalized enrichment score, so a negative NES for the sex contrast (male vs. female) would be interpreted as gene sets enriched in females and a positive NES would be interpreted as gene sets enriched in males. Only the top 5 positive and negative NES gene sets, ranked by Benjamini-Hochberg adjusted p-value are displayed.

**Supplemental Figure 27.**
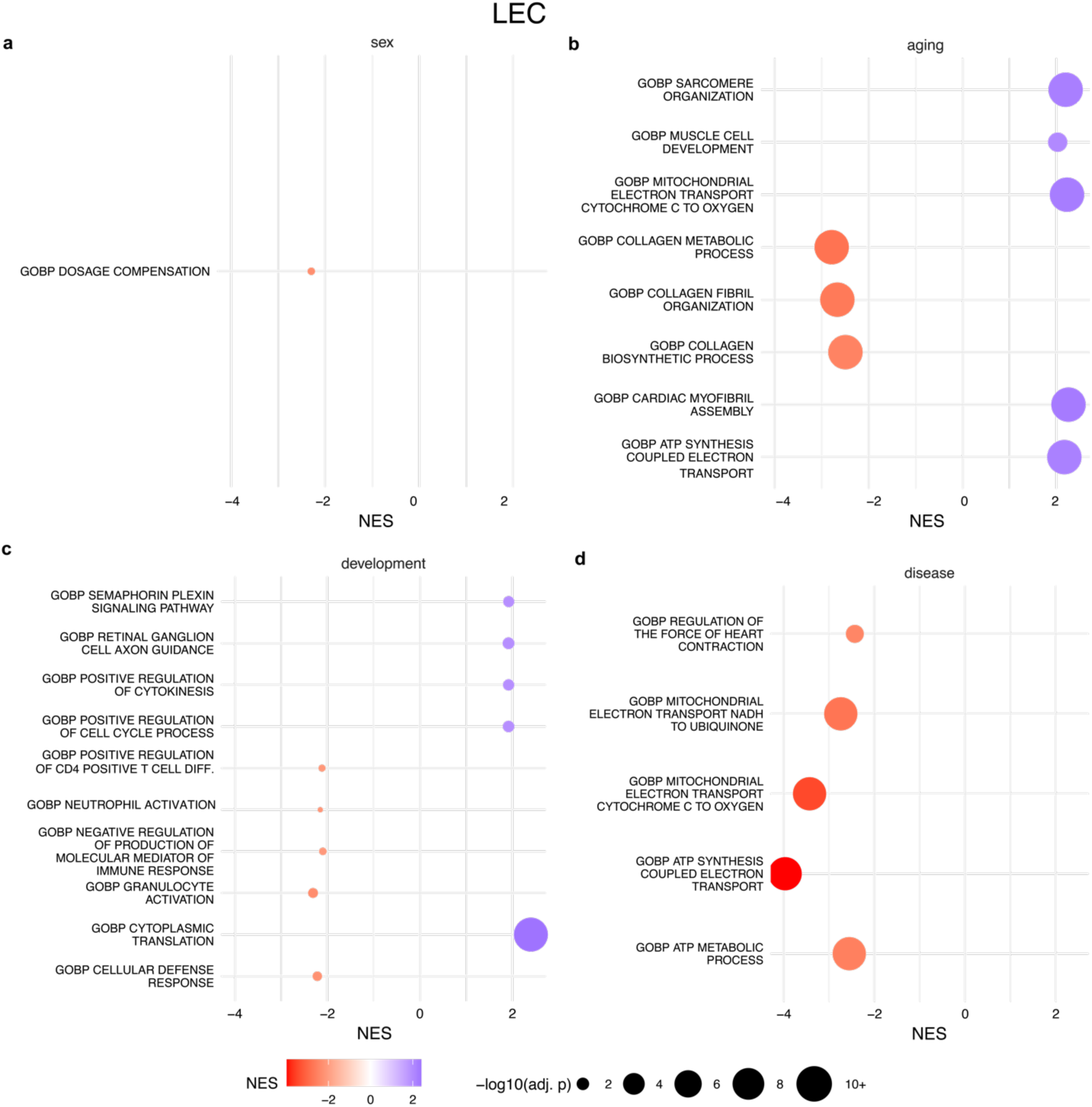
snRNA-seq LEC gene set enrichment analysis: Adipocyte gene set enrichment analysis using the ordered list of genes for **(a)** sex, **(b)** aging, **(c)** development and **(d)** disease contrasts. NES = normalized enrichment score, so a negative NES for the sex contrast (male vs. female) would be interpreted as gene sets enriched in females and a positive NES would be interpreted as gene sets enriched in males. Only the top 5 positive and negative NES gene sets, ranked by Benjamini-Hochberg adjusted p-value are displayed.

**Supplemental Figure 28.**
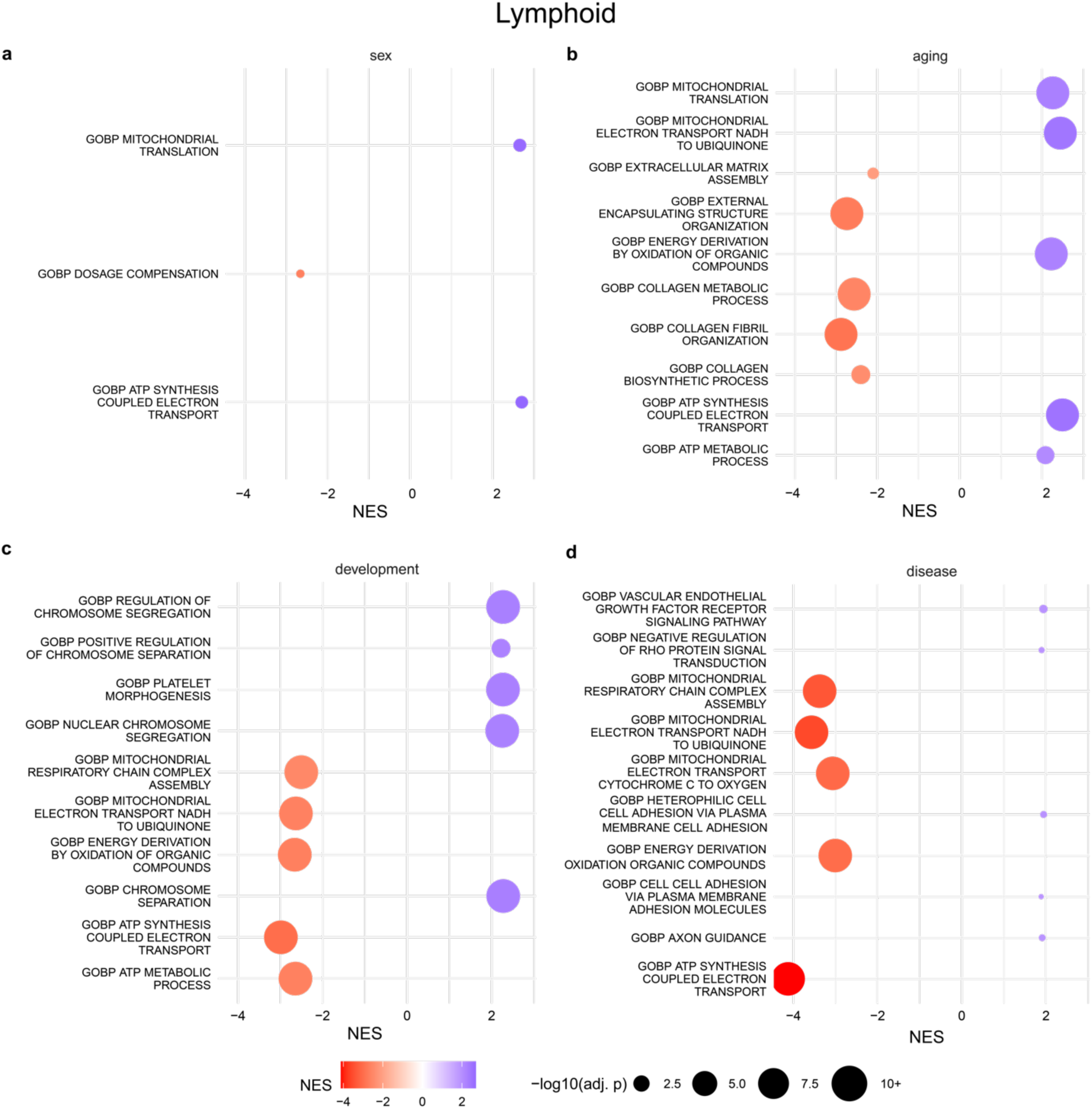
snRNA-seq Lymphoid gene set enrichment analysis: Adipocyte gene set enrichment analysis using the ordered list of genes for **(a)** sex, **(b)** aging, **(c)** development and **(d)** disease contrasts. NES = normalized enrichment score, so a negative NES for the sex contrast (male vs. female) would be interpreted as gene sets enriched in females and a positive NES would be interpreted as gene sets enriched in males. Only the top 5 positive and negative NES gene sets, ranked by Benjamini-Hochberg adjusted p-value are displayed.

**Supplemental Figure 29.**
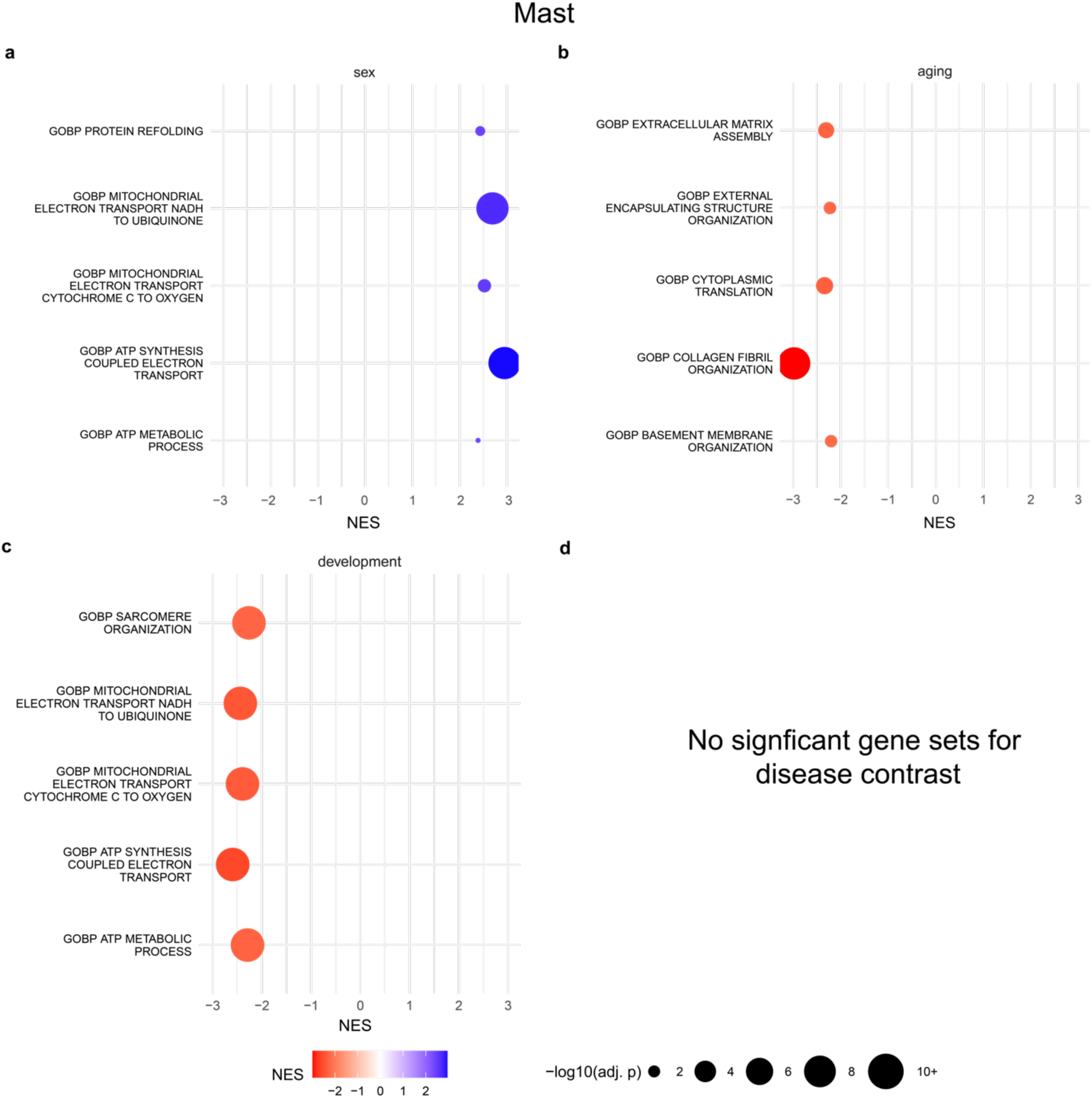
snRNA-seq Mast gene set enrichment analysis: Adipocyte gene set enrichment analysis using the ordered list of genes for **(a)** sex, **(b)** aging, **(c)** development and **(d)** disease contrasts. NES = normalized enrichment score, so a negative NES for the sex contrast (male vs. female) would be interpreted as gene sets enriched in females and a positive NES would be interpreted as gene sets enriched in males. Only the top 5 positive and negative NES gene sets, ranked by Benjamini-Hochberg adjusted p-value are displayed.

**Supplemental Figure 30.**
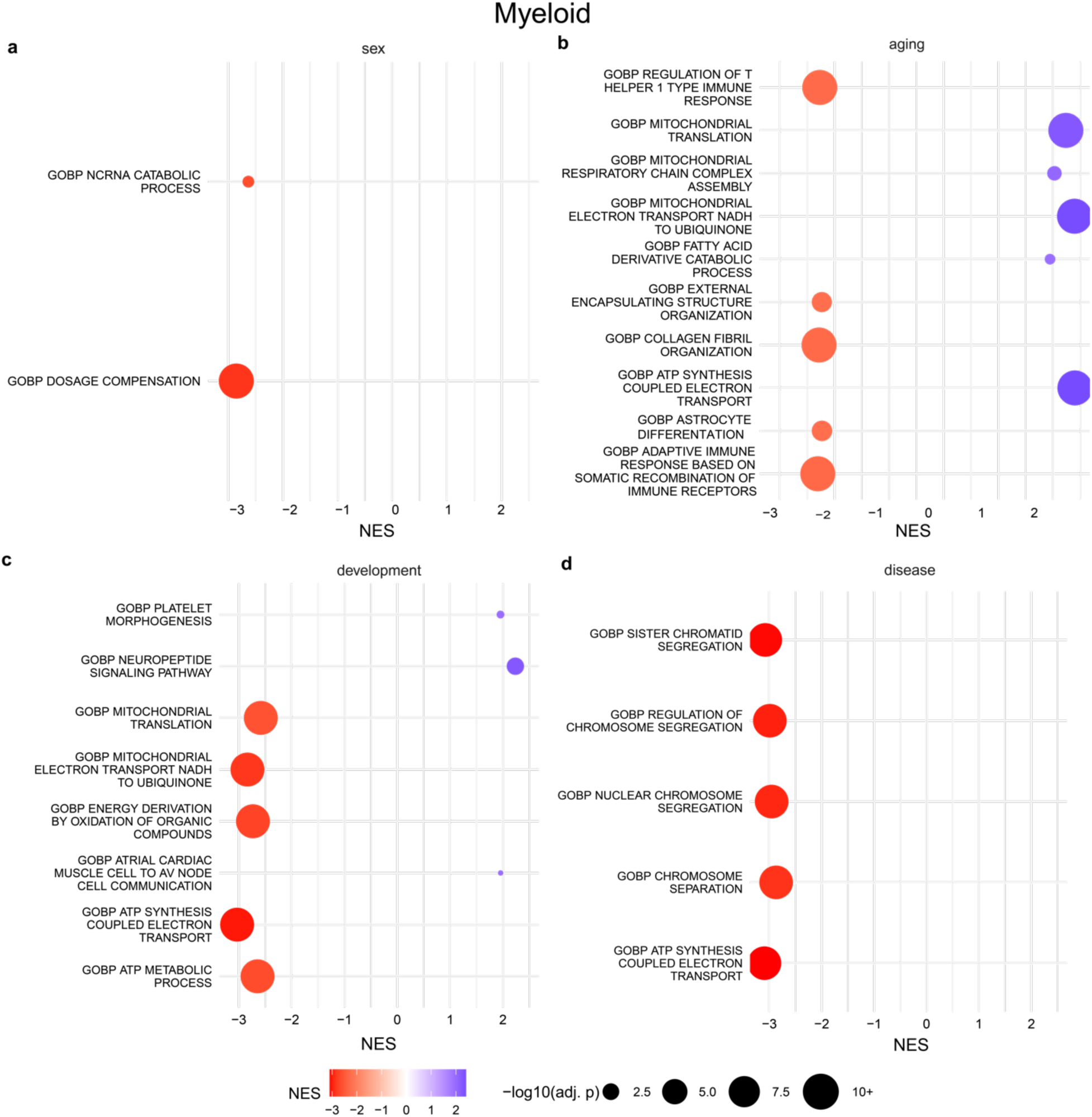
snRNA-seq Myeloid gene set enrichment analysis: Adipocyte gene set enrichment analysis using the ordered list of genes for **(a)** sex, **(b)** aging, **(c)** development and **(d)** disease contrasts. NES = normalized enrichment score, so a negative NES for the sex contrast (male vs. female) would be interpreted as gene sets enriched in females and a positive NES would be interpreted as gene sets enriched in males. Only the top 5 positive and negative NES gene sets, ranked by Benjamini-Hochberg adjusted p-value are displayed.

**Supplemental Figure 31.**
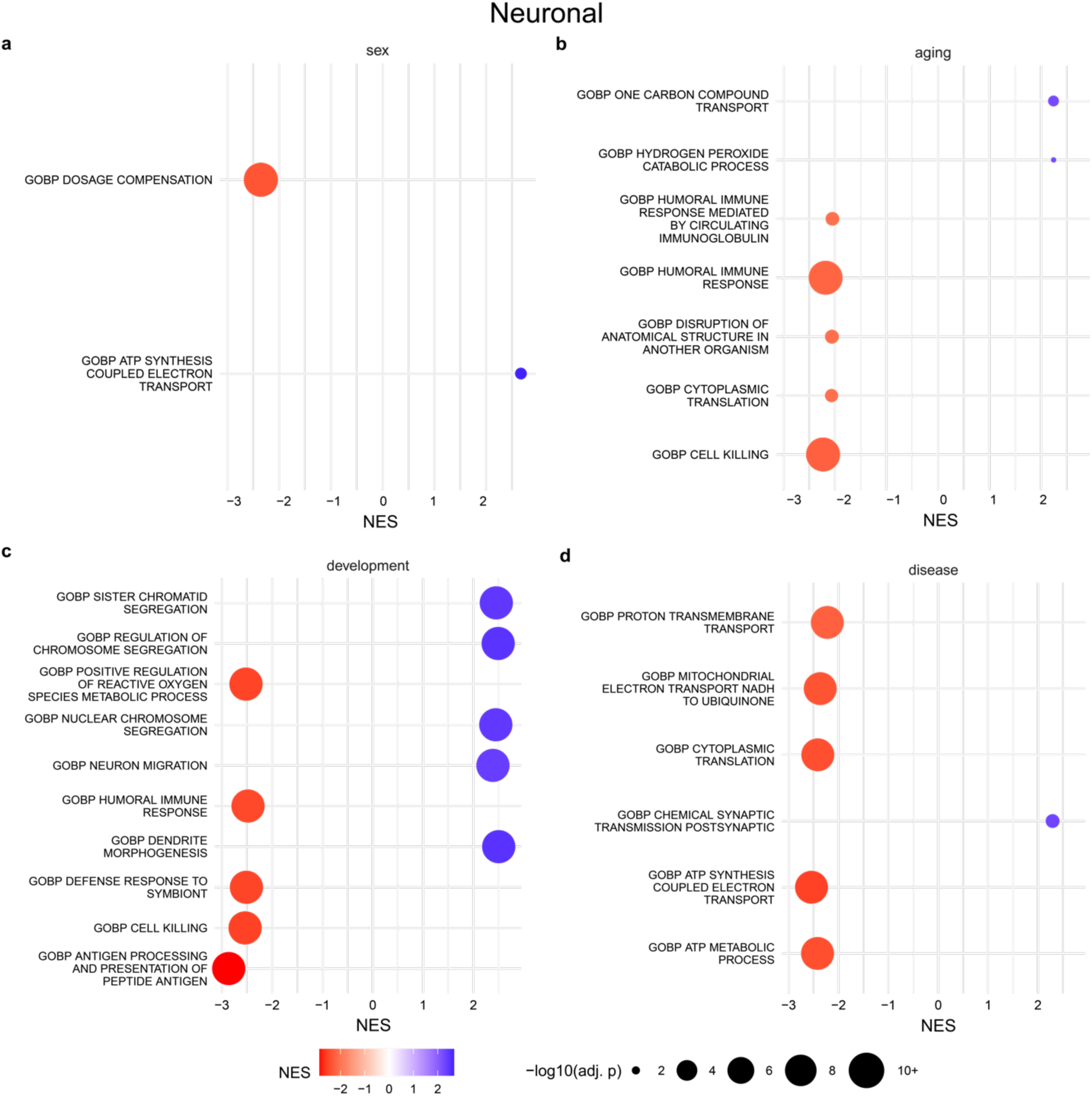
snRNA-seq Neuronal gene set enrichment analysis: Adipocyte gene set enrichment analysis using the ordered list of genes for **(a)** sex, **(b)** aging, **(c)** development and **(d)** disease contrasts. NES = normalized enrichment score, so a negative NES for the sex contrast (male vs. female) would be interpreted as gene sets enriched in females and a positive NES would be interpreted as gene sets enriched in males. Only the top 5 positive and negative NES gene sets, ranked by Benjamini-Hochberg adjusted p-value are displayed.

**Supplemental Figure 32.**
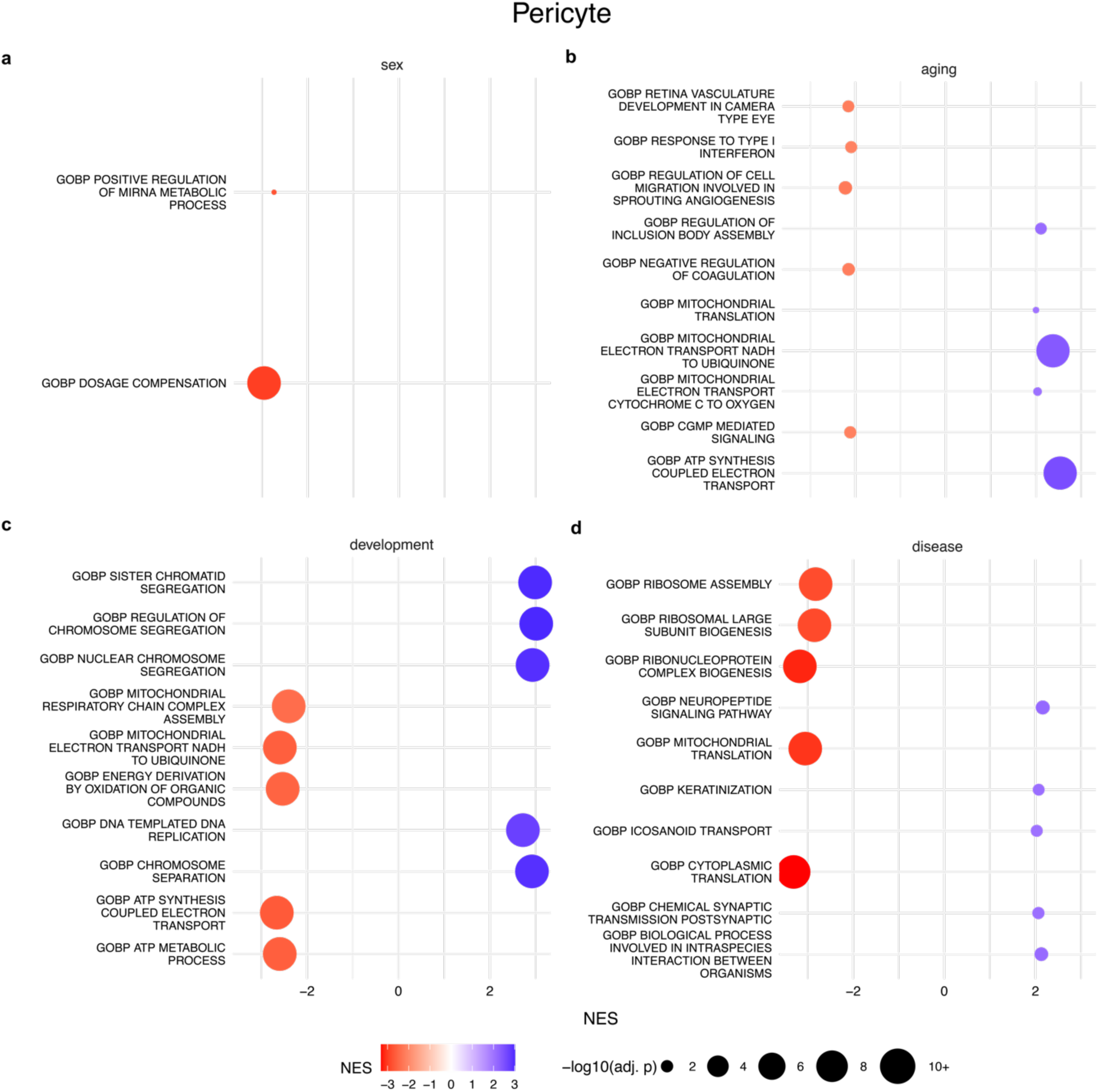
snRNA-seq Pericyte gene set enrichment analysis: Adipocyte gene set enrichment analysis using the ordered list of genes for **(a)** sex, **(b)** aging, **(c)** development and **(d)** disease contrasts. NES = normalized enrichment score, so a negative NES for the sex contrast (male vs. female) would be interpreted as gene sets enriched in females and a positive NES would be interpreted as gene sets enriched in males. Only the top 5 positive and negative NES gene sets, ranked by Benjamini-Hochberg adjusted p-value are displayed.

**Supplemental Figure 33.**
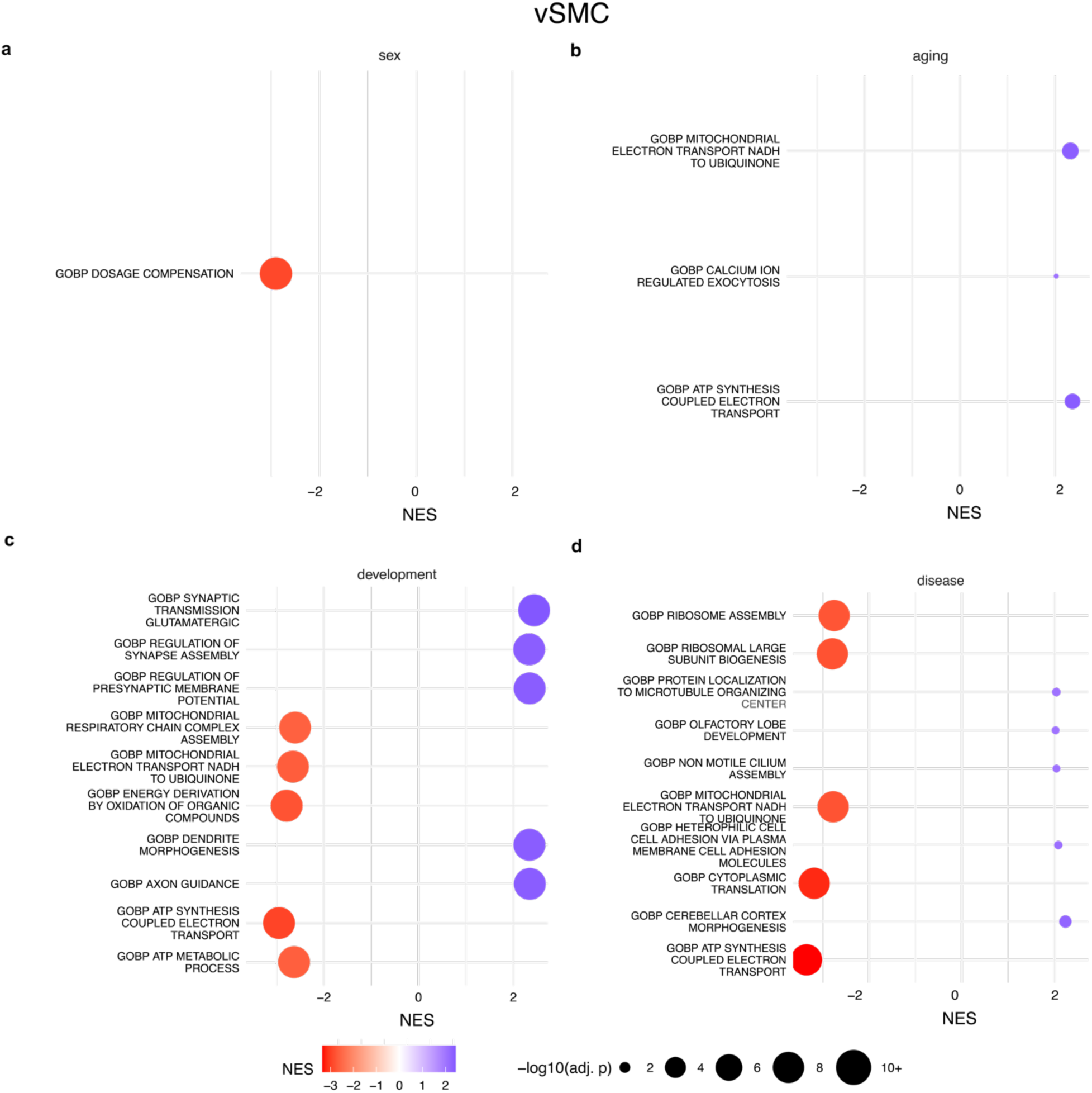
snRNA-seq vSMC gene set enrichment analysis: Adipocyte gene set enrichment analysis using the ordered list of genes for **(a)** sex, **(b)** aging, **(c)** development and **(d)** disease contrasts. NES = normalized enrichment score, so a negative NES for the sex contrast (male vs. female) would be interpreted as gene sets enriched in females and a positive NES would be interpreted as gene sets enriched in males. Only the top 5 positive and negative NES gene sets, ranked by Benjamini-Hochberg adjusted p-value are displayed.

**Supplemental Figure 34.**
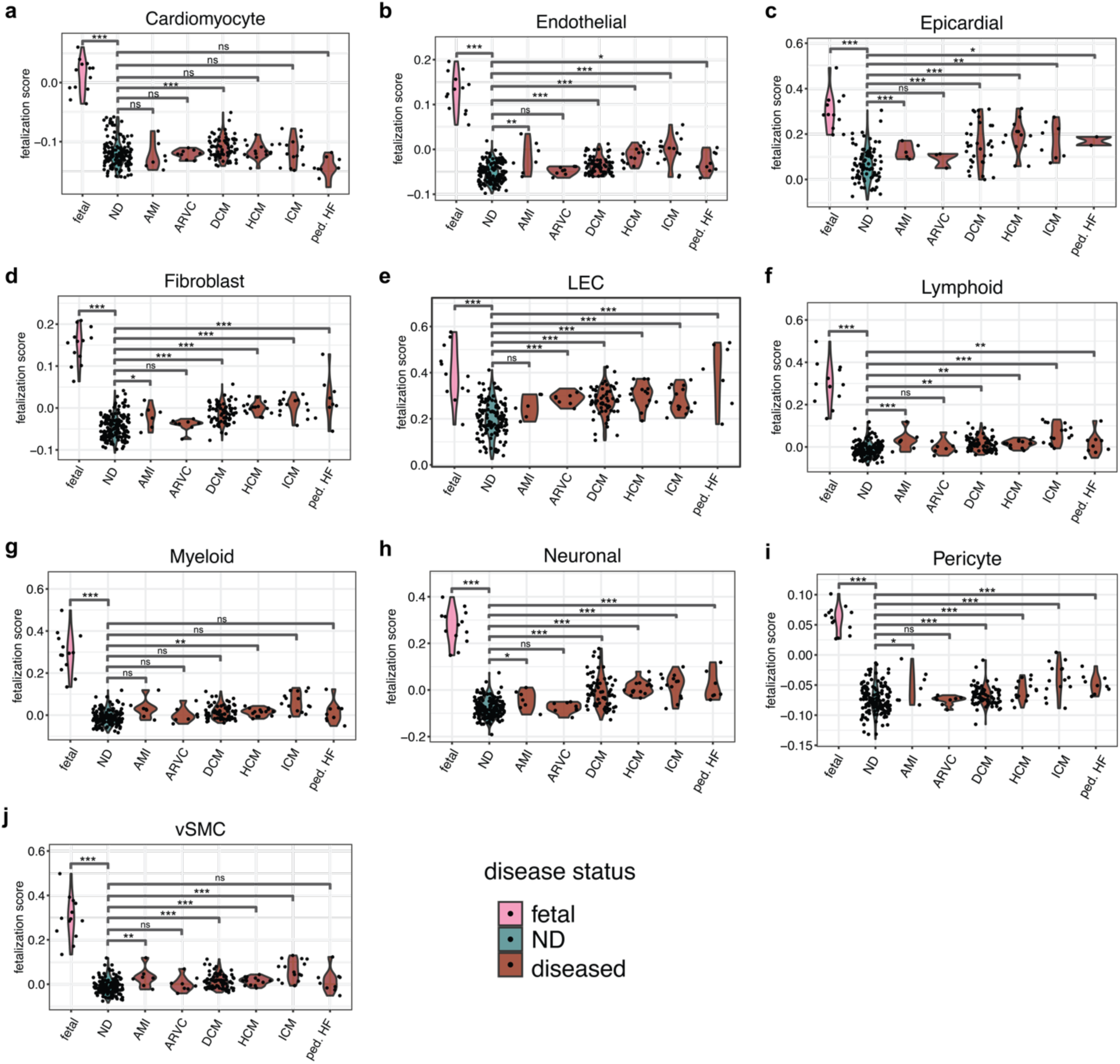
Fetal reactivation score across disease subtypes: Using the intersecting up DEGs in fetal (vs. postnatal young) and disease (binarized: Y vs N) contrasts, we calculate a fetal reactivation score per cell using scanpy’s score_genes() function. The mean score per donor was calculated and displayed in the violin plots above. We used cell-type specific fetal reactivation genes sets for all cell types (Cardiomyocyte, Epicardial, Endothelial, Fibroblast, LEC, Lymphoid, Myeloid, Neuronal, Pericyte, vSMC) that displayed significant overlap between fetal and disease DEGs.

**Supplemental Figure 35.**
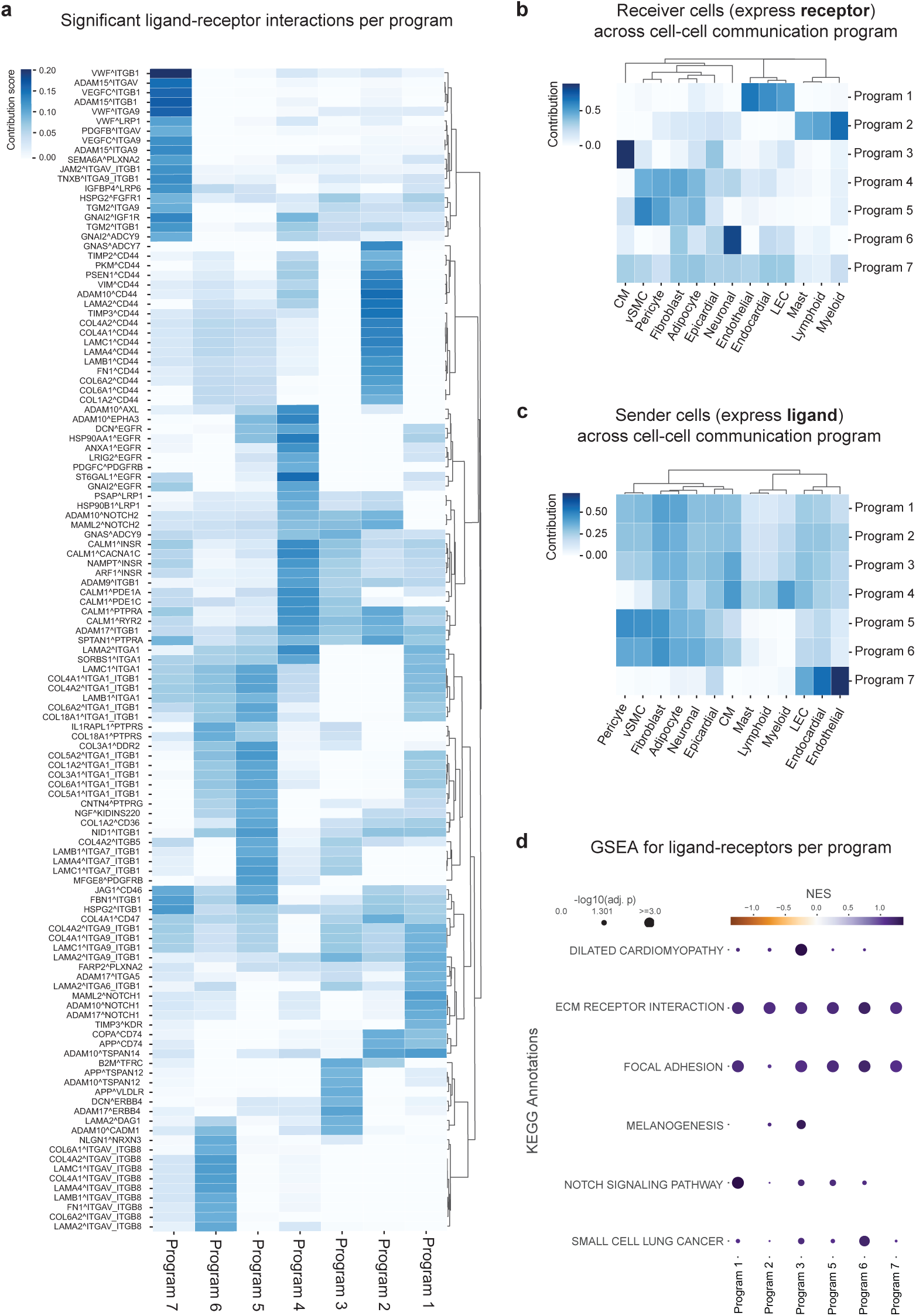
Additional cell-cell communication analyses with liana x tensorcell2cell: **(a)** Significant ligand-receptor interactions per cell-cell communication program **(b)** Contribution of receptor-expressing receiver cell types in each program **(c)** Contribution of sender-expression receiver cell types in each program **(d)** GSEA for ligand-receptor interactions in each program.

**Supplemental Figure 36.**
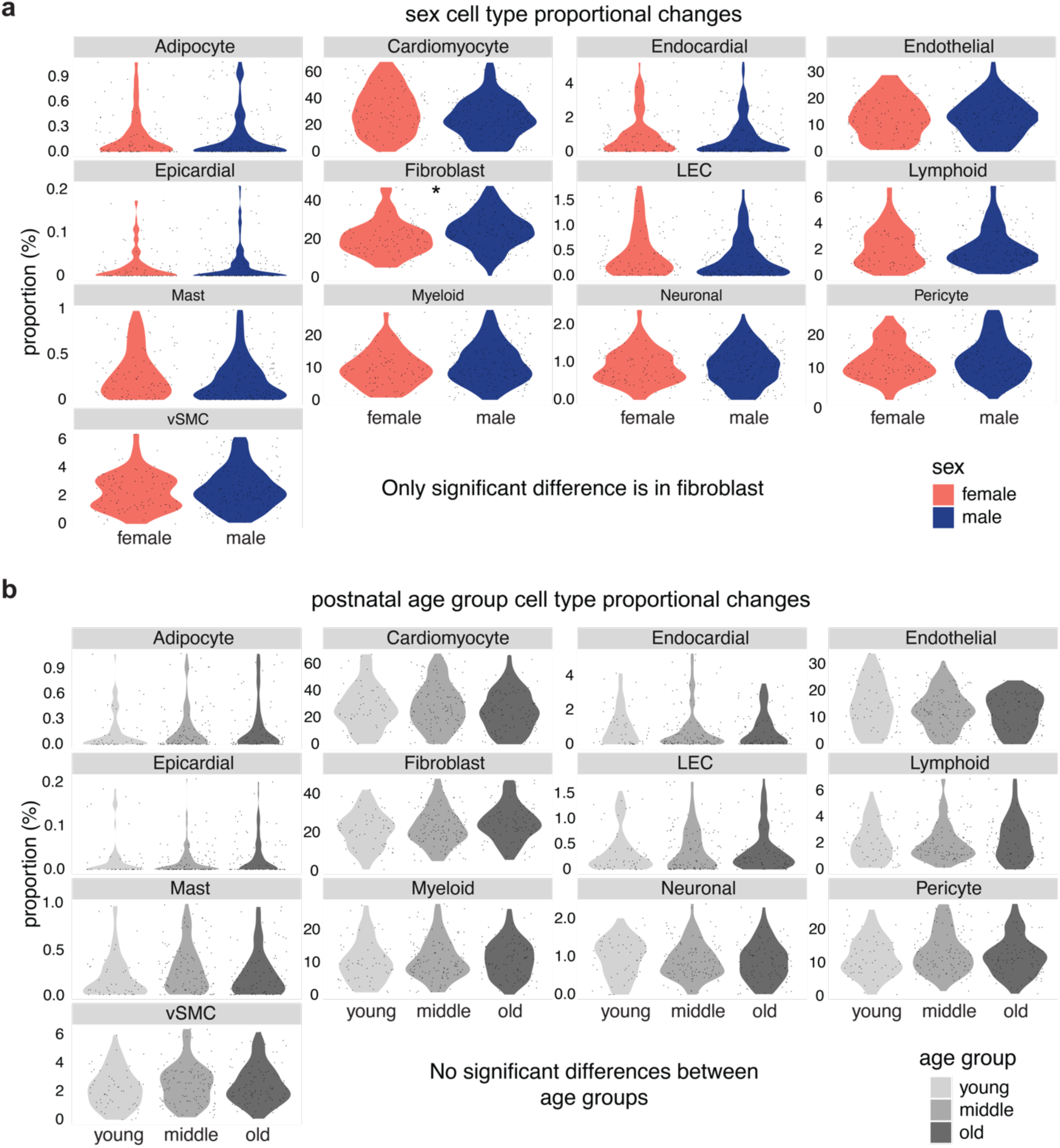
Cell type proportion analysis by sex and age group: Cell type proportional analysis by sex and aging. Significant changes per contrast was computed using scanpro. **(a)** Cell type proportions by sex. Statistical significance is determined while for technology, study, and age. There are no significant differences except with higher proportions of fibroblasts in males. **(b)** Cell type proportions by age group (young, middle, and old). Statistical significance is determined while for sex, study, and age. There are no cell types displaying significant proportional changes across age.

**Supplemental Figure 37.**
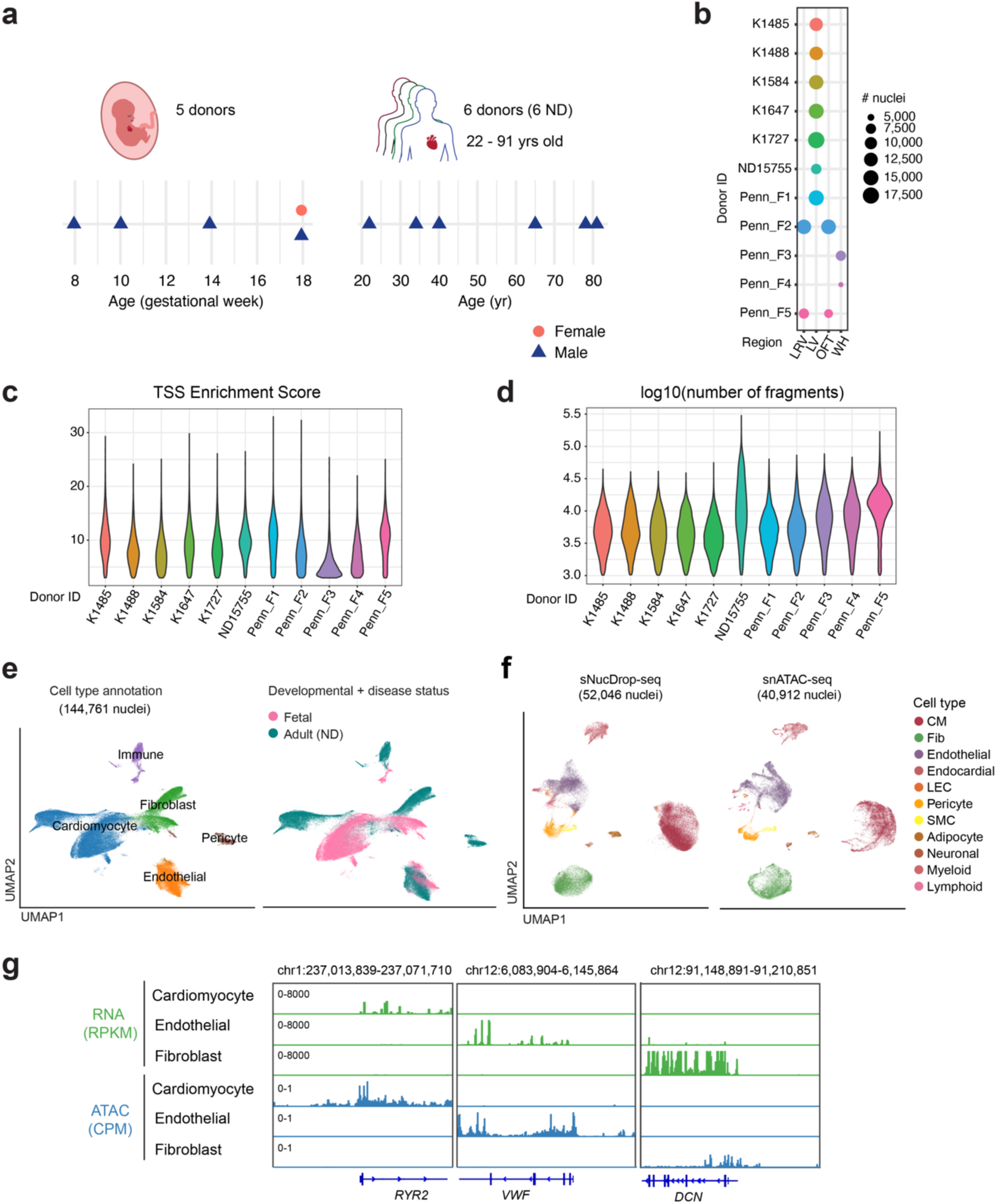
Single-nucleus ATAC-seq dataset of human cardiac development and aging generated in this study: **(a)** Summary of donors by sex and developmental stage. **(b)** Sampled regions and number of nuclei per donor **(c-d)** Summary statistics of snATAC-seq libraries, including TSS enrichment score and # of fragment per nucleus, respectively. (**e**) UMAP by cell type and developmental and disease status, respectively. (**f**) UMAP of integrated snRNA-seq (left) and snATAC-seq (right) datasets of adult human left ventricle (LV, non-diseased) from this study. The cell-types are annotated based on snRNA-seq. **(g)** Genomic track views of normalized gene expression (RNA) and chromatin accessibility (ATAC) for three cell-type specific marker genes (*RYR2* for cardiomyocytes, *VWF* for endothelial cells, and *DCN* for fibroblasts).

**Supplemental Figure 38.**
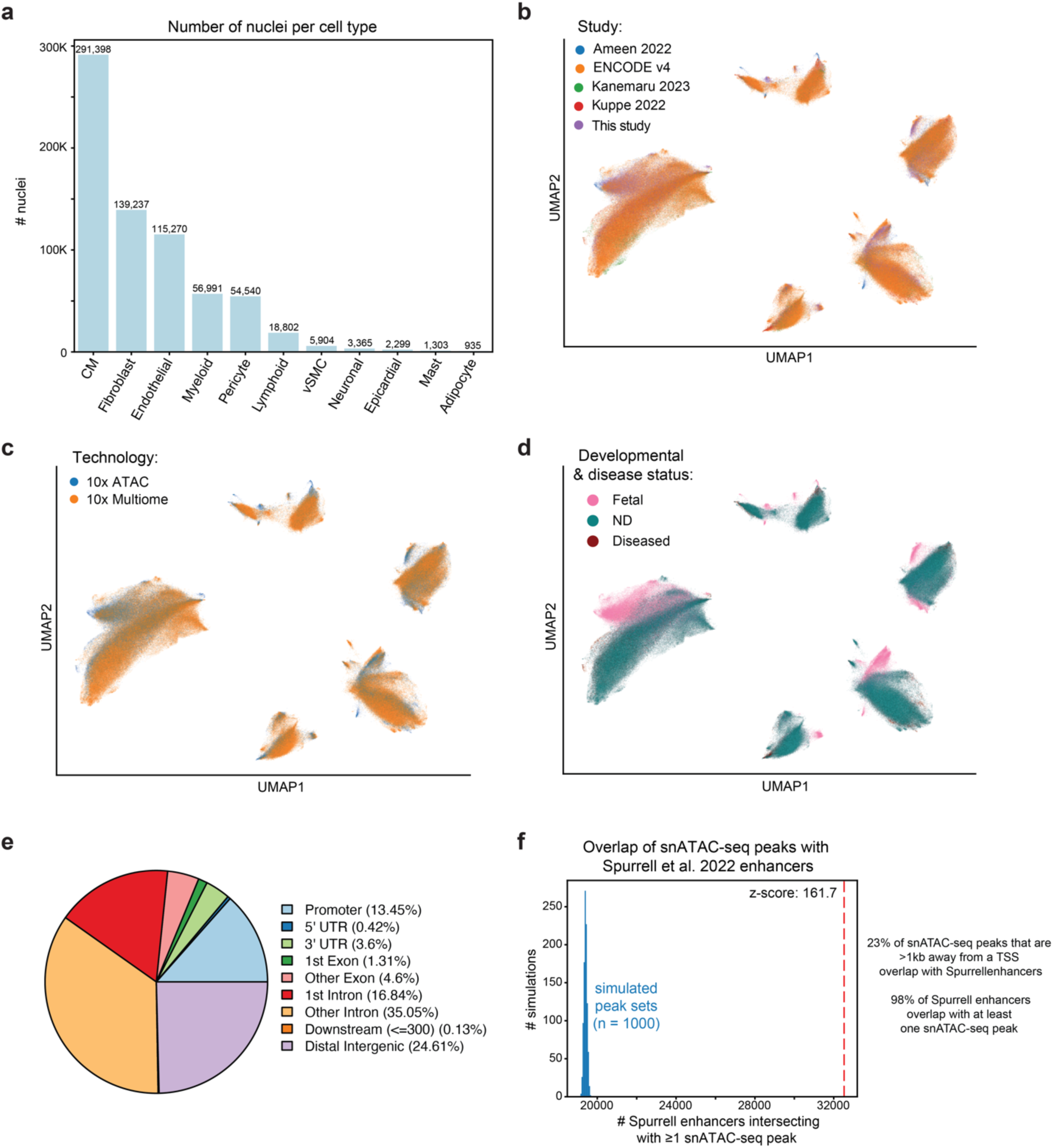
Additional snATAC-seq plots: **(a)** Number of nuclei per cell type in aggregated snATAC-seq dataset **(b-d)** UMAP for snATAC-seq plots by study, technology, and developmental + disease status, respectively **(e)** Genomic feature annotation of snATAC-seq peaks using ChIPSeeker (**f)** Overlap between non-promoter proximal peaks (>1 kb away from TSS) and Spurrell 2022 enhancers

**Supplemental Figure 39.**
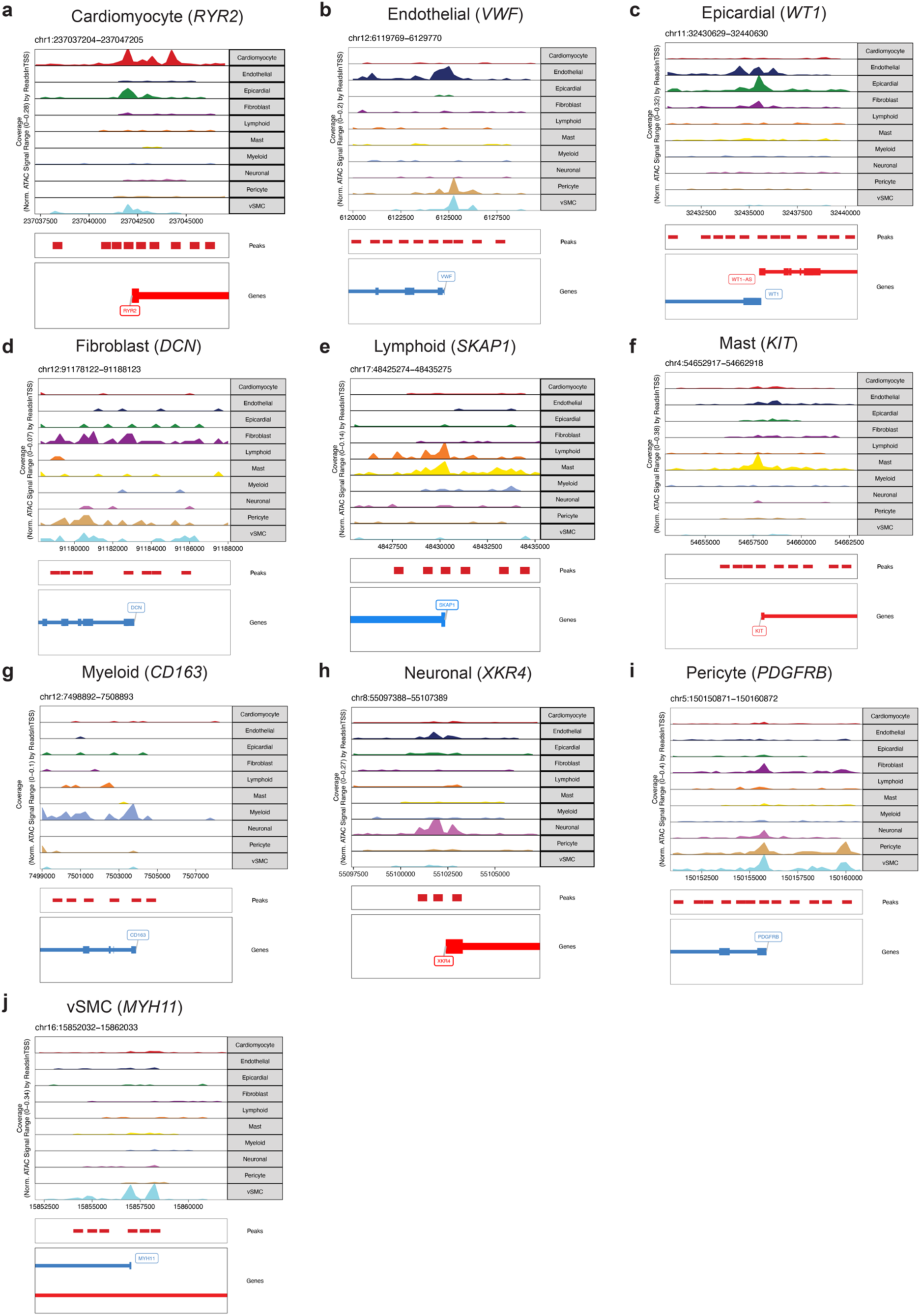
Marker gene promoter accessibility using snATAC-seq: Genome browser views of the snATAC-seq accessibility tracks for the promoter region (+/- 1 kb) of marker genes in Fig. 1e, generated using ArchR.

**Supplemental Figure 40.**
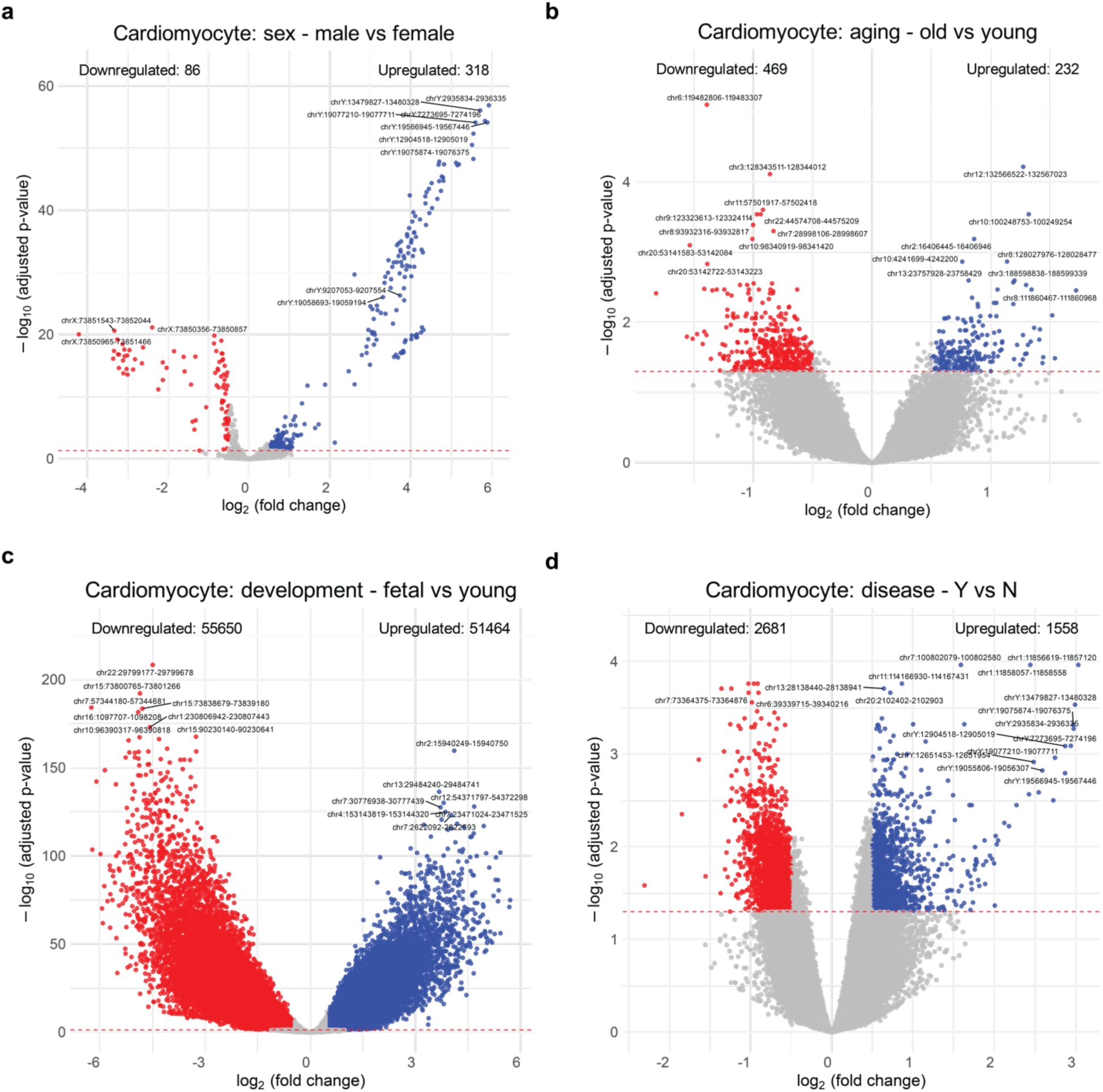
snATAC-seq Cardiomyocyte volcano plots: Using DESeq2, donor-pseudobulked snATAC-seq Cardiomyocyte volcano plots for the **(a)** sex, **(b)** aging, **(c)** development and **(d)** disease contrasts. Significance thresholds: |log2 fold change| > 0.5 and Benjamini-Hochberg adjusted p-value < 0.05.

**Supplemental Figure 41.**
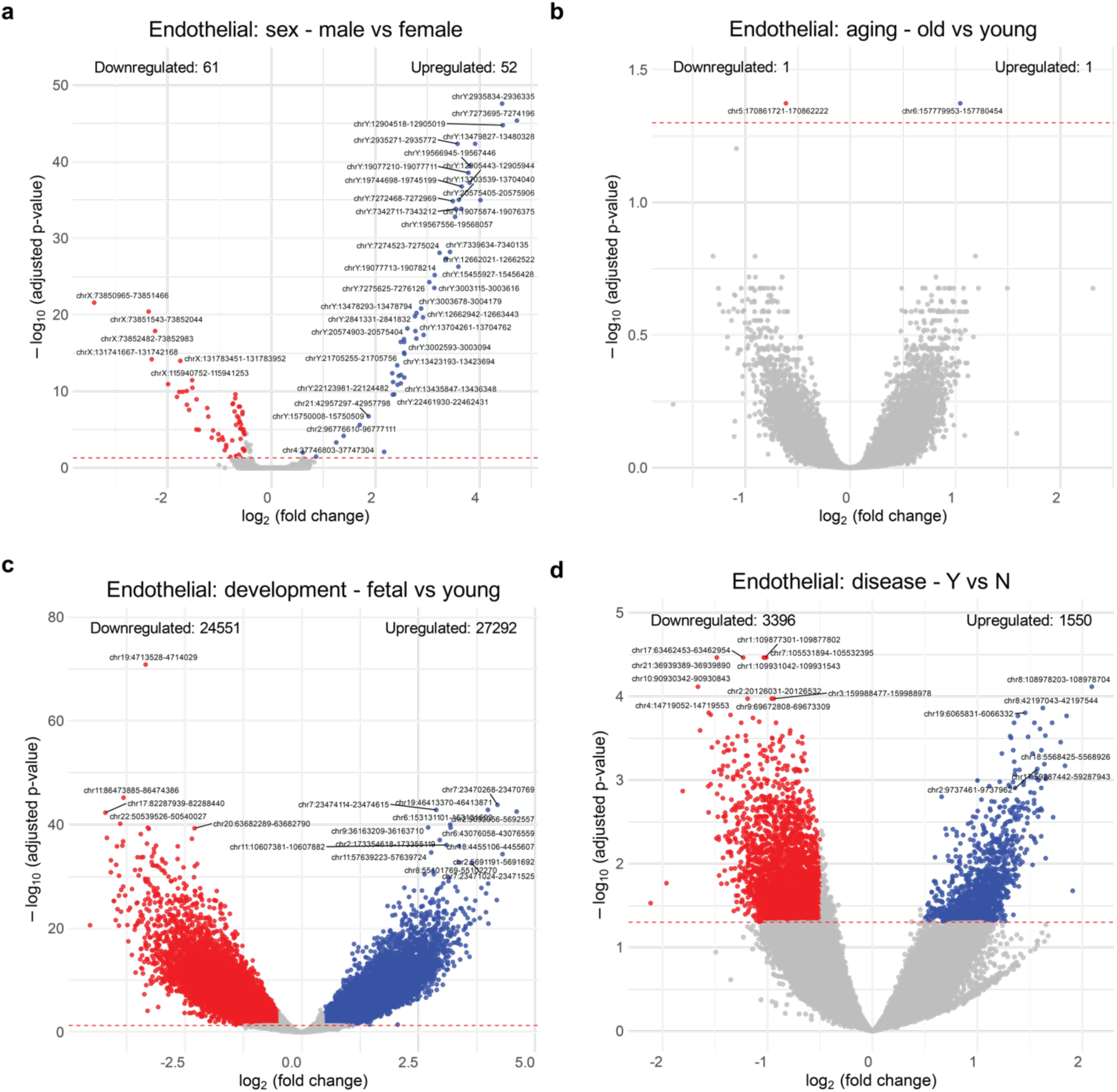
snATAC-seq Endothelial volcano plots: Using DESeq2, donor-pseudobulked snATAC-seq Endothelial volcano plots for the **(a)** sex, **(b)** aging, **(c)** development and **(d)** disease contrasts. Significance thresholds: |log2 fold change| > 0.5 and Benjamini-Hochberg adjusted p-value < 0.05.

**Supplemental Figure 42.**
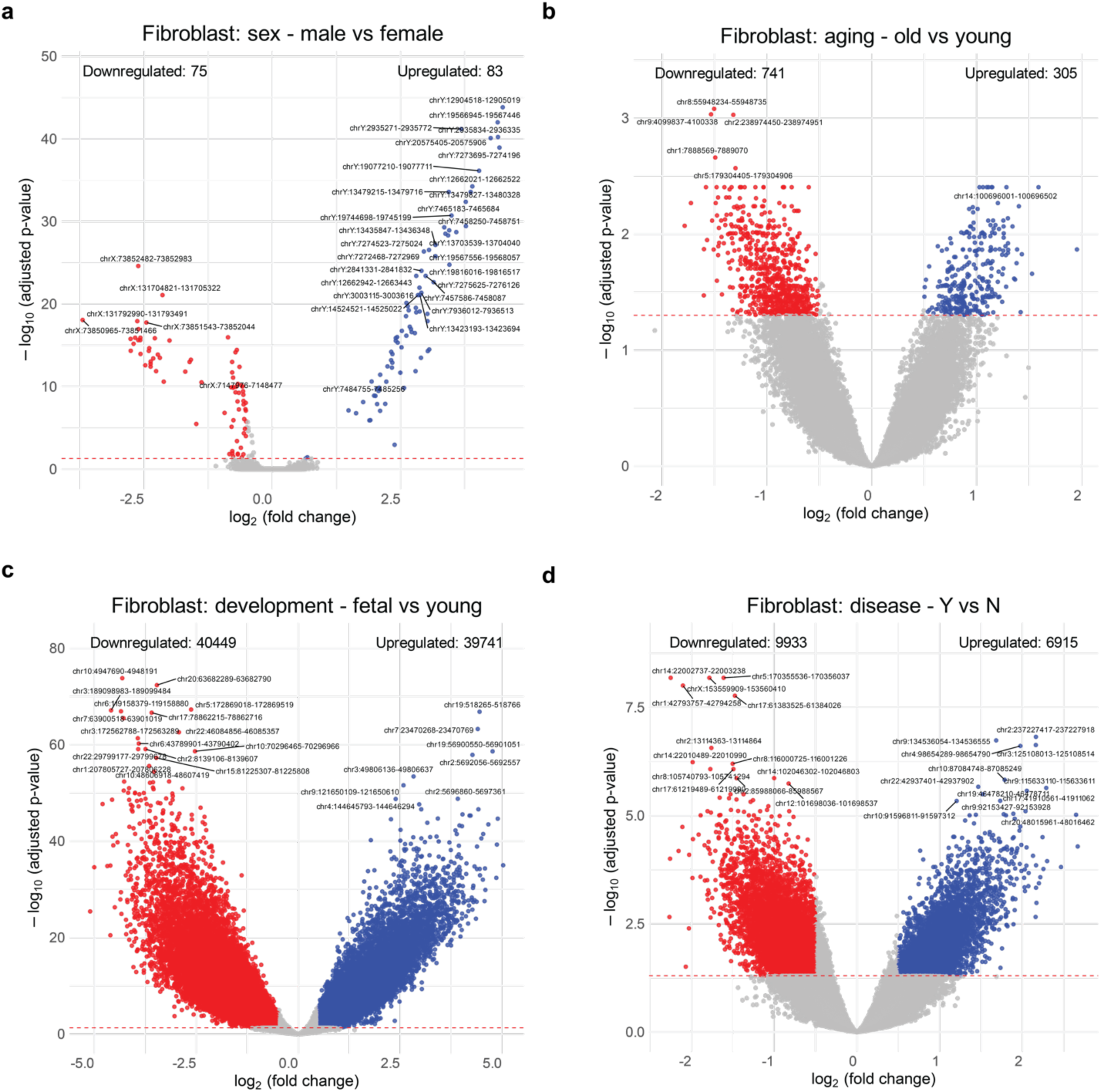
snATAC-seq Fibroblast volcano plots: Using DESeq2, donor-pseudobulked snATAC-seq Fibroblast volcano plots for the **(a)** sex, **(b)** aging, **(c)** development and **(d)** disease contrasts. Significance thresholds: |log2 fold change| > 0.5 and Benjamini-Hochberg adjusted p-value < 0.05.

**Supplemental Figure 43.**
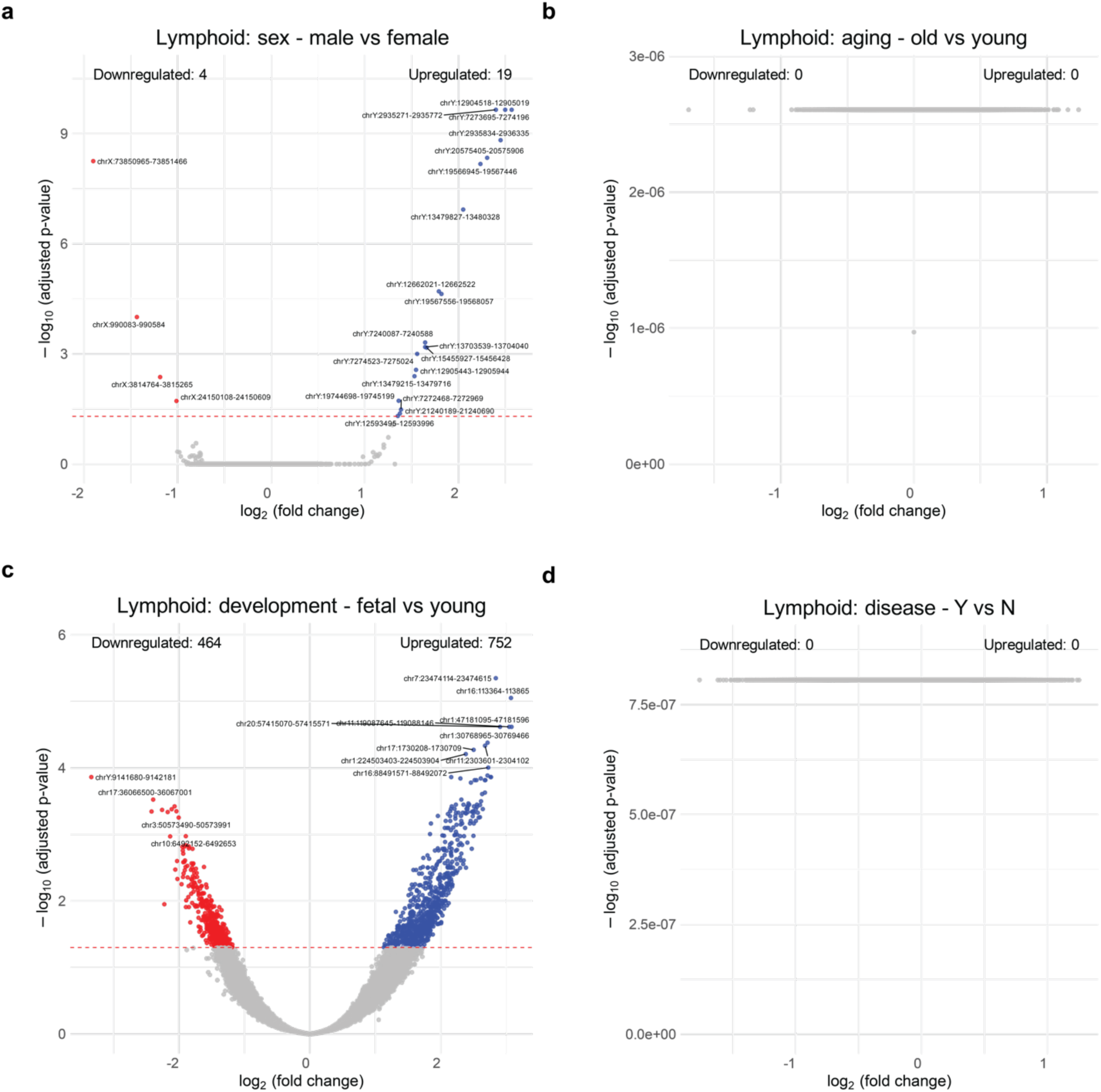
snATAC-seq Lymphoid volcano plots: Using DESeq2, donor-pseudobulked snATAC-seq Lymphoid volcano plots for the **(a)** sex, **(b)** aging, **(c)** development and **(d)** disease contrasts. Significance thresholds: |log2 fold change| > 0.5 and Benjamini-Hochberg adjusted p-value < 0.05.

**Supplemental Figure 44.**
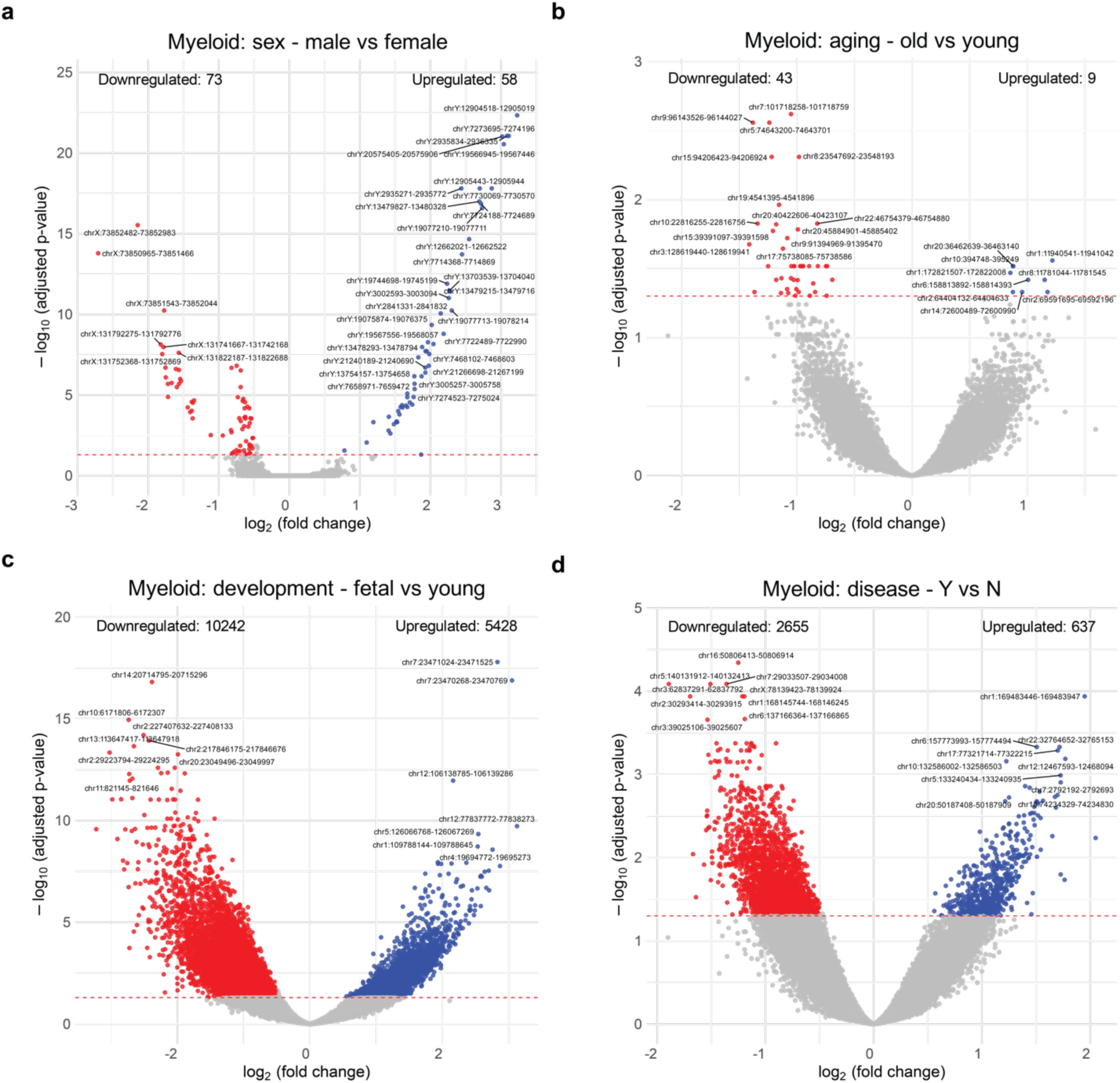
snATAC-seq Myeloid volcano plots: Using DESeq2, donor-pseudobulked snATAC-seq Myeloid volcano plots for the **(a)** sex, **(b)** aging, **(c)** development and **(d)** disease contrasts. Significance thresholds: |log2 fold change| > 0.5 and Benjamini-Hochberg adjusted p-value < 0.05.

**Supplemental Figure 45.**
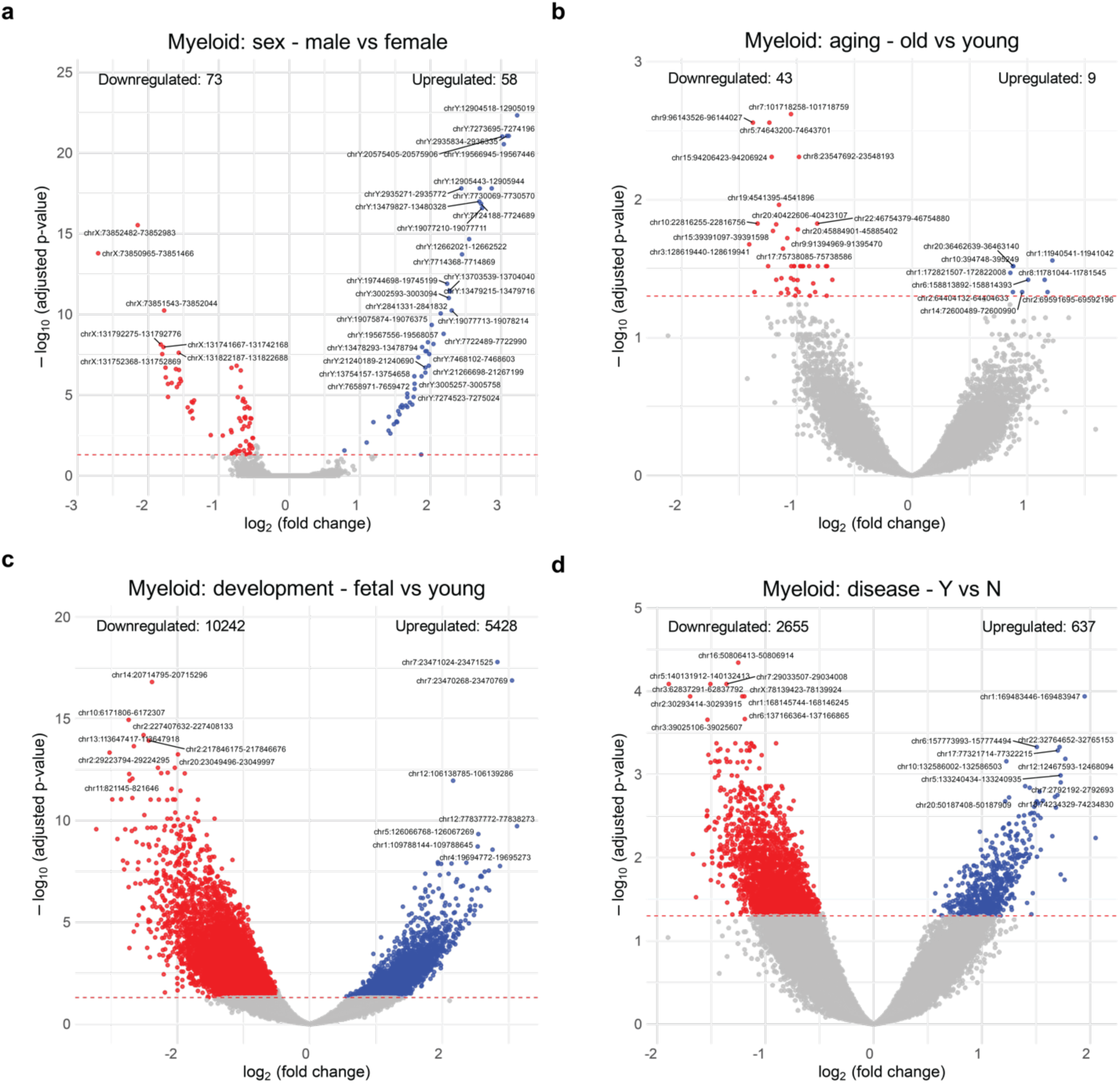
snATAC-seq Pericyte volcano plots: Using DESeq2, donor-pseudobulked snATAC-seq Pericyte volcano plots for the **(a)** sex, **(b)** aging, **(c)** development and **(d)** disease contrasts. Significance thresholds: |log2 fold change| > 0.5 and Benjamini-Hochberg adjusted p-value < 0.05.

**Supplemental Figure 46.**
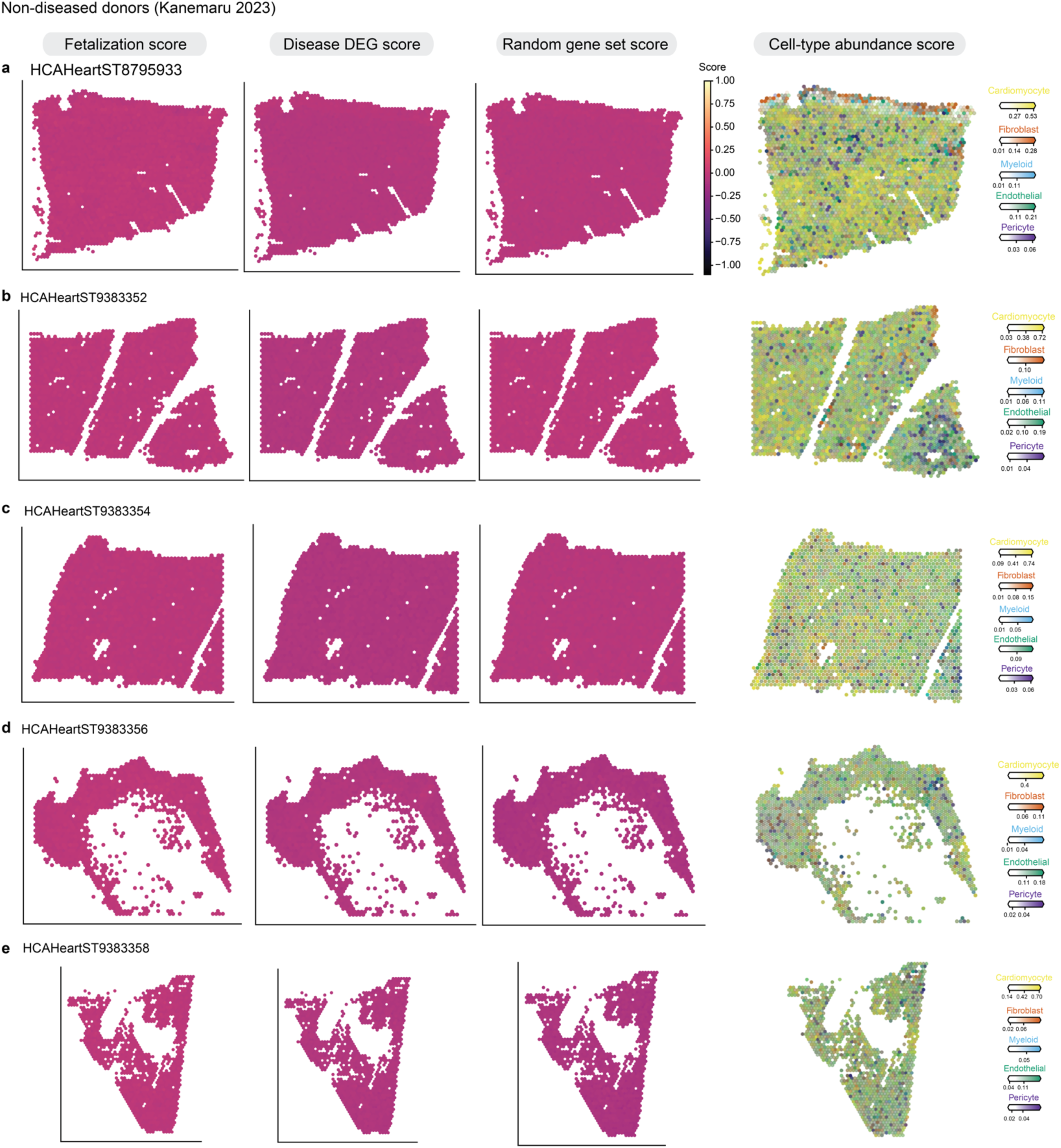
Kanemaru 2023 spatial non-diseased fetal reactivation: Gene activity scores were calculated using scanpy’s score_genes function. The gene scores for fetal reactivation genes (upregulated DEGs in both disease and fetal states), disease up DEGs, and a random set of genes of same size of fetal reactivation genes are shown for each donor. Scores are the weighted by the cell type proportions from cell2location. Cell type proportions for the 5 most common cell types from snRNA-seq are shown on right. All Kanemaru donors did not have diagnosed disease. **(a)** HCAHeartST8795933 **(b)** HCAHeartST9383352 **(c)** HCAHeartST9383354 **(d)** HCAHeartST9383356 **(e)** HCAHeartST9383358.

**Supplemental Figure 47.**
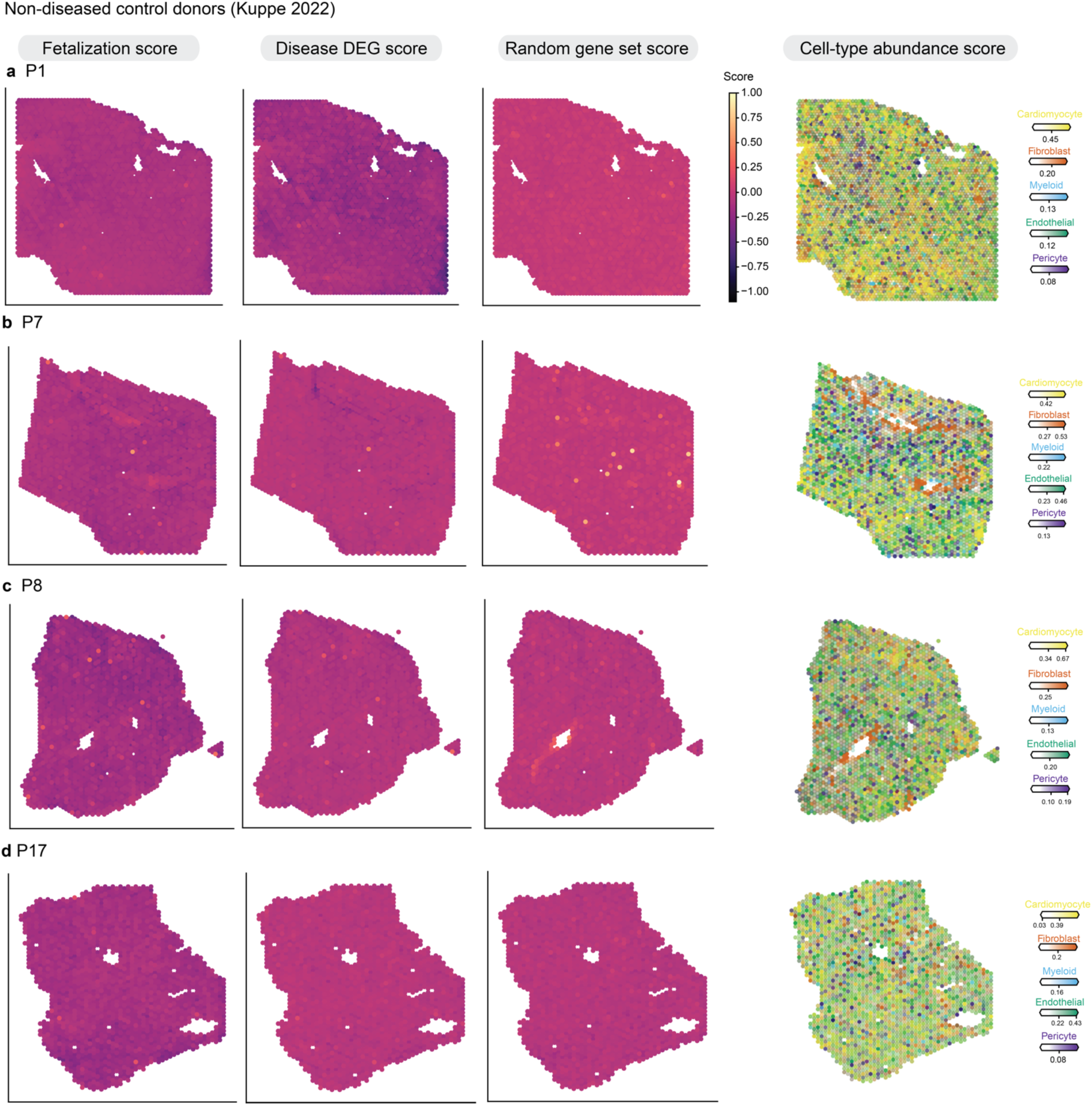
Kuppe 2022 spatial non-diseased fetal reactivation: Gene activity scores were calculated using scanpy’s score_genes function. The gene scores for fetal reactivation genes (upregulated DEGs in both disease and fetal states), disease up DEGs, and a random set of genes of same size of fetal reactivation genes are shown for each donor. Scores are the weighted by the cell type proportions from cell2location. Cell type proportions for the 5 most common cell types from snRNA-seq are shown on right. This figure includes spatial scores for non-diseased donors **(a)** P1 **(b)** P7 **(c)** P8 **(d)** P17.

**Supplemental Figure 48.**
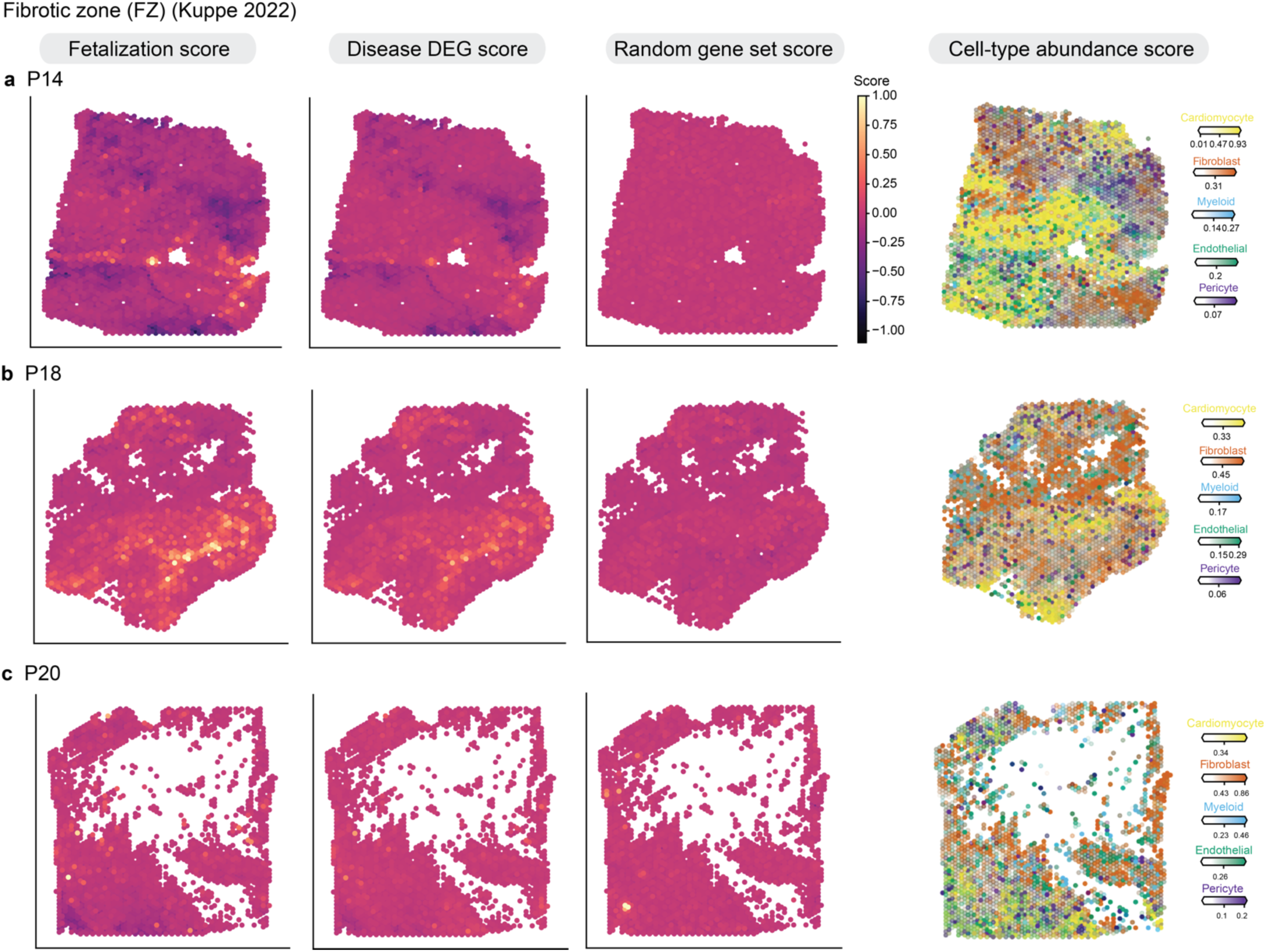
Kuppe 2022 spatial fibrotic zone fetal reactivation: Gene activity scores were calculated using scanpy’s score_genes function. The gene scores for fetal reactivation genes (upregulated DEGs in both disease and fetal states), disease up DEGs, and a random set of genes of same size of fetal reactivation genes are shown for each donor. Scores are the weighted by the cell type proportions from cell2location. Cell type proportions for the 5 most common cell types from snRNA-seq are shown on right. This figure includes spatial scores for the fibrotic zones of these donors **(a)** P14 **(b)** P18 **(c)** P20.

**Supplemental Figure 49.**
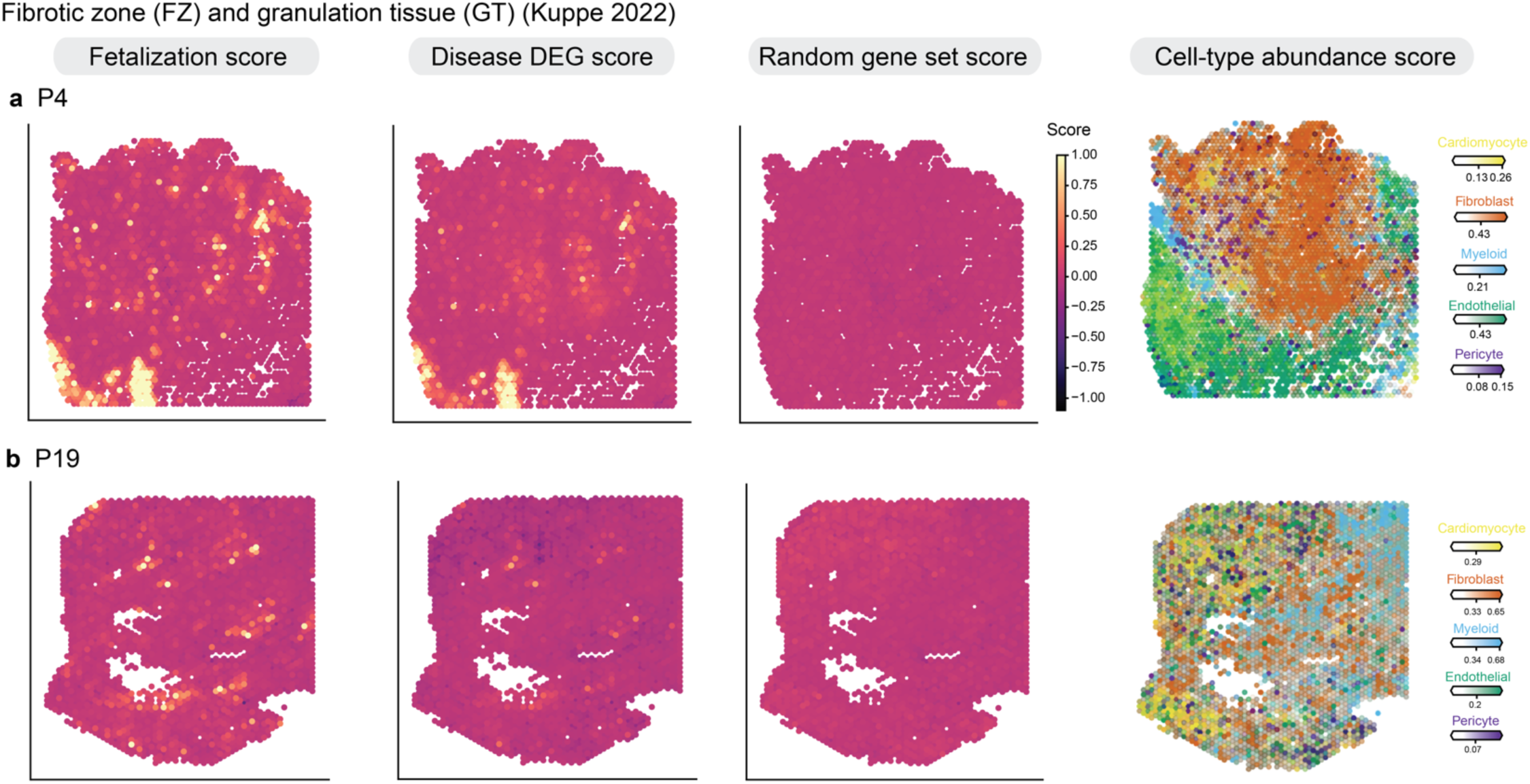
Kuppe 2022 spatial fibrotic zone + granulation tissue fetal reactivation: Gene activity scores were calculated using scanpy’s score_genes function. The gene scores for fetal reactivation genes (upregulated DEGs in both disease and fetal states), disease up DEGs, and a random set of genes of same size of fetal reactivation genes are shown for each donor. Scores are the weighted by the cell type proportions from cell2location. Cell type proportions for the 5 most common cell types from snRNA-seq are shown on right. This figure includes spatial scores for the fibrotic zone and granulation tissue of these donors **(a)** P4 **(b)** P19.

**Supplemental Figure 50.**
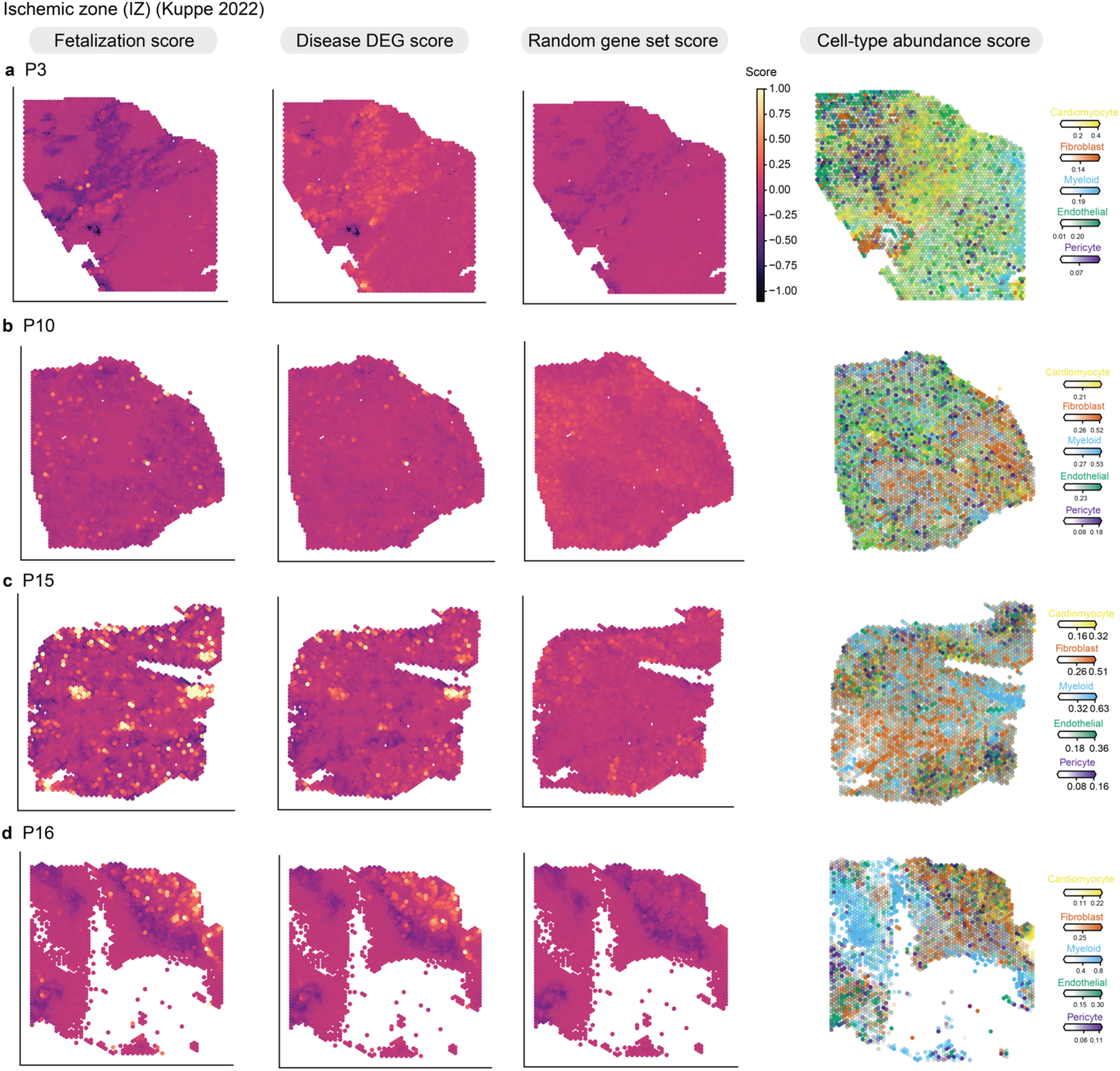
Kuppe 2022 spatial ischemic zone fetal reactivation: Gene activity scores were calculated using scanpy’s score_genes function. The gene scores for fetal reactivation genes (upregulated DEGs in both disease and fetal states), disease up DEGs, and a random set of genes of same size of fetal reactivation genes are shown for each donor. Scores are the weighted by the cell type proportions from cell2location. Cell type proportions for the 5 most common cell types from snRNA-seq are shown on right. This figure includes spatial scores for the ischemic zone and granulation tissue of these donors **(a)** P3 **(b)** P10 **(c)** P15 **(d)** P16.

**Supplemental Figure 51.**
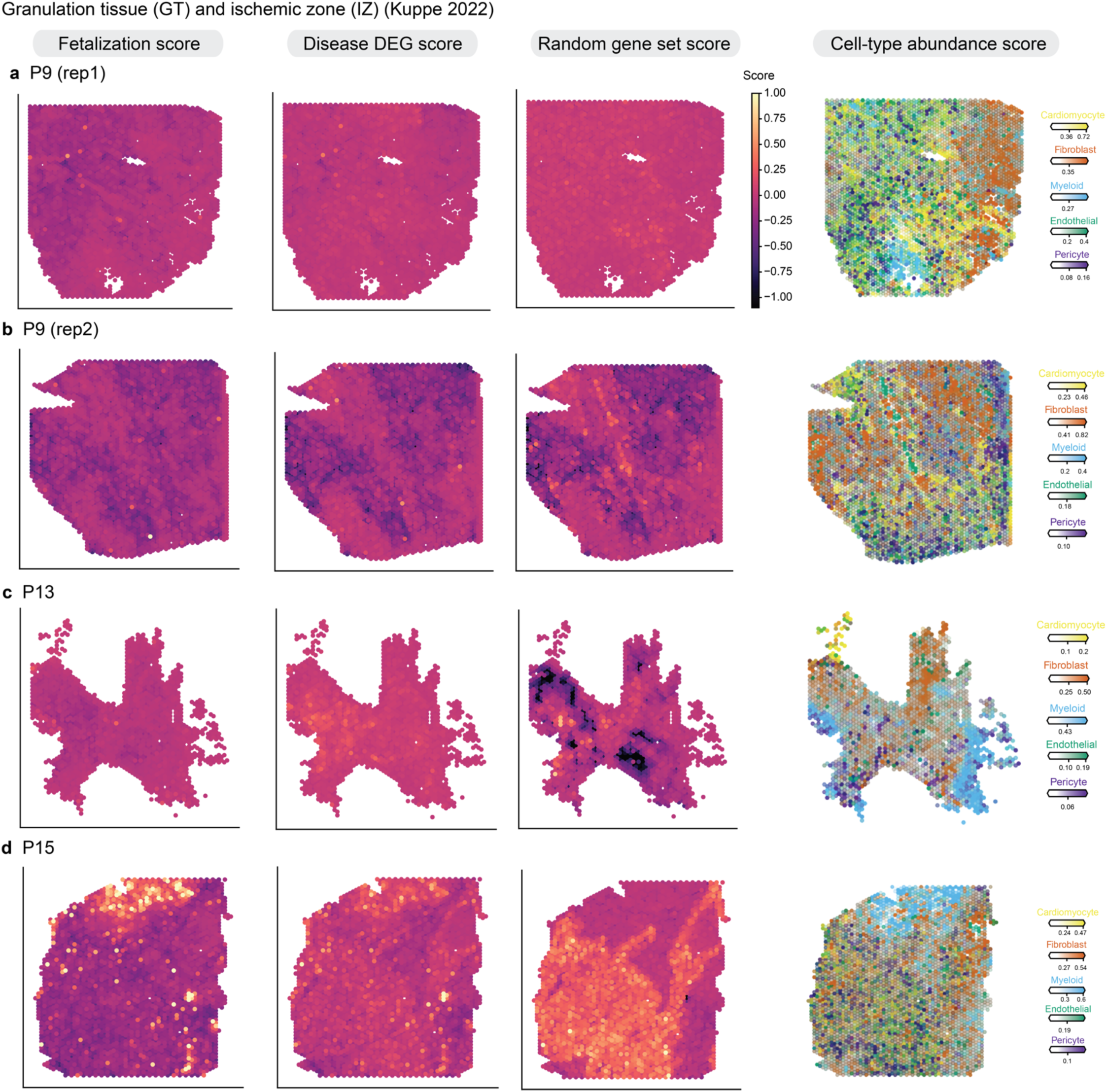
Kuppe 2022 spatial ischemic zone and granulation tissue fetal reactivation: Gene activity scores were calculated using scanpy’s score_genes function. The gene scores for fetal reactivation genes (upregulated DEGs in both disease and fetal states), disease up DEGs, and a random set of genes of same size of fetal reactivation genes are shown for each donor. Cell type proportions for the 5 most common cell types from snRNA-seq are shown on right. Scores are the weighted by the cell type proportions from cell2location. This figure includes spatial scores for the ischemic zone and granulation tissue of these donors **(a)** P9 **(b)** P9, second replicate **(c)** P13 **(d)** P15.

**Supplemental Figure 52.**
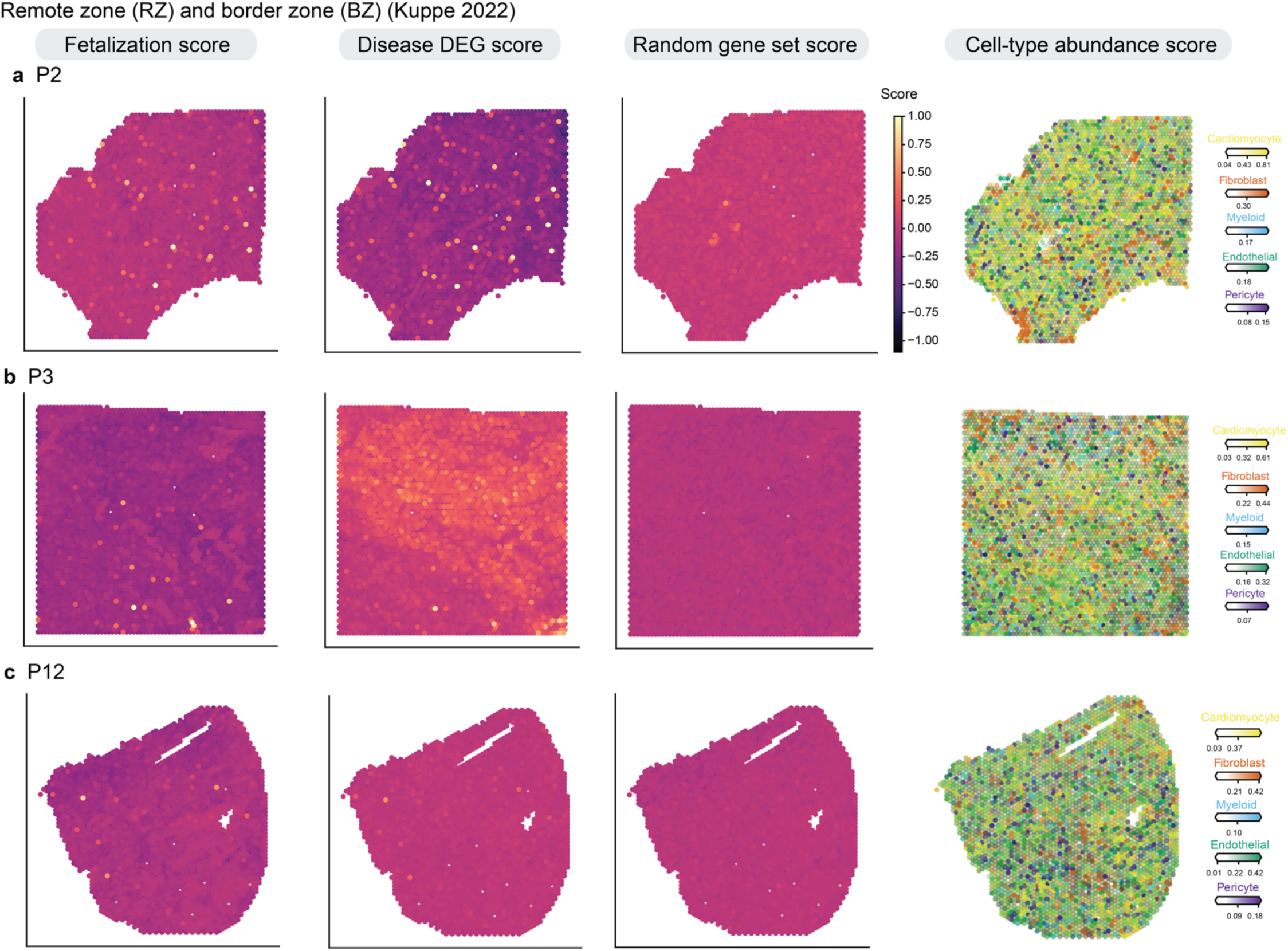
Kuppe 2022 spatial remote zone and border zone fetal reactivation: Gene activity scores were calculated using scanpy’s score_genes function. The gene scores for fetal reactivation genes (upregulated DEGs in both disease and fetal states), disease up DEGs, and a random set of genes of same size of fetal reactivation genes are shown for each donor. Scores are the weighted by the cell type proportions from cell2location. Cell type proportions for the 5 most common cell types from snRNA-seq are shown on right. This figure includes spatial scores for the remote zone and border zone of these donors **(a)** P2 (**b**) P3 **(c)** P12.

**Supplemental Figure 53.**
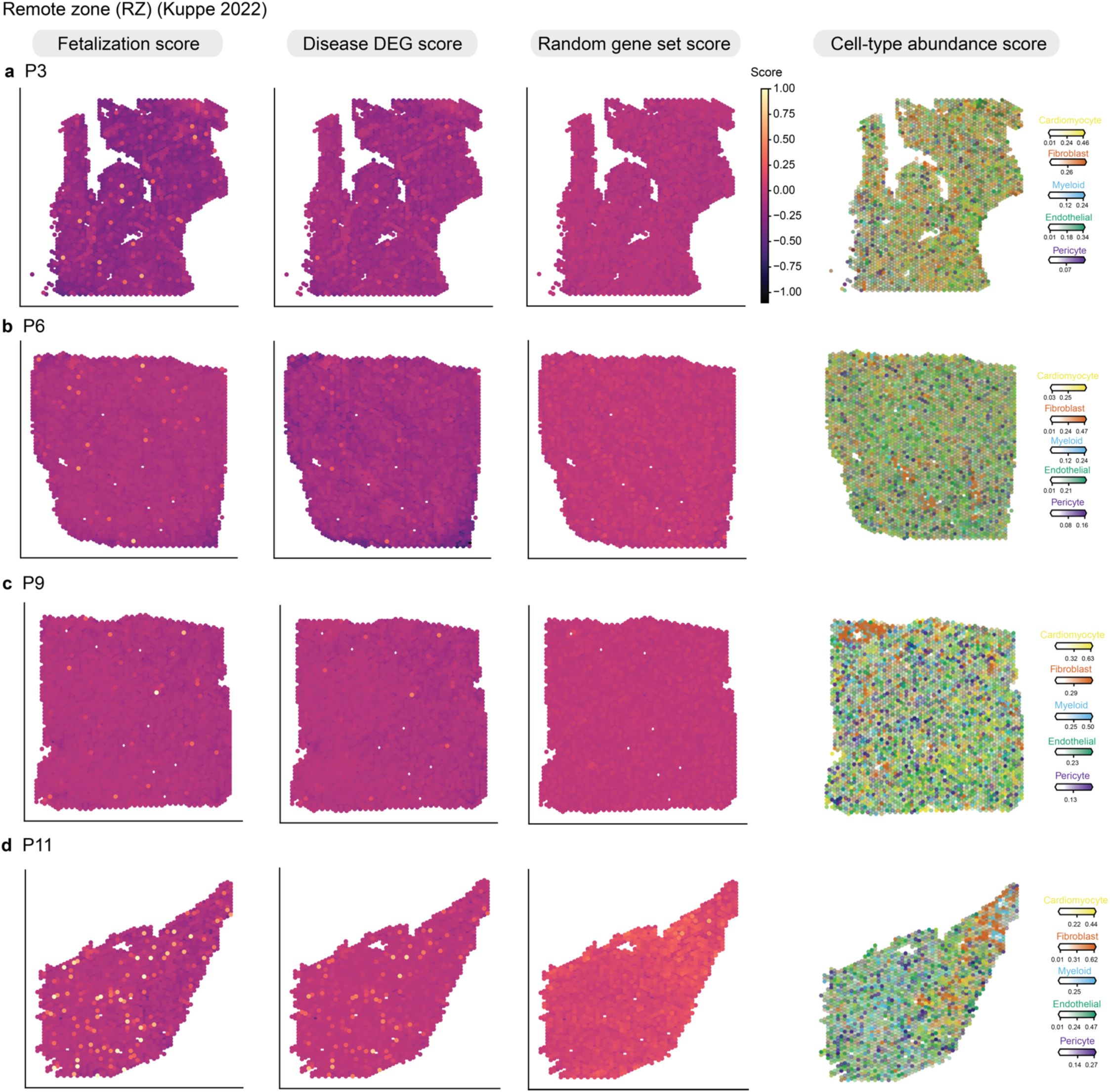
Kuppe 2022 spatial remote zone fetal reactivation: Gene activity scores were calculated using scanpy’s score_genes function. The gene scores for fetal reactivation genes (upregulated DEGs in both disease and fetal states), disease up DEGs, and a random set of genes of same size of fetal reactivation genes are shown for each donor. Scores are the weighted by the cell type proportions from cell2location. Cell type proportions for the 5 most common cell types from snRNA-seq are shown on right. This figure includes spatial scores for the remote zone of these donors **(a)** P3 **(b)** P6 **(c)** P19 **(d)** P11.

**Supplemental Figure 54.**
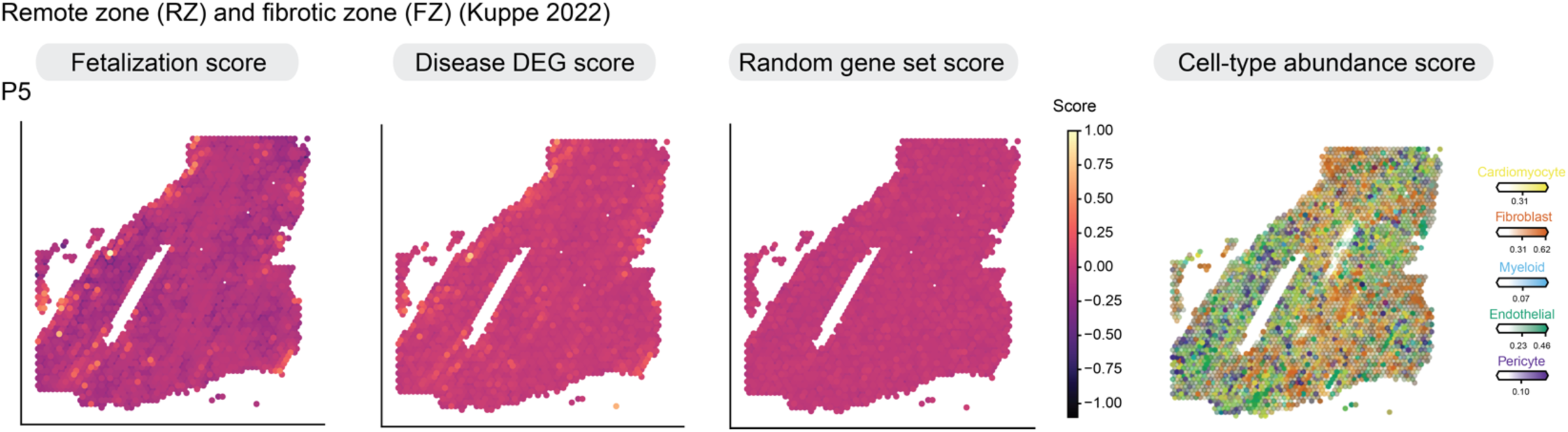
Kuppe 2022 spatial remote zone and fibrotic zone fetal reactivation: Gene activity scores were calculated using scanpy’s score_genes function. The gene scores for fetal reactivation genes (upregulated DEGs in both disease and fetal states), disease up DEGs, and a random set of genes of same size of fetal reactivation genes are shown for each donor. Scores are the weighted by the cell type proportions from cell2location. Cell type proportions for the 5 most common cell types from snRNA-seq are shown on right. This figure includes spatial scores for the remote zone and fibrotic zone of these donors P5.

**Supplemental Figure 55.**
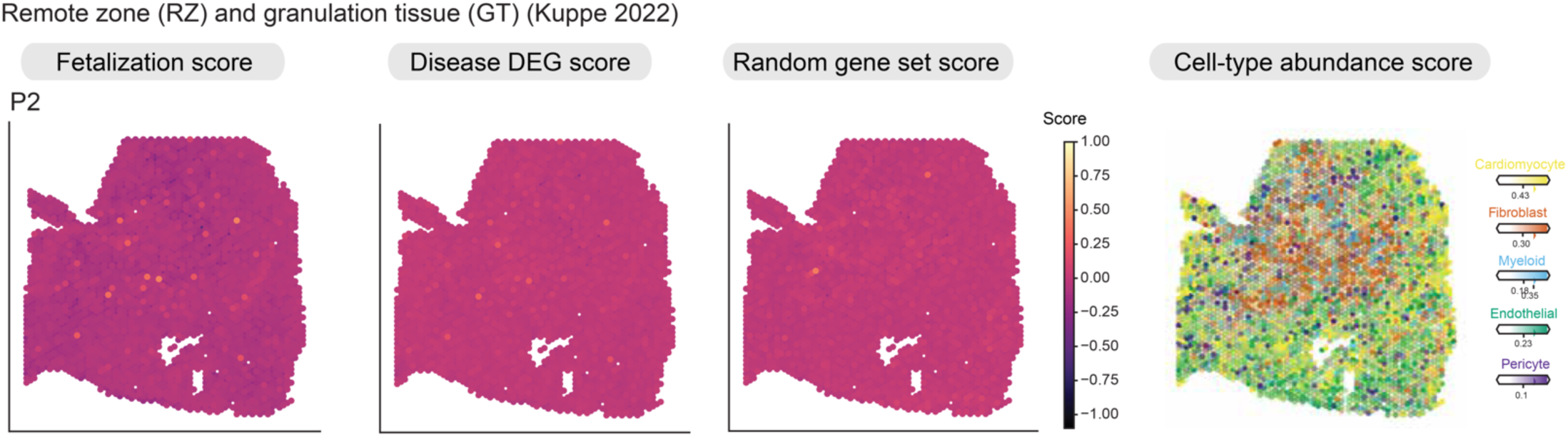
Kuppe 2022 spatial remote zone and granulation tissue fetal reactivation: Gene activity scores were calculated using scanpy’s score_genes function. The gene scores for fetal reactivation genes (upregulated DEGs in both disease and fetal states), disease up DEGs, and a random set of genes of same size of fetal reactivation genes are shown for each donor. Scores are the weighted by the cell type proportions from cell2location. Cell type proportions for the 5 most common cell types from snRNA-seq are shown on right. This figure includes spatial scores for the remote zone and granulation tissue of these donors P2.

