## Supplemental Table S1-41 for "Integrated single-cell and spatial multiomic analysis reveals widespread reactivation of developmental programs in diseased human hearts": List_of_supplemental_tables.docx

- S1: Sources of snRNA-seq and snATAC-seq data
- S2: snRNA-seq donor level metadata
- S3: snATAC-seq donor level metadata
- S4: Number of differentially expressed genes (DEGs) per cell type and contrast
- S5: Adipocyte DEGs
- S6: Cardiomyocyte DEGs
- S7: Endocardial DEGs
- S8: Endothelial DEGs
- S9: Epicardial DEGs
- S10: Fibroblast DEGs
- S11: LEC DEGs
- S12: Lymphoid DEGs
- S13: Mast DEGs
- S14: Myeloid DEGs
- S15: Neuronal DEGs
- S16: Pericyte DEGs
- S17: vSMC DEGs
- S18: Adipocyte GSEA results
- S19: Cardiomyocyte GSEA results
- S20: Endocardial GSEA results
- S21: Endothelial GSEA results
- S22: Epicardial GSEA results
- S23: Fibroblast GSEA results
- S24: LEC GSEA results
- S25: Lymphoid GSEA results
- S26: Mast GSEA results
- S27: Myeloid GSEA results
- S28: Neuronal GSEA results
- S29: Pericyte GSEA results
- S30: vSMC GSEA results
- S31: Cell type proportion results by sex
- S32: Cell type proportion results by age group
- S33: Cell type proportion results by developmental stage
- S34: Cell type proportion results by disease status
- S35: Number of differentially accessible regions (DARs) per cell type and contrast
- S36: Cardiomyocyte DARs
- S37: Endothelial DARs
- S38: Fibroblast DARs
- S39: Lymphoid DARs
- S40: Myeloid DARs
- S41: Pericyte DARs
